# Widely Distributed Biosynthetic Cassette Is Responsible for Diverse Plant Side-Chain-Cross-Linked Cyclopeptides

**DOI:** 10.1101/2022.09.15.507631

**Authors:** Stella T. Lima, Brigitte G. Ampolini, Ethan B. Underwood, Tyler N. Graf, Cody E. Earp, Imani C. Khedi, Jonathan R. Chekan

## Abstract

Cyclopeptide alkaloids are an abundant class of plant cyclopeptides with over 200 analogs described and bioactivities ranging from analgesic to antiviral. While these natural products have been known for decades, their biosynthetic basis remains unclear. Using a transcriptome-mining approach, we link the cyclopeptide alkaloids from *Ceanothus americanus* to dedicated RiPP precursor peptides and identify new, widely distributed split BURP-domain containing gene clusters. Precursor peptides from these biosynthetic cassettes directly map to both cyclopeptide alkaloids from *Ziziphus jujuba* and the structurally distinct hibispeptins from *Hibiscus syriacus*. Guided by our bioinformatic analysis, we identify and isolate new cyclopeptides from *Coffea arabica*, which we named arabipeptins. These results expand our understanding of the biosynthetic pathways responsible for diverse plant side chain cross-linked cyclopeptides and suggest the presence of previously unknown natural products or protein post-translational modifications that are widely distributed in eudicots.

---

Plants are prolific producers of cyclic, peptide derived natural products including cyclotides^[1]^, orbitides^[2]^, and cyclopeptide alkaloids^[2,3]^. Detailed biosynthetic studies have demonstrated that many of these compounds are ribosomally synthesized and post-translationally modified peptide (RiPP) natural products^[4–8]^. RiPPs do not have a family defining biosynthetic step or feature. Instead, they are classified by their route of biosynthesis wherein a ribosomally produced precursor peptide is modified by at least one tailoring enzyme (Figure 1A)^[9]^. The majority of RiPP precursor peptides are composed of a conserved leader sequence that is responsible for recognition by the tailoring enzymes and a core peptide that is modified. In the final biosynthetic step, a peptidase will typically release the core from the precursor peptide to generate the mature natural product. Because RiPPs are a large family of natural products with over 40 distinct classes, RiPP pathways have also been shown to deviate from this general route and include precursor peptides that house an array of multiple core sequences or are fused to autocatalytic tailoring enzymes (Figure 1A).^[5–7,10]^

**Figure 1.**
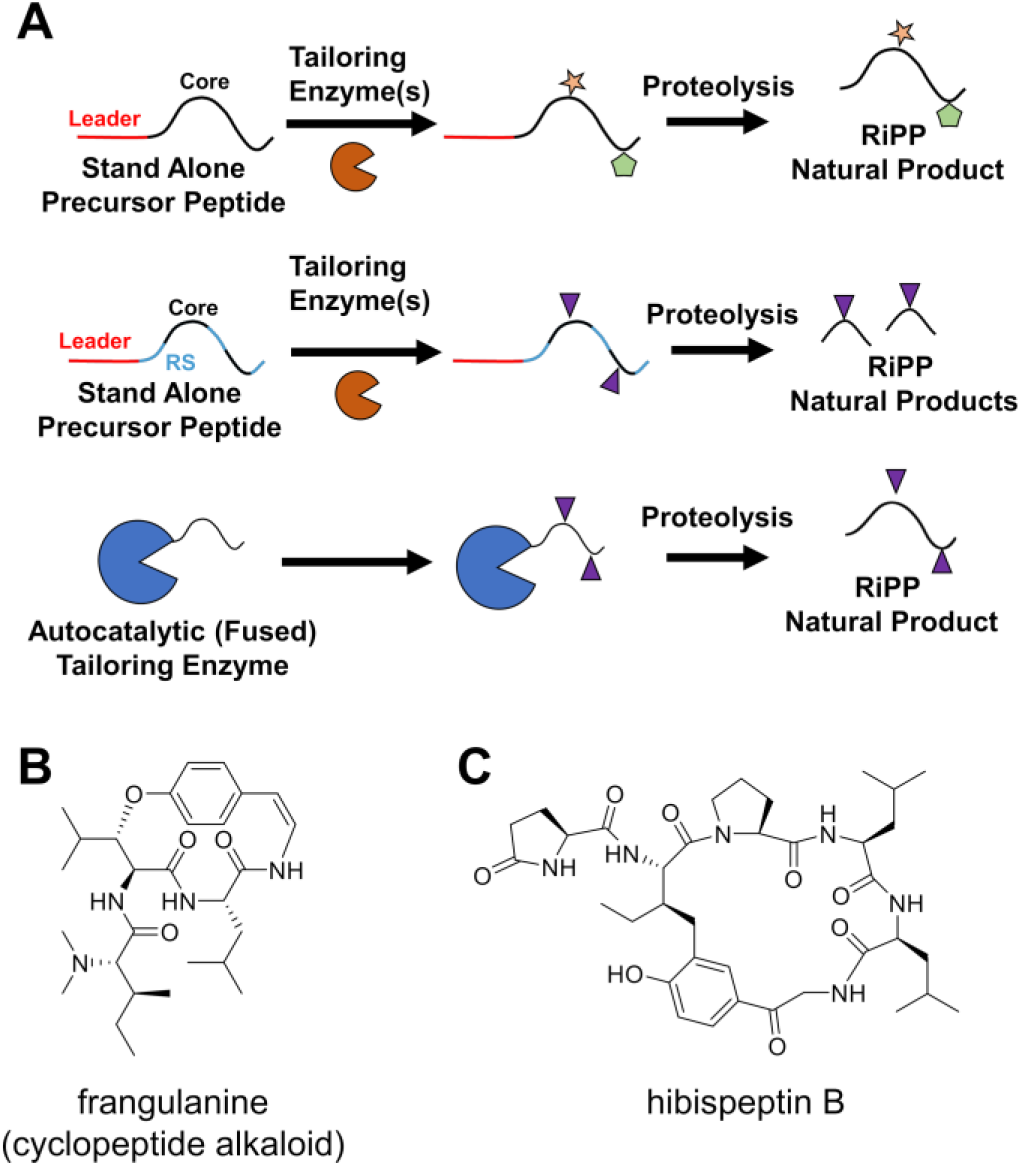
(A) Overview of RiPP biosynthetic routes. Catalyzed amino acid modifications are indicated with colored shapes. Recognition sequence, RS. (B and C) Plant side chain cross-linked cyclopeptides.

Cyclopeptide alkaloids are among the most abundant classes of plant-produced peptides with over 200 members (Figure 1B)^[2,3]^. They are four or five amino acids in size and are proposed to be formed from an ether linkage between the phenolic oxygen of tyrosine and the β-carbon of a nearby amino acid. The tyrosine also undergoes a decarboxylation and desaturation to generate the characteristic styrylamine moiety. Additional modifications including methylation and hydroxylation are also common. Cyclopeptide alkaloids have a wide range of bioactivities including antiviral^[11]^, sedative^[12]^, and analgesic^[13]^. Even though they were first identified over 50 years ago, little is known about their biosynthesis. Structurally, cyclopeptide alkaloids are related to diverse plant side chain cross-linked cyclopeptides that include lyciumins^[5]^, selanines^[6]^, moroidins^[7]^, and hibispeptins^[14,15]^ (Figure 1C). Recent work demonstrated that many of these cyclic peptides are derived from autocatalytic BURP-domains that install the characteristic amino acid side chain macrocyclizations^[5–7,16,17]^. In these systems, the core peptides are fused to the copper-dependent BURP-domains. While the structural similarities between these plant cyclopeptides hint at a conserved biosynthetic route for cyclopeptide alkaloids, no definitive link has yet been made.

To discover the biosynthetic basis for cyclopeptide alkaloids, we examined *Ceanothus americanus* L. (Rhamnaceae), which is commonly known as New Jersey tea. This plant is a well-known producer of cyclopeptide alkaloids with at least ten different molecules isolated, such as frangulanine (Figure 1B).^[18–20]^ Study of *C. americanus* would not only give insight into the biosynthesis of bioactive cyclopeptide alkaloids, but would also explain how a single plant is able to make so many diverse analogs.

In order to link the known cyclopeptide alkaloids to their precursor peptides, we generated transcriptomic data sets from the leaf, stem, and root of a single *C. americanus* individual. We first searched our translated transcriptomic data sets for BURP-domain containing transcripts using an available Hidden Markov Model (HMM) for the protein family (PF03181) to identify possible autocatalytic precursors (Supplementary Table). Surprisingly, there were no clear correlations between the amino acid sequences in the BURP-domains and the predicted core sequences of the cyclopeptide alkaloids that we confirmed to be present in the roots of our *C. americanus* samples by LC-MS (Figures S1-S8).

As the observed cyclopeptide alkaloids did not appear to be derived from autocatalytic BURP-domains, we queried the entire translated transcriptome for all four-amino acid sequences that could lead to the observed natural products (Figure S1). Clustering these candidates by sequence identity revealed related transcripts that corresponded to the observed cyclopeptide alkaloids (Figure 2A and S9). Each of these precursors contained an N-terminal signal sequence and a conserved leader peptide-like region (Figure S10). This is followed by a repeating recognition sequence and core structure that has previously been observed in cyanobactins^[10]^ and numerous cyclic peptides from plants^[5,6]^. The core sequences themselves are typically demarcated by a conserved N-terminal Asn and a C-terminal His (Figure S9 and S11). To confirm our hypothesis that these precursor peptides are directly linked to the observed cyclopeptide alkaloids, we identified E-L-W-Y and F-F-F-Y cores for which no known cyclopeptide alkaloids correspond (Figure S9). We searched the feature list obtained from LC-MS analysis of a *C. americanus* root extract for ions consistent with these new cyclopeptides and found features with the expected *m/z* and MS/MS fragmentation patterns (Figure S12 and S13).

**Figure 2.**
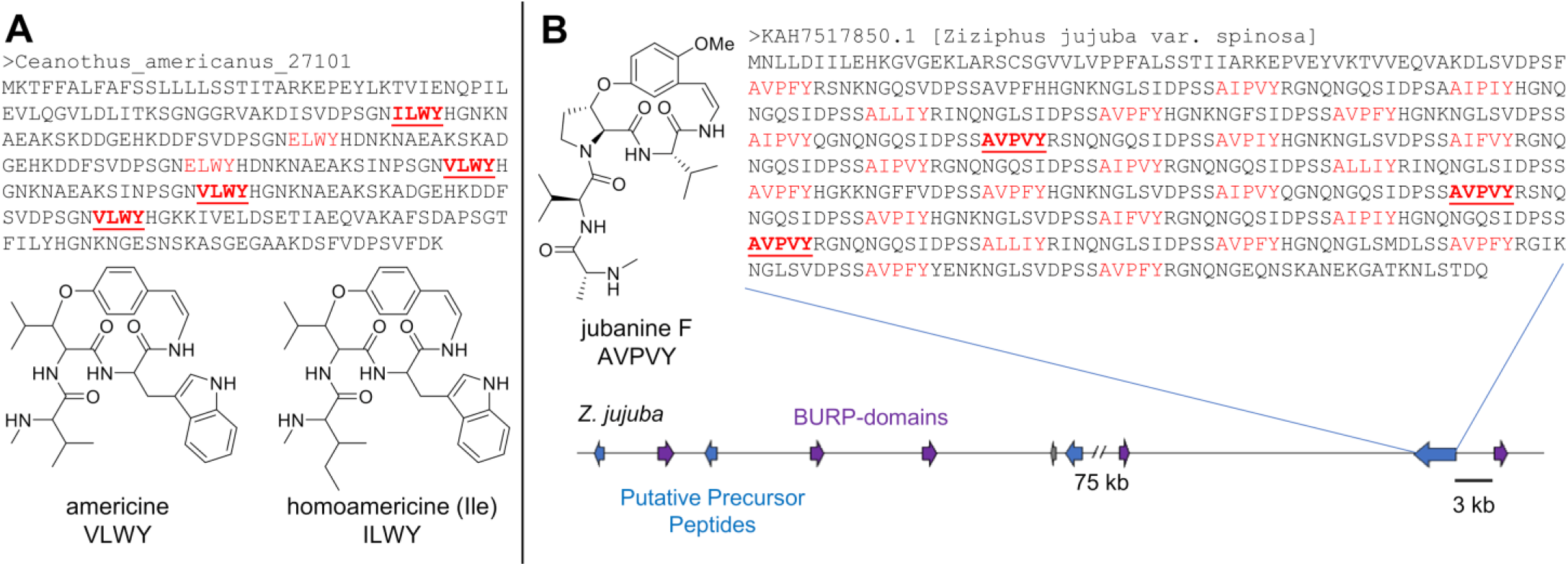
Stand alone precursor peptides from *C. americanus* (A) and *Z. jujuba* (B) contain multiple cores that match directly to known cyclopeptide alkaloids (red, bold). (B) Precursor peptides are co-clustered with BURP-domains in the genome of *Z. jujuba*. Predicted amino acid sequence is shown below the molecule name.

After linking *C. americanus* cyclopeptide alkaloids to dedicated precursor peptides, we sought to evaluate if these stand alone precursor peptides correspond to other known plant side chain crosslinked cyclopeptides. We used BLAST to query the genome of the prolific cyclopeptide alkaloid producer *Ziziphus jujuba* for homologs of the *C. americanus* precursor peptides. The resulting hits were compared to the anticipated core sequences of the known *Z. jujuba* cyclopeptide alkaloids and 15 molecules could be directly linked to a core sequence including jubanines F, G, H, I, J, and nummularine B (Figures 2B and S14).^[11]^ As with *C. americanus*, most precursor peptides contained an N-terminal signal sequence followed by a conserved leader-like region and repeating recognition sequence + core architecture (Figures 2B, S11, and S15).

Examination of the *Z. jujuba* genome revealed that 30 of the 35 annotated precursor peptides were co-clustered with a BURP-domain containing protein. Moreover, there were genomic loci containing multiple precursor peptides and BURP-domains arrayed together (Figure 2B). When aligned to the autocatalytic BURP-domains, the stand alone BURP-domains are shorter and lack the N-terminal extension that contains the core peptide(s) (Figure S16). The proximity of the BURP-domains to the precursors in the genomes along with chemistry consistent with a post-translational modification known to be catalyzed by BURP-domains indicates the presence of a new split BURP-domain biosynthetic system that parallels the fused autocatalytic BURP-domains.

To explore the distribution of this split BURP-domain system, we built a custom Hidden Markov Model (HMM) using PSI-BLAST results from the *C. americanus* precursor peptide (Supplementary Table) and analyzed all the Viridiplantae sequences available from the NCBI identical protein group database. Unexpectedly, a significant portion of the resulting sequences were fused to a BURP-domain. The detection of BURP-domain proteins is likely due to the presence of a highly conserved Ax6YWx7PMP motif that follows the signal peptide in many autocatalytic BURP-domains and stand alone precursor peptides (Figure S17). We used an existing BURP-domain HMM to differentiate between these fused and split biosynthetic systems. This ultimately allowed for the identification of 1,423 stand alone precursor peptides and 1,099 fused BURP-domain systems. We analyzed the genomic context of the stand alone precursor peptides to determine how often they co-cluster with BURP-domains. At least 58% of the precursor peptides were found within ten annotated genes of a BURP-domain (Supplementary Table). This overall high level of co-occurrence suggests that the precursor peptide and BURP-domain pairs form a conserved biosynthetic cassette.

In addition to co-occurrence, precursor peptides appear to be transcriptionally co-regulated with a split BURP-domain. Published work from *Z. jujuba* showed that sanjoinine A content was most closely associated with transcript levels of a genomic locus containing both a split BURP-domain and methyltransferase.^[21]^ We also analyzed the ATTED-II database^[22]^ for genes co-expressed with putative precursor peptides from *Arabidopsis thaliana, Vitis vinifera, Populus trichocarpa, Glycine max, Medicago truncatula*, and *Solanum lycopersicum*. In each case, split BURP-domain containing proteins are amongst the three most co-regulated genes (Supplementary Table).

The precursor peptides for the split systems were further explored by generating sequence similarity networks (SSNs)^[23]^ (Figures 3 and S18). Putative core sequences were manually annotated using the observed recognition sequence elements from *C. americanus* and *Z. jujuba* (Figure S11 and Supplementary Table). Throughout the entire sequence similarity network, a C-terminal tyrosine is a highly conserved feature of the core. The largest cluster is characterized by putative four-membered core sequences with a Ser or Thr in the second position and an almost completely conserved Tyr (Figure S11). Even though examination of the genomic context reveals a 59% co-localization rate with BURP-domains (Supplementary Table), no small molecules have been isolated from plants that match this motif to our knowledge^[2]^. The vast majority of these precursor peptides belong to the organ specific protein family (PF10950).^[24]^ Members of this family contain an N-terminal signal peptide followed by a series of features exhibiting RiPP precursor peptide characteristics, including a conserved leader peptide-like motif and alternating hypothetical recognition sequence and core peptide repeats containing a completely conserved tyrosine.^[25]^ While the exact function of organ specific proteins remains unclear, they are hypothesized to play a role in abiotic stress response, root development, nodule formation, and establishment of mycorrhizal interactions.^[24–27]^ Whether these proteins are processed to secondary metabolites or simply post-translationally modified requires further study.

**Figure 3.**
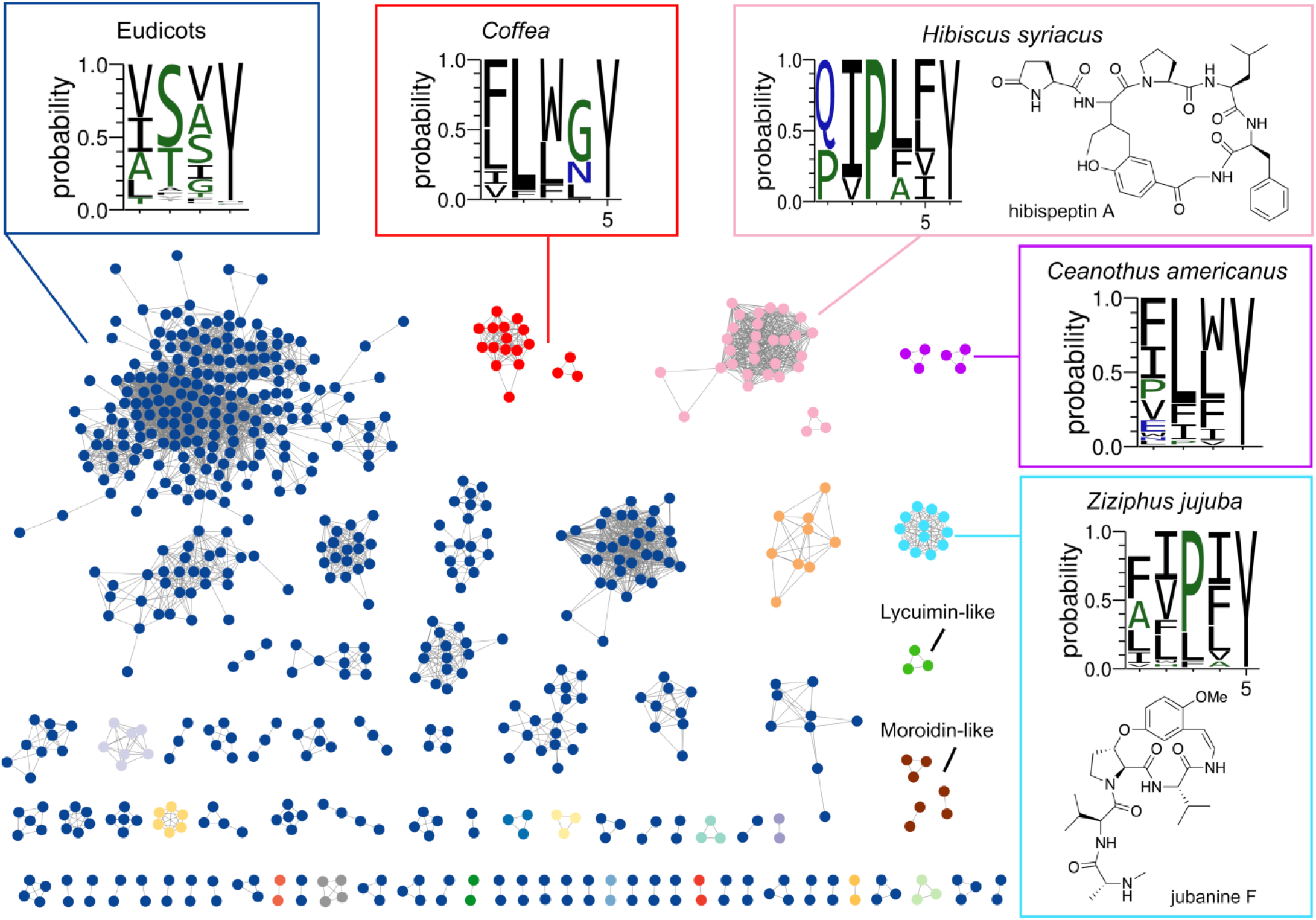
Sequence similarity network of stand alone precursor peptides using an alignment score of 70. Singlets are shown in Figure S18. Precursor peptides with cores containing serine or threonine in the second position are represented in navy blue. Clusters from *H. syriacus* (pink), *Z. jujuba* (light blue), *Coffea* (red), and *C. americanus* (purple) are indicated along with those with lycuimin-like (green) and moroidin-like (brown) core sequences. Amino acid probabilities of the core sequences for these clusters are depicted.

In contrast to the prevalence of Ser/Thr containing cores without a clear link to a known natural product or peptide modification, other clusters could be mapped to known plant side chain cross-linked cyclopeptide natural products. For example, the cyclopeptide alkaloid precursor peptides from *C. americanus* and *Z. jujuba* formed discrete clusters (Figure 3).

In addition to clusters from the known cyclopeptide alkaloid producers, a large cluster of putative precursor peptides from *Hibiscus syriacus* was also present in the SSN (Figure 3). Precursor peptides from this cluster were rich in the core sequences Q-I-P-L-F-Y and Q-I-P-L-L-Y, which match the known side chain cross-linked cyclopeptides hibispeptin A and B, respectively (Figure 1C, 3, and S19).^[14,15]^ As observed with the cyclopeptide alkaloids, BURP-domains were also found clustered with these precursor peptides (Supplementary Table). In addition to the known hibispeptins, the presence of uncharacterized core sequences suggested that undiscovered analogs may exist (Supplementary Table). Therefore, we analyzed *H. syriacus* root extracts by LC-MS. As expected, detected ions matching the known molecules hibispeptin A and B were observed and supported by MS/MS fragmentation (Figures S20 and S21). To discover unreported hispeptin analogs that could be explained by the precursor peptides, we generated a GNPS molecular network^[28]^ (Figure S22). A six-node cluster was found that contained both hibispeptin A and B. In addition to suspected derivatives of hibispeptin A and B, a node with an *m/z* of 669.3638 was identified. This *m/z* matches the predicted value (669.3606 *m/z*, Δ 4.77 ppm) for the [M+H]^+^ ion of the Q-V-P-L-V-Y core peptide which is found in the same gene as the hibispeptin A and B cores (Figure S19). MS/MS analysis supported the presence and structure of this new hibispeptin analog (Figure S23).

Along with putatively linking known molecules to their precursor peptides, we also sought to use our SSN to discover new side chain cross-linked cyclopeptides from a previously unknown producer. Multiple members of the *Coffea* genus, including *Coffea arabica*, had potential precursor peptides with clear tyrosine-containing sequence repeats (Figures 3, S11, and S24). Notably, over 50% of these *Coffea* precursor peptide sequences co-localized with BURP-domain proteins (Figures 4A, S25, and Supplementary Table). Therefore, we analyzed methanol extracts of *C. arabica* by LC-MS and generated a combined GNPS molecular network with *C. americanus* to identify features related to cyclopeptide alkaloids (Figure 4B). Twelve nodes from *C. arabica* clustered together and we could clearly map five of these mass features to core sequences (Figure S24 and S26-S30). Based on the MS/MS fragmentation and analogy to known cyclopeptides, these new secondary metabolites from *C. arabica* were proposed to be four or five amino acids in size and cyclized through an ether bond between the conserved C-terminal Tyr and the β-carbon of Ile/Leu. No decarboxylation was observed at the C-terminal Tyr, similar to cyclo-[VPIFY] from *G. max*^[6]^. We confirmed the predicted structure for the 711.3503 *m/z* ion by using spectroscopic analysis. Key HMBC and NOESY correlations supported the C-O crosslink between the β-carbon of leucine and the phenolic oxygen of tyrosine (Figure 4C). We named this new side chain cross-linked cyclopeptide arabipeptin A.

**Figure 4.**
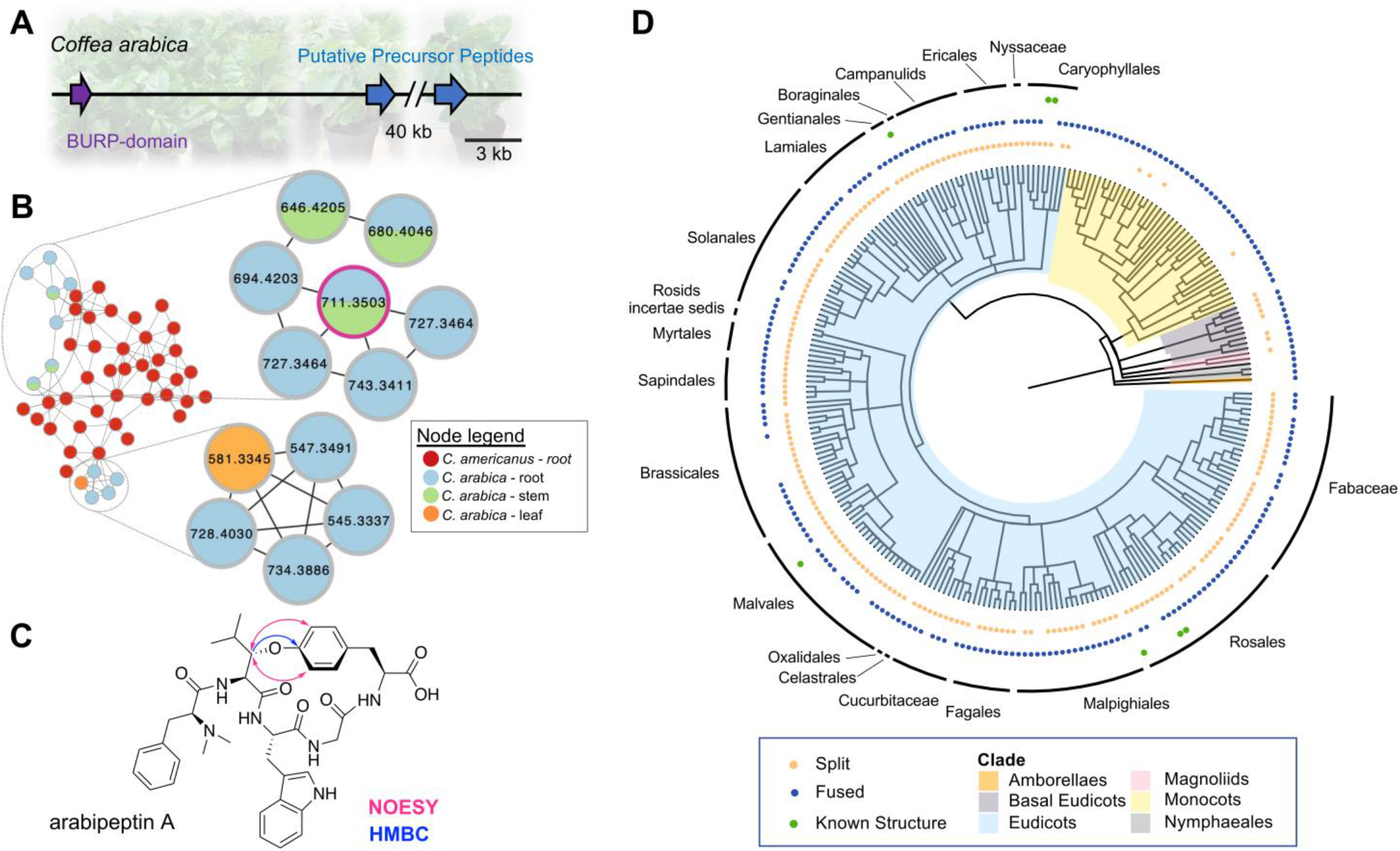
(A) Representative gene cluster from *C. arabica*. (B) Global Natural Product Social (GNPS) molecular network of metabolomic datasets from *C. arabica* and *C. americanus*. Indicated nodes were re-analyzed to generate new clusters. Nodes represent parent masses. Pie charts represent spectra of defined plant tissues. (C) Structure of characterized arabipeptin A. Pink and blue arrows represent key NOESY and HMBC correlations, respectively. (D) Cladogram of Viridiplantae. Blue and gold dots correspond to the presence of fused BURP-domains and stand alone precursor peptides, respectively. As every genome is not present or fully annotated, the lack of a dot does not suggest an absence in the organism. Green dots indicate the presence of known structures derived from a stand alone precursor peptide. Clades are highlighted and orders for the core eudicots are labeled adjacent to the corresponding black bar.

Even though *C. americanus*, *C. arabica*, and *H. syriacus* all contain these split BURP-domain pathways, they are not closely related by phylogeny. To better understand the prevalence and distribution of the split BURP-domain systems in Viridiplantae, a cladogram was constructed from all species identified as having a fused or split BURP-domain system by our custom HMM (Figure 4D and S31). This analysis showed that both the split precursor peptides and fused BURP-domains are nearly ubiquitous among eudicots. We also mapped natural products with known structures that can be attributed to stand alone precursor peptides onto the cladogram. These molecules were distributed across multiple orders including rosales (cyclopeptide alkaloids), malvales (hibispeptins), and gentianales (arabipeptins).

In summary, we linked cyclopeptide alkaloids from *C. americanus* to their precursor peptides and related these results to additional cyclopeptide alkaloids from *Z. jujuba*. Unexpectedly, the precursor peptides were not found fused to BURP-domains as has been the case for other plant side chain cross-linked cyclopeptides.^[5–7]^ Instead the cyclopeptide alkaloids appear to be the first example of a split BURP-domain biosynthetic system. A similar split and fused enzyme/precursor system was recently shown to exist in the borosin family of RiPPs^[29,30]^

In addition to cyclopeptide alkaloids, we found that the known molecules hibispeptin A and B are formed from stand alone precursor peptides. Our isolation of arabipeptin A from the roots of *C. arabica* validates that the core peptide sequences can indeed be used as a guide for the discovery of new plant metabolites. By identifying the precursor peptides and their co-localization of BURP-domains, our results lay the foundation for future studies to completely elucidate and re-engineer the biosynthesis of cyclopeptide alkaloids and hibispeptins.

## Acknowledgements

We acknowledge Dr. Daniel Todd for the collection and interpretation of mass spectrometry data. This work was supported by the University of North Carolina at Greensboro (research start-up funds and Faculty First Grant to J.R.C. and Undergraduate Research and Creativity Award to I.C.K.) and American Society of Pharmacognosy (Research Starter Grant to J.R.C.).

