## Supplementary Data for "Widely Distributed Biosynthetic Cassette Is Responsible for Diverse Plant Side-Chain-Cross-Linked Cyclopeptides"

### Table of Contents

1. **Data Deposition**

RNA Seq……………………………………………………………………… S3
Plant Vouchers …………………………………………………………… S3
NMR ………………………………………………………………………… S3
Mass Spectra ……………………………………………………………… S3

2. **Methods**

Metabolomics ……………………………………………………………… S3
RNA Extraction and Sequencing ………………………………………… S4
Transcriptome Assembly and Analysis …………………………………… S4
Hidden Markov Model & Cladogram ……………………………………… S5
Sequence Similarity Network ……………………………………………… S6
Purification of arabipeptin A ……………………………………………… S6
Structure elucidation of arabipeptin A …………………………………… S6
Marfey's reaction - arabipeptin A ………………………………………… S7

3. **Supplementary Figures**

Figure S1. Cyclopeptide alkaloids from *C. americanus* ………………… S9
Figure S2. MS/MS spectrum of ceanothine C or ceanothine D ………… S10
Figure S3. MS/MS spectrum of adouetine X or frangulanine …………… S11
Figure S4. MS/MS spectrum of ceanothine B …………………………… S12
Figure S5. MS/MS spectrum of ceanothine A …………………………… S13
Figure S6. MS/MS spectrum of americine ……………………………… S14

Figure S7. MS/MS spectrum of homoamericine ...……………………………….…..…… S15
Figure S8. MS/MS spectrum of adouetine Y and ceanothine E ………………………..…. S16
Figure S9. Core sequences from the precursor peptides from *C. americanus* ……...…. S19
Figure S10. Putative precursor peptides from *C. americanus* ……………….……….. S20
Figure S11. Weblogos for core sequences ……….……………..………………..……. S21
Figure S12. MS/MS spectrum of CAM554 ..……………………………………….... S22
Figure S13. MS/MS spectrum of CAM603 …...…………………………………...…. S23
Figure S14. Putative precursor peptides from *Ziziphus jujuba* ……………..………… S28
Figure S15. Putative precursor peptide alignment from *Ziziphus jujuba* ..…....……..…. S29
Figure S16. Amino acid sequence alignment of split/fused BURP-domains …....……. S30
Figure S17. Precursor peptide and fused BURP-domain sequence conservation …..... S31
Figure S18. SSN2 generated using the singlets from SSN1 ……..………..………..…. S32
Figure S19. Putative precursor peptide sequences from *Hibiscus syriacus* …..……… S33
Figure S20. MS/MS spectrum of hibispeptin A ……………..…..…………………… S34
Figure S21. MS/MS spectrum of hibispeptin B ……..……………..…………….…… S35
Figure S22. *H. syriacus* cyclopeptide-containing GNPScluster………………..………… S36
Figure S23. MS/MS spectrum of HIS669 ……………………………………..……… S37
Figure S24. Putative precursor peptides from the *C. arabica* ..……..…………...………… S39
Figure S25. Genomic organization of the arabipeptin precursor peptides ……...…… S40
Figure S26. MS/MS spectrum of arabipeptin A …………………….………….……… S41
Figure S27. MS/MS spectrum of CAR547  …..………………………………………… S42
Figure S28. MS/MS spectrum of CAR646 ……….........................................………… S43
Figure S29. MS/MS spectrum of CAR680 …….…………………………...………… S44
Figure S30. MS/MS spectrum of CAR694 …..……………………………………..….. S45
Figure S31. Annotated Cladogram of Viridiplantae species ……………..…………… S46
NMR key correlations ………………..……………………….…………….………… S47
Figure S32. $^{1}$H-NMR spectrum of arabipeptin A ……………….………………….. S49
Figure S33. $^{1}$H-$^{1}$H-COSY NMR spectrum of arabipeptin A ………..……..…………… S50
Figure S34. $^{1}$H-$^{13}$C-HSQC NMR spectrum of arabipeptin A.……….……………….. S51
Figure S35. $^{1}$H-$^{13}$C-HMBC NMR spectrum of arabipeptin A ……………..…………… S52
Figure S36. $^{1}$H-$^{1}$H-NOESY NMR spectrum of arabipeptin A ……………..…………… S53
Figure S37. $^{1}$H-$^{1}$H-TOCSY NMR spectrum of arabipeptin A …………..……………… S54
Figure S38. Marfey's analysis for arabipeptin A ……………………..………………… S56

4. **Supplementary References** …………..…………………………….……….……….. S57

**Data Deposition**

*C. americanus* Sequencing (NCBI)

BioProject: PRJNA840870

BioSamples: SAMN28561782 (Leaf), SAMN28561783 (Stem), SAMN28561784 (Root)

SRA: SRR19334864 (Leaf), SRR19334863 (Stem), SRR19334862 (Root)

Plants

*Coffea arabica* purchased from JM Bamboo (Hacienda Heights, California, US) and voucher deposited as NCU Accession # 676128.

NMR

NMR of arabipeptin A was deposited to NP-MRD as NP0331083.

Mass Spectrometry

Mass spectrometry data of arabipeptin A was deposited to MassIVE as MSV000090329.

Mass spectrometry metabolomic data for *Ceanothus americanus*, *Coffea arabica* and *Hibiscus syriacus* were deposit to MassIVE as MSV000090329.

**Metabolomics**

Extraction

*C. americanus* compound presence confirmation was performed in a two-step extraction. Extractions used 1 to 2 g of fresh plant tissue (root, stem, or leaf). The dried, ground plant tissue powder was extracted first in petroleum ether overnight. The filtrate was kept and added to methanol and extracted overnight and filtered. After rotary evaporation, the residue was resuspended in water with 0.1% formic acid and then fractionated with 15, 30, 45, and 60% aqueous acetonitrile and 0.1% formic acid using a *Hypersep* C18 column (Thermo Fisher). These fractions were dried with a Savant SpeedVac SPD120 and the remaining residue was resuspended in 50% acetonitrile and water with 0.1% formic acid (ACN + 0.1% FA) for analysis.

Using dry, ground roots of *C. arabica* (7.87 g) and *H. syriacus* (2.60 g), methanol extractions were carried out overnight. Crude methanol extracts were filtered, concentrated *in vacuo* and resuspended in 10 mL of acidic water (0.1% of formic acid). 2 mL were applied into Hypersep C18 200 mg columns (Thermo Scientific) and fractionation with 15, 30, 45, 60 and 100% of acetonitrile with 0.1% of formic acid was performed. Fractions 15, 30, 45 and 60% were pooled together and concentrated in a Thermo Scientific SPD120 speedvac. For UPLC-MS analysis, concentrated fractions were resuspended in 500 µL of acidic water (0.1% of formic acid).

Analysis

Extracts were analyzed using a UPLC-MS system with a Acquity UPLC (Waters), and LC column Acquity UPLC BEH 1.7 µm C18 reverse phase 130Å 2.1x50 mm. The following gradient was used: 0 min, 5% B; 1 min, 5% B; 11 min 100% B; 12 min, 100% B; 12.1 min, 5% B; 13 min, 5% B where solvent A was water with 0.1% of formic acid and solvent B was acetonitrile with 0.1% of formic acid. Mass spectrometric analysis was completed in a Q Exactive Plus (Thermo Scientific) in positive ion mode with a mass range 300-1,000 *m/z* and dd-MS2 (data-dependent MS/MS), collision energies at 20 eV, 25 eV and 30 eV. LC-MS data were analyzed using the Qual Browser Thermo Xcalibur software package (v.3.0.63 Thermo Scientific).

Molecular Network

Two molecular networks were created for *C. arabica* and *H. syriacus* following the workflow available at https://ccms-ucsd.github.io/GNPSDocumentation[1]. A third molecular network with *C. arabica* roots and *C. americanus* roots was generated as well with the aim of comparing *C. arabica* metabolic profile with a well-known cyclopeptide producing plant. Molecular networks were generated using the default settings and the following parameters: Network TopK = 10, Maximum Connected Component Size= 100, Min Pairs Cosine= 0.7 for *C. americanus and C. arabica*, and Min Pairs Cosine= 0.6 for *H. syriacus*. Molecular networking data were analyzed and edited with Cytoscape (v.3.9.1). Candidates cyclopeptides were identified by manual analysis of fragmentation patterns.

### RNA Extraction and Sequencing

RNA was extracted from the leaf, stem, and root of *C. americanus* using method five from the previous publication by Sim, Ho, and Phang.[2] Briefly, this method utilized an extraction buffer composed of 100 mM Tris-HCl (pH 8.0), 2 M NaCl, 20 mM EDTA, 2 % cetrimonium bromide (w/vol), and 50 mM dithiothreitol. The extraction buffer was added to a flash-frozen ground sample at a 1:10 (tissue:buffer) ratio and centrifuged. The extraction was followed by the addition of ethanol (0.3 vol) and a first round of chloroform-isoamyl (C:I) alcohol (24:1) extraction which was followed with centrifugation. After C:I purification LiCl was added to a final concentration of 2 M. After precipitation at -80 $^{o}$C for 2 hours after incubation, the pellet was retrieved by centrifugation and dissolved in 400 µL of diethyl pyrocarbonate (DEPC) treated water. After this, another round of C:I purification, centrifugation, and LiCl precipitation was carried out. After another two hours in the -80 $^{o}$C freezer the pellet was once again isolated and dissolved in 50 µL DEPC water. RNA extractions started with 0.2 to 0.3 g of material and yielded between 0.8 to 2.5 µg of RNA. The quality of the RNA was analyzed using both SpectraDrop and the Biotium Broad Range RNA Quantification kit.

Purified RNA samples were sent to the University of North Carolina at Chapel Hill High Throughput Sequencing Facility (HTSF) lab for quality checks and sequencing. The RNA Sequencing was completed using an Illumina NextSeqP1 at 2x100PE. The library was prepared using KAPA RNA HyperPrep Kit with RiboErase. Sequence yield for each sample was as follows: 21.1 million 2x100PE reads for leaf, 25.0 million 2x100PE reads for stem, and 22.5 million 2x100PE reads for root. See Data Deposition section for accession codes.

### Transcriptome Assembly and Analysis

We used a Trinity, TransDecoder, and CD-HIT pipeline to assemble and analyze the RNA-Seq data to discover the cyclopeptide alkaloid precursor peptides. Trinity[3,4] was used with default settings to yield a combined leaf, stem, and root assembly of 67,599 total transcripts with 41,548 Trinity 'genes' and a 42.30% GC content. The leaf, stem, and root data sets were also assembled separately. After using TransDecoder[4] to translate the transcripts, the assembly was then searched for all of the predicted cores that could lead to the *C. americanus* metabolomic data (Figure S1-S8). From the root alone, this generated 2189 transcripts with a putative core. In order to identify authentic precursor peptides, we relied on the conserved logic of RiPP biosynthetic pathways.[5] Typically, precursor peptides contain a conserved leader sequence responsible for recognition by the RiPP tailoring enzymes (Figure 1A). The variable core region is modified and subsequently released from the leader as the final natural product. If the *C. americanus* cyclopeptide alkaloids followed this model, the precursor peptides should be largely conserved by amino acid sequence and contain short variable regions that correspond to core sequences. Therefore, we clustered the candidate sequences by amino acid sequence similarity using the *CD-HIT* UCSD web server.[6] By using a 40% amino acid sequence identity threshold, we generated 441 groups of potential transcripts. These groups were visually analyzed to look for clusters composed of different putative cores within transcripts that had high levels of amino acid sequence similarity. This allowed us to identify a cluster containing putative cores that matched seven of the cyclopeptide alkaloids found in *C. americanus*. The entire transcriptome was later searched using our custom Hidden Markov Model to identify all precursor peptide transcripts.

**<u>Hidden Markov Model & Cladogram</u>**

A list of candidate peptides was generated using a PSI-BLAST of the *C. americanus* precursor sequence (Supplementary Table). The resulting protein sequences were aligned and used to create a custom Hidden Markov Model using the hmmbuild function of HMMER v3.3.2.[7] The NCBI Viridiplantae database (17,867,506 sequences) was downloaded on March 20, 2022 and analyzed with our precursor peptide HMM at the default E-value of 10 as to include proteins with even low levels of amino acid similarity. The BURP-domain protein family HMM[8] was downloaded on March 18, 2022 and ran against the Viridiplantae sequences with an E-value cutoff of 10. Hits that were common to both results were considered fused BURP-domain systems, while those only present in the precursor peptide HMM results were considered split. Lists of protein accessions for fused and split systems were then made. Species names were extracted from the lists and duplicates were removed.

Species names from eudicots and monocots with annotated genomes were downloaded from NCBI on June 9, 2022 and May 5, 2022 respectively. These were used to calculate percent abundance by counting exact matches between our species lists and the NCBI species. We found that 89.1% of the 202 annotated eudicot genomes deposited to NCBI contain precursor peptides from split BURP-domain systems. Additionally, 79.2% contain fused BURP-domains related enough to stand alone precursor peptides to be identified from our HMM. In contrast, fused BURP-domains were more prevalent in monocots with 83.7% of the 49 unique monocot species with annotated genomes containing at least one. Split BURP–domain systems were present in only 32.7% of annotated monocot genomes.

Finally, the species identified as having fused and split BURP-domain systems were used to create a cladogram. Species names were converted to taxids for use with phyloT v2[9]. The resulting tree was annotated with clade information, presence of a stand alone precursor peptides, fused BURP-domains, and presence of a known natural product derived from the split BURP-domains using the R ggtree and ggtreeExtra packages.

**Sequence Similarity Network**

A sequence similarity network (SSN1) was constructed from the list of split candidates using the Enzyme Function Initiative enzyme similarity tool.[10] The network was colorized and then visualized in Cytoscape v3.9.1 with an alignment score of 70. A second sequence similarity network was constructed at an alignment score of 30 from the singlets in SSN1 and also visualized in Cytoscape. Cores from each sequence were annotated if the sequence matched a known molecule or if the sequence followed a motif similar to the NxxxYH/R motif found in *C. americanus.*

**Purification of arabipeptin A**

Arabipeptin A was extracted from 30 ground *Coffea arabica* plants purchased from JM Bamboo (Hacienda Heights, California, US). 24.40 g of dried roots was obtained and sequentially extracted with methanol twice for 4 h and overnight, each time with a volume of 500 mL. Crude methanol extracts were filtered, concentrated *in vacuo* and resuspended in 75 mL of acidic water (0.1% formic acid). The aqueous solution was extracted five times with 1-butanol (75 mL). Pooled organic layers were concentrated *in vacuo* and resuspended in 25 mL of methanol and further dried by vacuum centrifugation in a Thermo Scientific SPD120 speedvac. The dried material (1.9 g) was dissolved in 2 mL of water and fractionated *via* flash chromatography on a CombiFlash Rf (Teledyne ISCO) with the following parameters: injection concentration 950 mg/mL; RediSep High Performance 150 g HP C18 reverse phase column; LC gradient solvent A water; solvent B acetonitrile; 0 column volume (CV) 10% B, 1 CV 10% B, 13 CV 40% B, 5 CV 40% B; flow rate 85 mL/min; 25 mL fractions were collected; wavelength window 200-300 nm; evaporative light scattering detection was used to observe the relative quantity of material.

Selected fractions containing arabipeptin A (as observed by mass spectrometry) were further subjected to preparative HPLC on a Varian Pro Star chromatography system coupled to a Finnigan LTQ (Thermo) mass spectrometer using the following parameters: 41 mg and 24 mg of fraction were dissolved in 1 mL each of methanol and injected separately; Atlantis Prep T3 5µm 19x250 mm LC column; LC gradient solvent A water (0.1% of formic acid); solvent B acetonitrile (0.1% of formic acid); 0 min, 15% B, 30 min 45% B, 5 min 45% B; flow rate 15 mL/min; fraction volume of 7.5 mL; wavelengths 220 nm and 280 nm and the wavelength window 190-400 nm; ELSD and MS data collected concurrently. Resolution of arabipeptin A containing fractions by semipreparative HPLC was carried out on the same instrument with the following parameter changes: 4.8 mg of fraction was dissolved in 0.2 mL of methanol and injected; Semi Prep Luna 5µm PFP(2) 100Å 250x10 mm LC column; LC gradient solvent A water (0.1% of formic acid); solvent B acetonitrile (0.1% of formic acid); 0 min, 15% B, 30 min 45% B, 5 min 45% B; flow rate 4.70 mL/min; fraction volume of 2.35 mL.

**Structure elucidation of arabipeptin A**

MS/MS fragmentation pattern of arabipeptin A revealed putative iminium ion mass for phenylalanine and other consistent fragments related to the core peptide sequence and proposed molecule (Figure S26). The final product of purification (arabipeptin A) was analyzed by 1D and 2D NMR in methanol-$d_4$ in 5 mm NMR tubes (Wilmad LabGlass). NMR instrumentation was a JEOL ECA-500 MHz NMR spectrometer (JEOL Ltd.). $^1$H NMR (500 MHz, METHANOL-$D_3$) δ 7.47 (d, *J* = 7.9 Hz, 1H), 7.26 (d, *J* = 8.0 Hz, 1H), 7.20 (d, *J* = 4.4 Hz, 4H), 7.11 (m, 1H), 7.01 (s, 1H), 7.03 (t, *J* = 7.5 HZ, 1H), 7.01 (s, 1H), 6.97 (dd, *J* = 8.0, 7.0 Hz, 1H), 6.87 (d, *J* = 8.5 Hz, 1H), 6.82 (d, *J* = 8.5 Hz, 1H), 6.73 (d, *J* = 8.4 Hz, 1H), 6.61 (d, *J* = 8.0 Hz, 1H), 4.62 (d, *J* = 10.1 Hz, 1H), 4.42 (dd, *J* = 10.1, 2.0 Hz, 1H), 4.33 (m, 1H), 4.28 (t, *J* = 6.8 Hz, 1H), 3.77 (d, *J* = 17.0 Hz, 1H), 3.43 (dd, *J* = 9.1, 5.2 Hz, 1H), 3.15 (m, 1H), 3.07 (dd, *J* = 13.7, 9.2 Hz, 1H), 3.00 (d, *J* = 12.6 Hz, 1H), 2.98 (m, 2H), 2.83 (dd, *J* = 7.1 Hz, 1H), 2.43 (s, 6H), 1.97 (p, J = 6.3 Hz, 1H), 0.98 (d, *J* = 6.8 Hz, 3H), 0.92 (d, *J* = 6.8 Hz, 3H); HRMS (EIC) calculated for $C_{39}H_{46}N_6O_7$ 711.3501, found 711.3503 (Δ 0.2904 ppm) $[M+H]^+$. Arabipeptin A macrocyclization between Tyr5 and Leu2 is supported by HMBC correlations between Leu2-Hβ (δ 4.42 ppm, 1H) and Tyr5-C4 (δ 157.8 ppm) and NOESY correlations between Leu2-Hβ (δ 4.42 ppm, 1H) and Tyr5-H3/5 (δ 6.61 ppm, 1H/δ 6.73 ppm, 1H). Stereochemistry of the ether bond is proposed to be in the *S*-configuration based on *J*-coupling constants. The observed *J*-coupling constants ($J_{\alpha\text{-}\beta}$(Leu2) = 10.1 Hz and $J_{\beta\text{-}1}$(Leu2) = 2.0 Hz) indicate an L-erythro (*anti*) configuration at the $C_\alpha$-$C_\beta$-bonds of Leu2 based on literature precedent. Related 14-membered ring cyclopeptide alkaloids with reported *J*-values for the same bonds have similar coupling constants ($J_{\alpha\text{-}\beta}$ = ~8.0 Hz) when they are in L-erythro (*anti*) conformation.[11,12] In contrast, the threo (*syn*) confirmation is reported to have $J_{\alpha\text{-}\beta}$ = ~2.0 Hz.[12] Moreover, our observed $J_{\alpha\text{-}\beta}$(Leu2) = 10.1 Hz is consistent with an *anti*-conformation as predicted by the Karplus equation. Finally, this assignment is consistent with our observed NOESY spectra.

### Marfey's reaction - arabipeptin A

For each amino acid standard to be tested, 0.2 mg of the amino acid was weighed into 4 mL reaction vials. To each amino acid were added 50 µL of water, 20 µL of 1 M $NaHCO_3$, and 100 µL 1% Marfey's reagent (Na-(2,4-dinitro-5-fluorophenyl)-L-alaninamide, Thermo Scientific) in acetone. The reaction mixture was incubated at 40 ºC for 1 h with periodic agitation. The reactions were quenched by adding 10 µL of 2 N HCl and dried under a stream of nitrogen. Dried derivatized reactions were resuspended in ~1.7 mL of MeOH and injected individually (2 µL), and as a combined mixture (2 µL) onto UPLC-MS (Waters Acquity UPLC system coupled to LTQ Orbitrap XL mass spectrometer), with the following parameters: gradient conditions were 15-85% acetonitrile (0.1% of formic acid) and water (0.1% of formic acid) over 8 min, on a BEH C18 column, and the derivatized products were observed at 340 nm as well as monitored by their expected $[M+H]^+$ ion.

For arabipeptin A, long (24 h) and a short (4 h) incubations were carried out to assure the hydrolysis of the ether bond and integrity of the tryptophan residue, respectively. Hydrolyzed and derivatized arabipeptin A was generated by adding 0.5 mL of 6 N HCl to a 0.2 mg sample in a 4 mL reaction vial, incubating for 24 or 4 h at 90 ºC, and then evaporation under a stream of nitrogen. To the hydrolysis product was added 25 µL of water, 10 µL of 1 M $NaHCO_3$ and 50 µL of 1% Marfey's reagent in acetone. This was incubated for 1 h at 40 ºC with periodic agitation. The reaction was quenched by adding 5 µL of 2 N HCl, dried under a stream of nitrogen and resuspended in 200 µL of MeOH. The sample was injected (6 µL) onto UPLC-MS using the same conditions and the same instrument as amino acids standards. Tryptophan standards were digested and derivatized concurrently with arabipeptin A to show that the tryptophan is degraded during 24 h digestion, but not in the 4 h digestion.

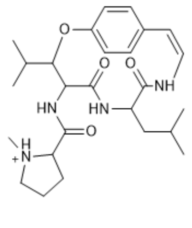

ceanothine C (Leu)
Exact Mass: 471.2966
PLLY

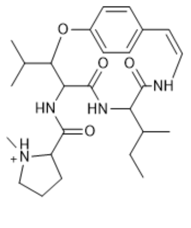

ceanothine C (Ile)
Exact Mass: 471.2966
PLIY

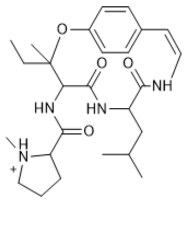

ceanothine D
Exact Mass: 471.2966
PILY

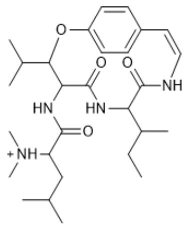

adouetine X
Exact Mass: 501.3435
LLIY

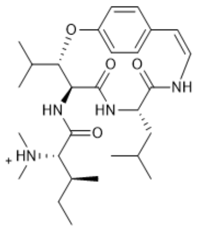

frangulanine
Exact Mass: 501.3435
ILLY

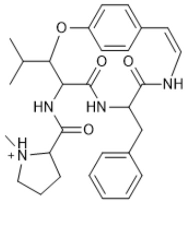

ceanothine B
Exact Mass: 505.2809
PLFY

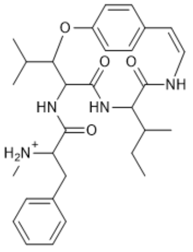

ceanothine A (Ile)
Exact Mass: 521.3122
FLIY

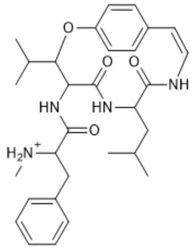

ceanothine A (Leu)
Exact Mass: 521.3122
FLLY

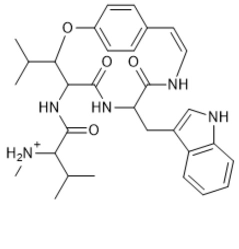

americine
Exact Mass: 546.3075
VLWY

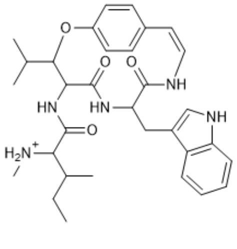

homoamericine (Ile)
Exact Mass: 560.3231
ILWY

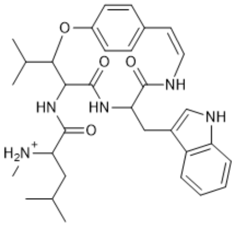

homoamericine (Leu)
Exact Mass: 560.3231
LLWY

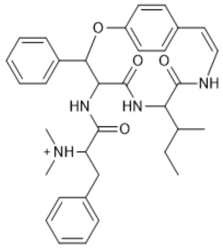

adouetine Y
Exact Mass: 569.3122
FFIY

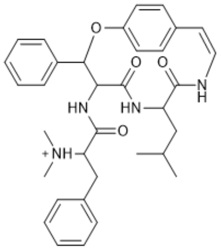

ceanothine E
Exact Mass: 569.3122
FFLY

**Figure S1.** Structures of the 10 cyclopeptide alkaloids proposed to be produced by *C. americanus* shown as [M+H$^+$]. Three molecules, ceanothine A, C, and homoamericine, have ambiguity about the identity of one amino acid. Both possibilities are shown. The hypothetical core amino acid sequence and exact mass is below each structure. Stereochemistry assignments are inconsistently reported in the literature. Frangulanine has been crystalized[13] and synthesized[14], giving strong evidence to stereochemical assignment. The remaining cyclopeptide alkaloid structures do not have strong evidence to support stereochemical assignment. Therefore, we have left them completely unassigned.

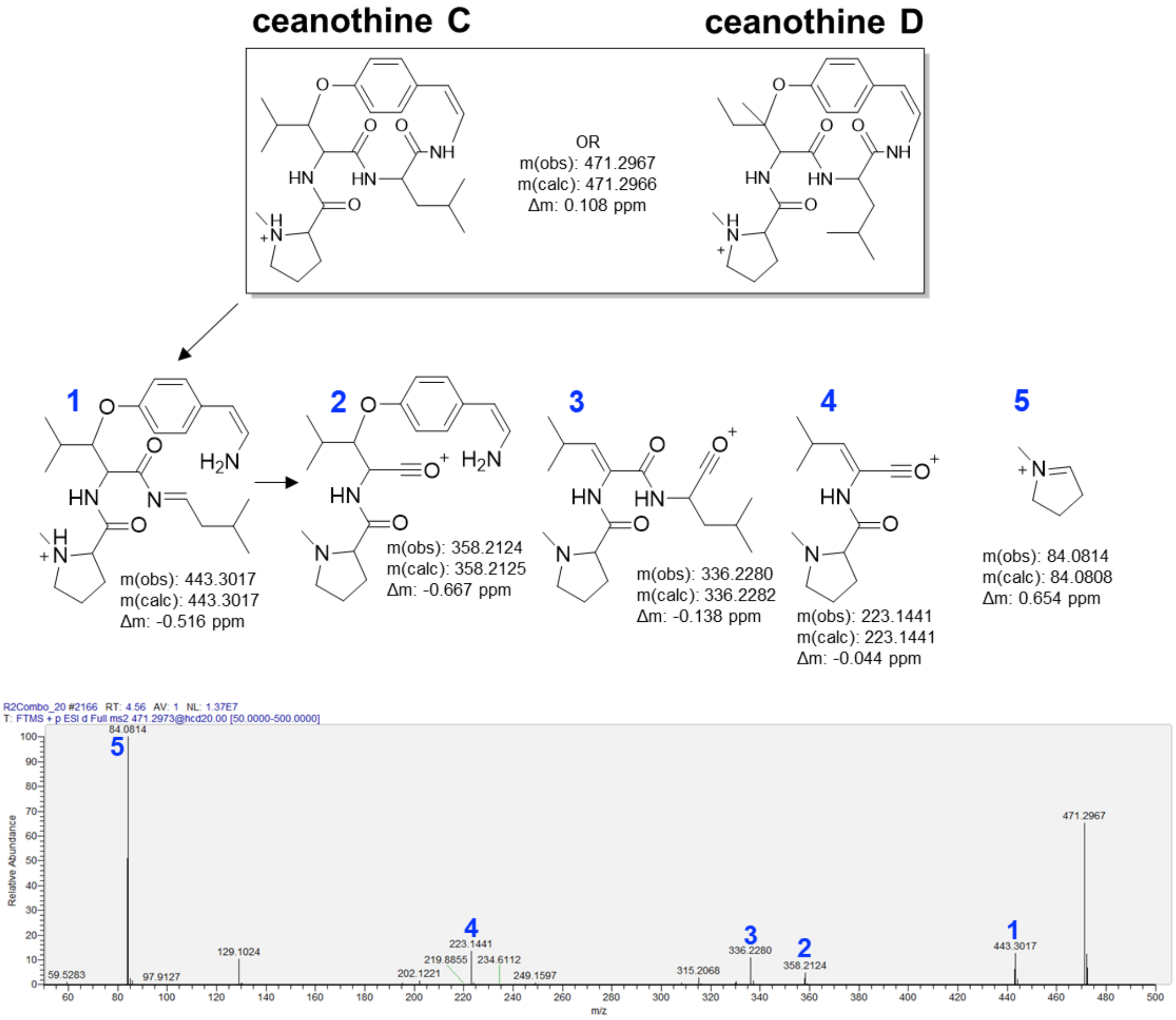


**Figure S2.** MS/MS spectrum of a feature from *C. americanus* root extract consistent with ceanothine C or ceanothine D.

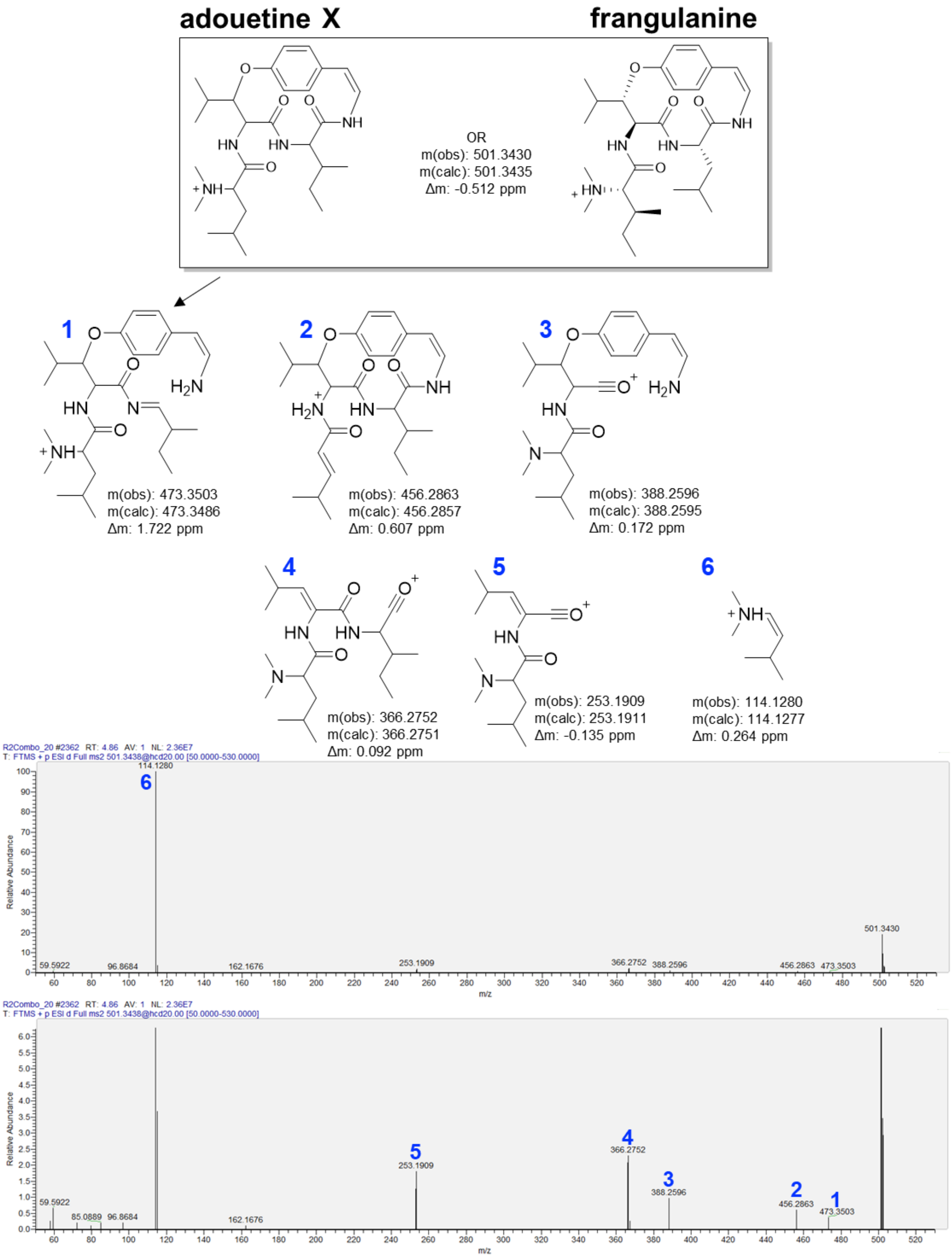


**Figure S3.** MS/MS spectrum of a feature from *C. americanus* root extract consistent with adouetine X or frangulanine.

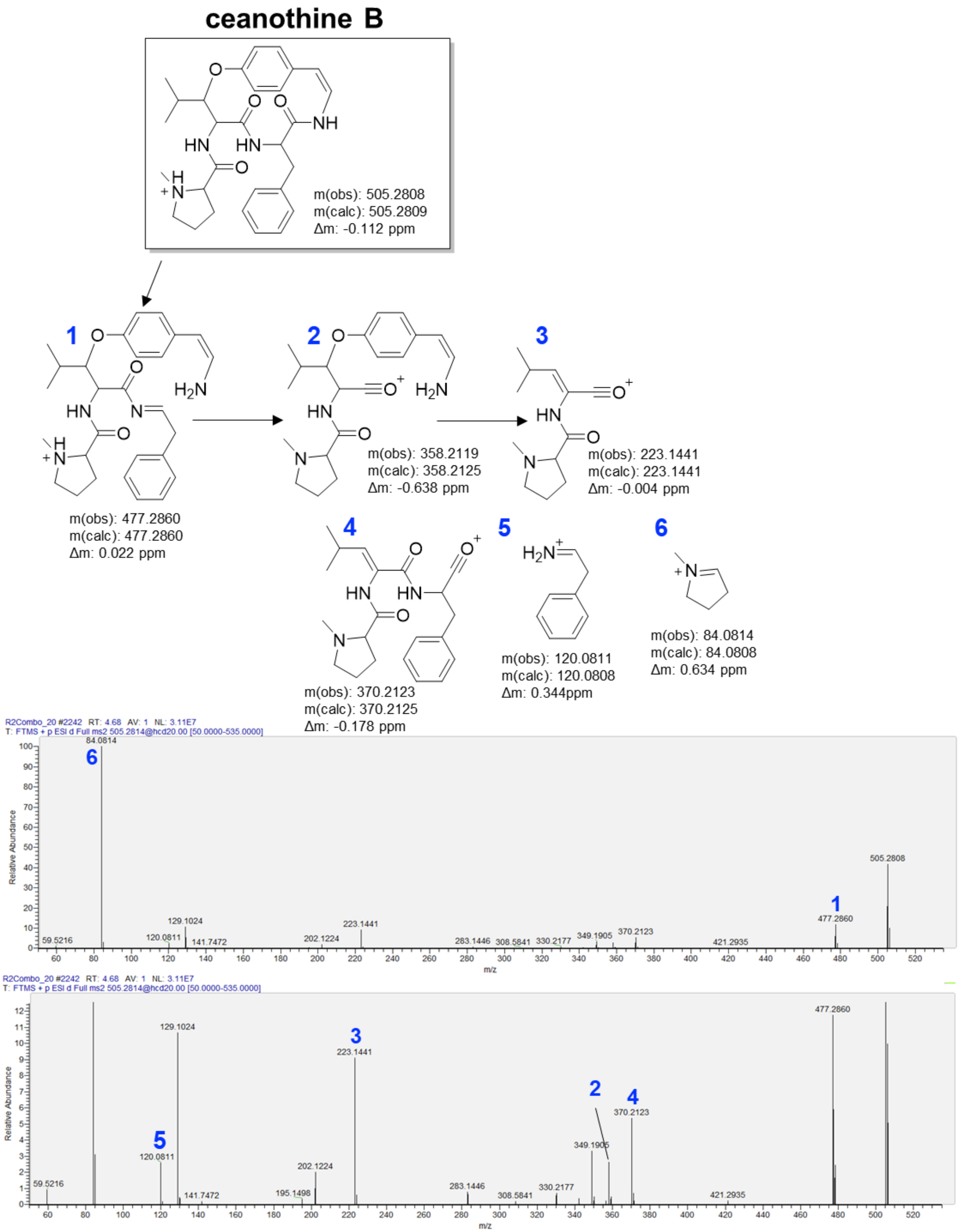


**Figure S4.** MS/MS spectrum of a feature from *C. americanus* root extract consistent with ceanothine B.

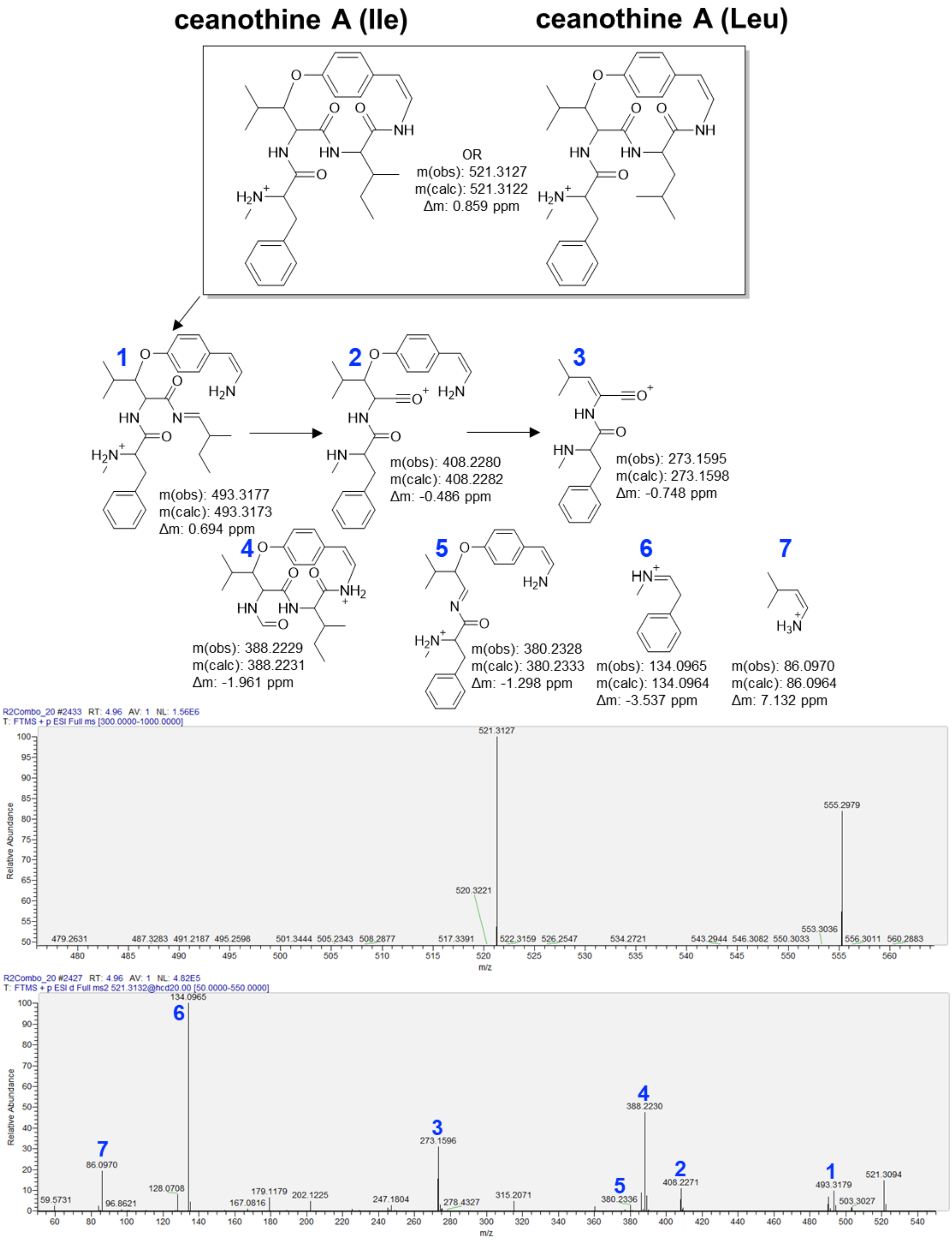


**Figure S5.** MS/MS spectrum of a feature from *C. americanus* root extract consistent with ceanothine A.

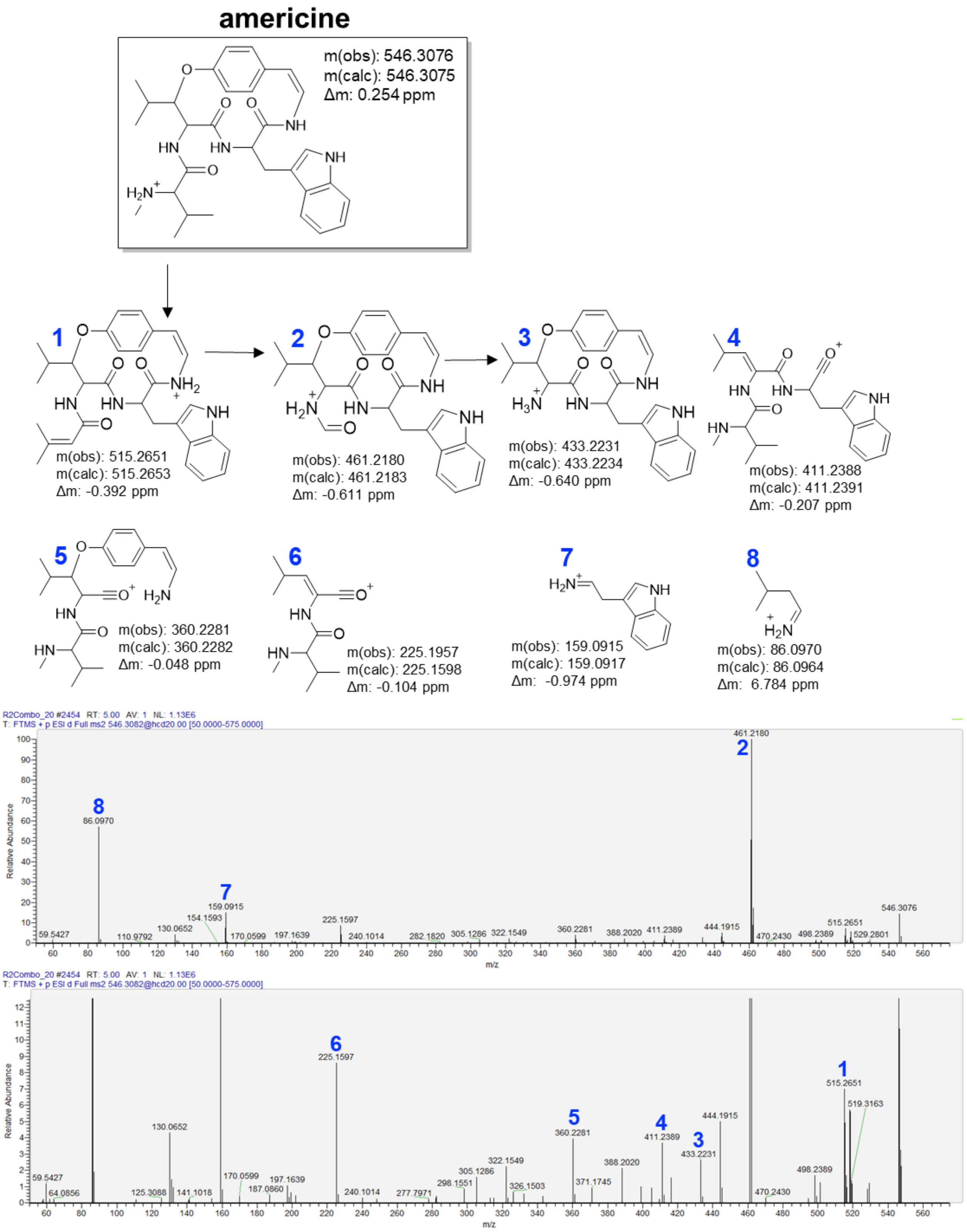


**Figure S6.** MS/MS spectrum of a feature from *C. americanus* root extract consistent with americine.

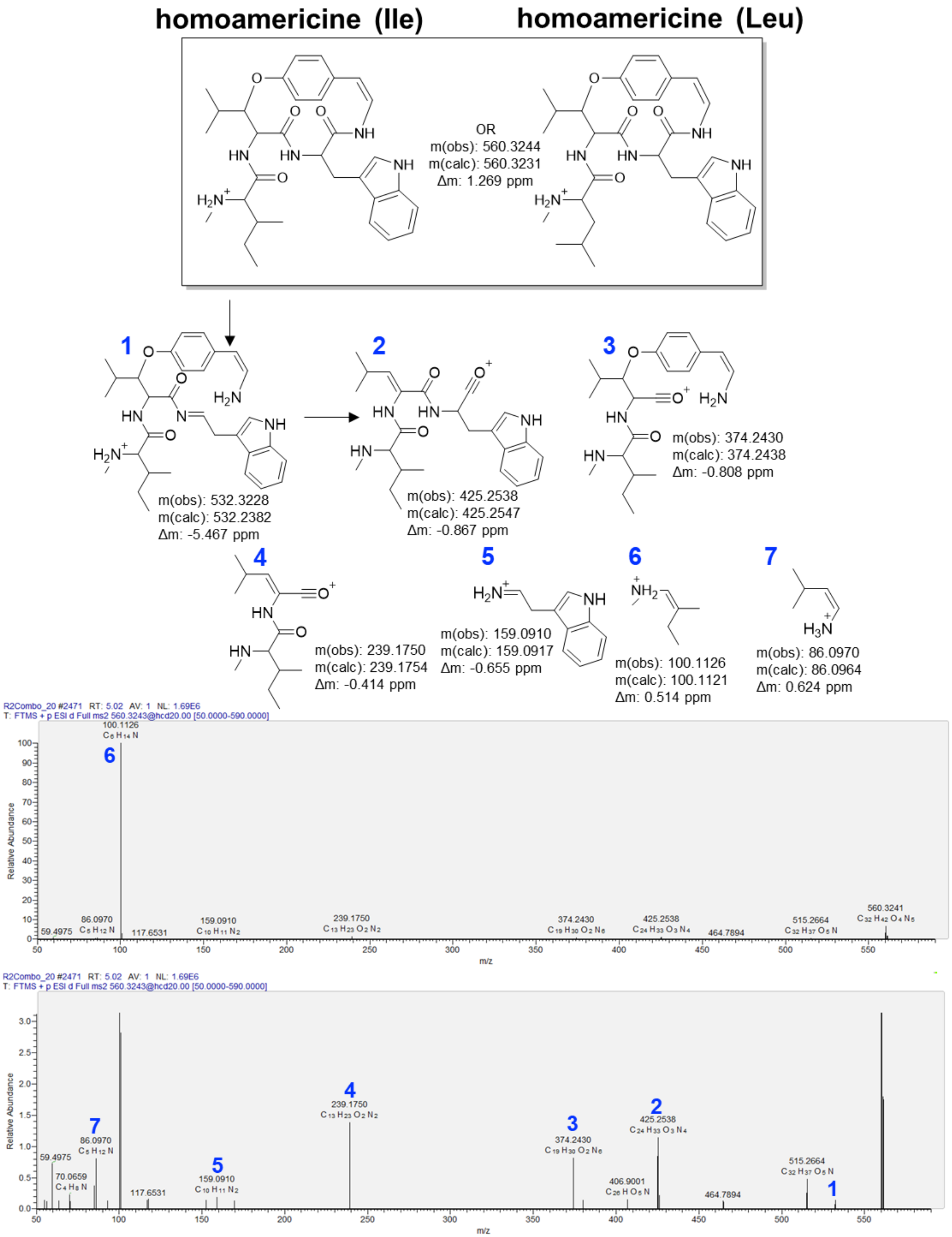


**Figure S7.** MS/MS spectrum of a feature from *C. americanus* root extract consistent with homoamericine.

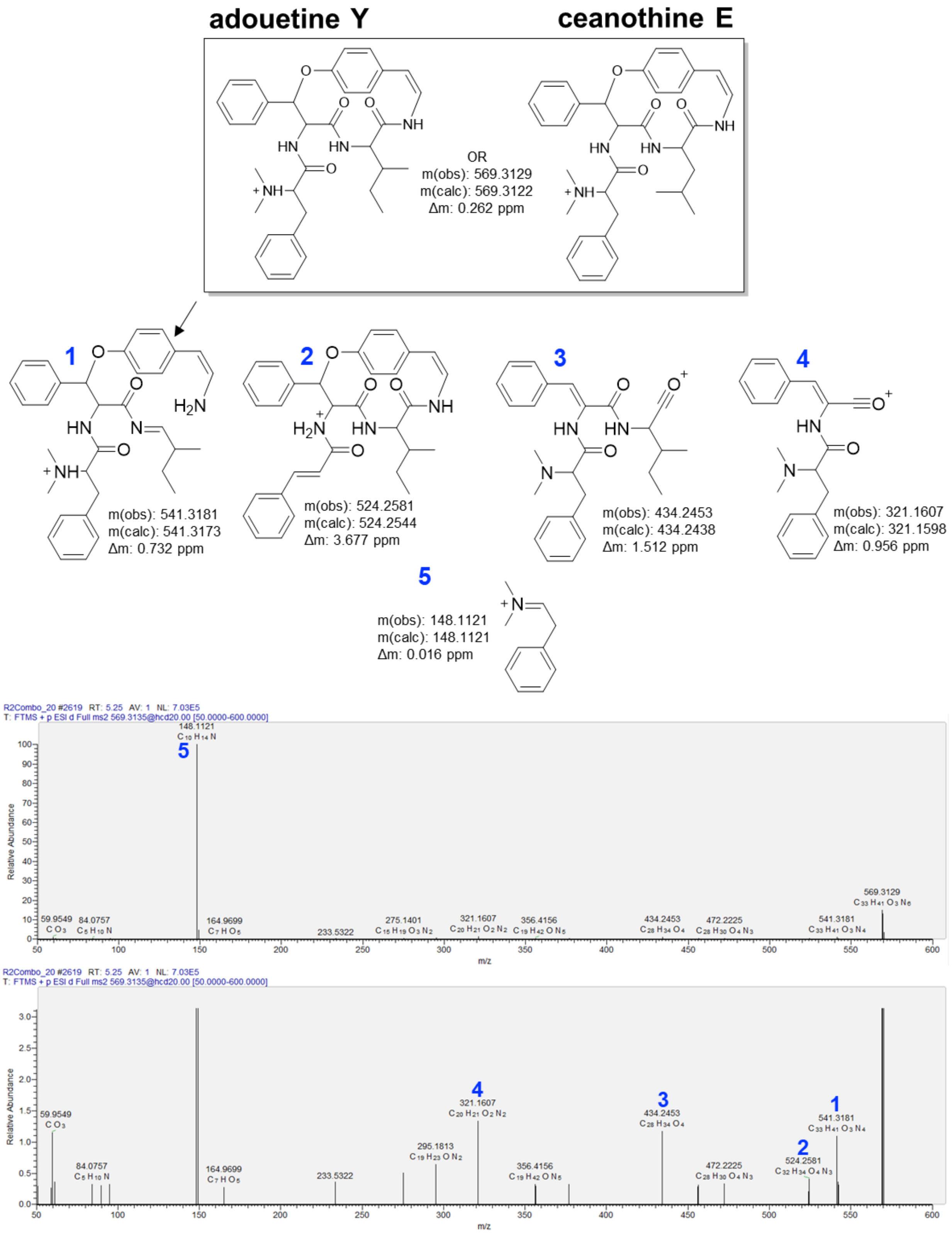


**Figure S8.** MS/MS spectrum of a feature from *C. americanus* root extract consistent with adouetine Y and ceanothine E.

>Ceanothus_americanus_27101
MKTFFALFAFSSLLLLSSTITARKEPEYLKTVIENQPILEVLQGV
LDLITKSGNGGRVAKDISVDPSGNILWYHGNKNAEAKSKDDGEHK
DDFSVDPSGNELWYHDNKNAEAKSKADGEHKDDFSVDPSGNELWY
HDNKNAEAKSINPSGN**VLWY**HGNKNAEAKSINPSGN**VLWY**HGNKN
AEAKSKADGEHKDDFSVDPSGN**VLWY**HGKKIVELDSETIAEQVAK
AFSDAPSGTFILYHGNKNGESNSKASGEGAAKDSFVDPSVFDK

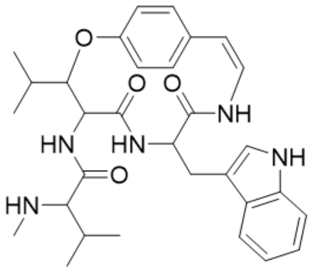

americine | VLWY

>Ceanothus_americanus_27097
PKKHFEFFEVDIKMKTFFALFAFSSLLLLSSTITARKEPEYLKTV
IENQPILEVLQGVLDLITKSGNGGRVAKDISVDPSGNILVYHGNK
NGEHVPKDASVDPSGNILLYHGNQNGKQVAKDFSVDPSGN
**FFIY**HNNKNAAAKSKADG

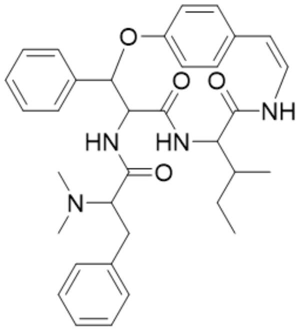

adouetine Y | FFIY

>Ceanothus_americanus_27105
SVDPSGN**ELWY**HDNKNAEAKSKADGEHKDDFSVDPSGN**ELWY**HDNK
NAEAKFLWYHGNKNAEAKSKADGEHKDDFSVDPSGNVLWYHGKKIV
ELDSETIAEQVAKAFSDAPSGTFILYHGNKNGESNSKASGEGAAKD
SFVDPSVFDK

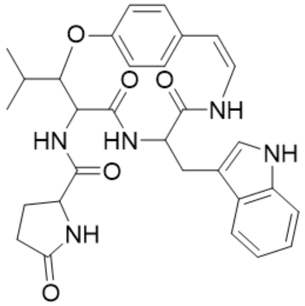

CAM544 | ELWY

>Ceanothus_americanus_27100
**ILWY**HGNKNAEAKSKDDGEHKDDFSTDPSGNFFLYHGNKIVELNSE
TNIAEQVAKAFSVAPSGTFIINLGNKMVSKM

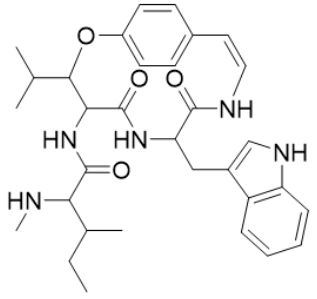

homoamericine | ILWY

```
>Ceanothus_americanus_27108
MKTFFVLLAFSSLLLSRTITARKEPAEYAKTIMENQPIPVLEVIQGVL
DLIAKSSRNGGRLAKDLSVDPLSNWLIYRGNQNGENVAKDLKDLSVDP
SSNFLIYRGNHNGENAAKDLSVDPLSNWLIYRGNQNGENVAKDLKDLS
VDPSSNFLIYRGNHNEENVAKDLSVDPSSNFLIYRGNHNEENVAKDLS
VDPSSNFLIYRGNHNEENVAKDLSVDPSSNFLIYHGNKNADPNSKANG
EGVAKGISPDHHDQ
```

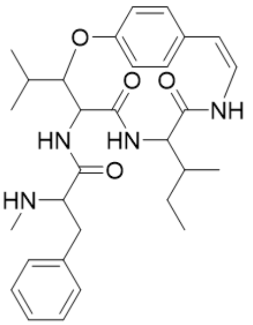

ceanothine A | FLIY

```
>Ceanothus_americanus_27111
GNFFFYHNNKNAVAKSKADEGELKDFSVDPSGNFFFYHNNKNAVAKSK
ADEGELKDFSVDPSGNFFFYHNNKNAVAKSKADEGELKDFSVDPSGNF
FFYHNNKNAVAKSKADEGELKDFSVDPSGNILLYHGNKNAEVKSNTDG
KHKEDFSVDPSGNPLFYHGNKNDDFSVDPSGNPLFYHGNKNGEH
```

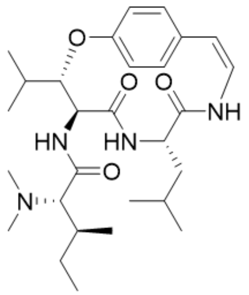

frangulanine | ILLY

```
>Ceanothus_americanus_27116
GNFFFYHNNKNAVAKSKADEGELKDFSVDPSGNFFFYHNNKNAVAKSKA
DEGELKDFSVDPSGNILLYHGNKNAEVKSNTDGKHKEDFSVDPSGNPLL
YHDNKNDDFSVDPSGNPLLYHDNKNGEHKDGFSVDPSGNPLFYHGNKND
DFSVDPSGNPLFYHGNKNGEHKDGFSVDPSGNPLLYHDNKNDDFSVDPS
GNPLFYHGNKNGEHKDGFSVDPSGNPLFYHGNKNGEHKDGFSVDPSG
```

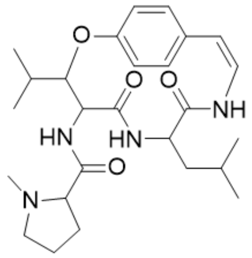

ceanothine C | PLLY

```
>Ceanothus_americanus_27100
ILWYHGNKNAEAKSKDDGEHKDDFSTDPSGNFFLYHGNKIVELNSETNI
AEQVAKAFSVAPSGTFIINLGNKMVSKM
```

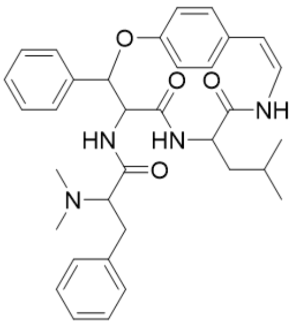

ceanothine E | FFLY

>Ceanothus_americanus_27112
GNFFFYHNNKNAVAKSKADEGELKDFSVDPSGNFFFYHNNKNAVAKSKA
DEGELKDFSVDPSGNFFFYHNNKNAVAKSKADEGELKDFSVDPSGNFFF
YHNNKNAVAKSKADEGELKDFSVDPSGNILLYHGNKNAEVKSNTDGKHK
EDFSVDPSGNPLLYHDNKNDDFSVDPSGNPLLYHDNKNGEHKDGFSVDP
SGNPLFYHGNKNDDFSVDPSGNPLFYHGNKNGEHKDGFSVDPSGNPLLY
HDNKNDDFSVDPSGNPLFYHGNKNGEHKDGFSVDPSGNPLFYHGNKNGE
HKDGFSVDPSG

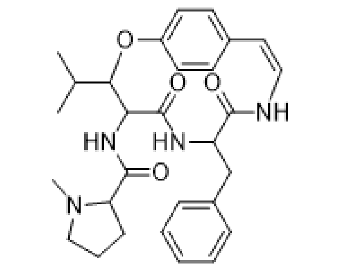


ceanothine B | PLFY

>Ceanothus_americanus_27112
GNFFFYHNNKNAVAKSKADEGELKDFSVDPSGNFFFYHNNKNAVAKSKA
DEGELKDFSVDPSGNFFFYHNNKNAVAKSKADEGELKDFSVDPSGNFFF
YHNNKNAVAKSKADEGELKDFSVDPSGNILLYHGNKNAEVKSNTDGKHK
EDFSVDPSGNPLLYHDNKNDDFSVDPSGNPLLYHDNKNGEHKDGFSVDP
SGNPLFYHGNKNDDFSVDPSGNPLFYHGNKNGEHKDGFSVDPSGNPLLY
HDNKNDDFSVDPSGNPLFYHGNKNGEHKDGFSVDPSGNPLFYHGNKNGE
HKDGFSVDPSG

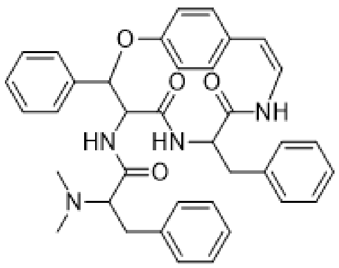


CAM603 | FFFY

**Figure S9.** Putative core sequences from the precursor peptides map to known cyclopeptide alkaloids from *C. americanus*. Core sequences corresponding to E-L-W-Y and F-F-F-Y are also shown.

>Ceanothus_americanus_27101

MKTFFALFAFSSLLLLSSTITARKEPEYLKTVIENQPILEVLQGVLDLITK
SGNGGRVAKDISVDPSGNILWYHGNKNAEAKSKDDGEHKDDFSVDPSGNEL
WYHDNKNAEAKSKADGEHKDDFSVDPSGNELWYHDNKNAEAKSINPSGNVL
WYHGNKNAEAKSINPSGNVLWYHGNKNAEAKSKADGEHKDDFSVDPSGNVL
WYHGKKIVELDSETIAEQVAKAFSDAPSGTFILYHGNKNGESNSKASGEGA
AKDSFVDPSVFDK*

>Ceanothus_americanus_27108

MKTFFVLLAFSSLLLSRTITARKEPAEYAKTIMENQPIPVLEVIQGVLDLI
AKSSRNGGRLAKDLSVDPLSNWLIYRGNQNGENVAKDLKDLSVDPSSNFLI
YRGNHNGENAAKDLSVDPLSNWLIYRGNQNGENVAKDLKDLSVDPSSNFLI
YRGNHNEENVAKDLSVDPSSNFLIYRGNHNEENVAKDLSVDPSSNFLIYRG
NHNEENVAKDLSVDPSSNFLIYHGNKNADPNSKANGEGVAKGISPDHHDQ*

**Figure S10.** Putative precursor peptides from *C. americanus* have an N-terminal signal peptide (yellow) as predicted by SignalP 6.0.[15] Repeated core sequences (red) are separated by repeated putative recognition sequences (blue). A core-less leader peptide-like region (un-highlighted) is found immediately after the signal peptide.

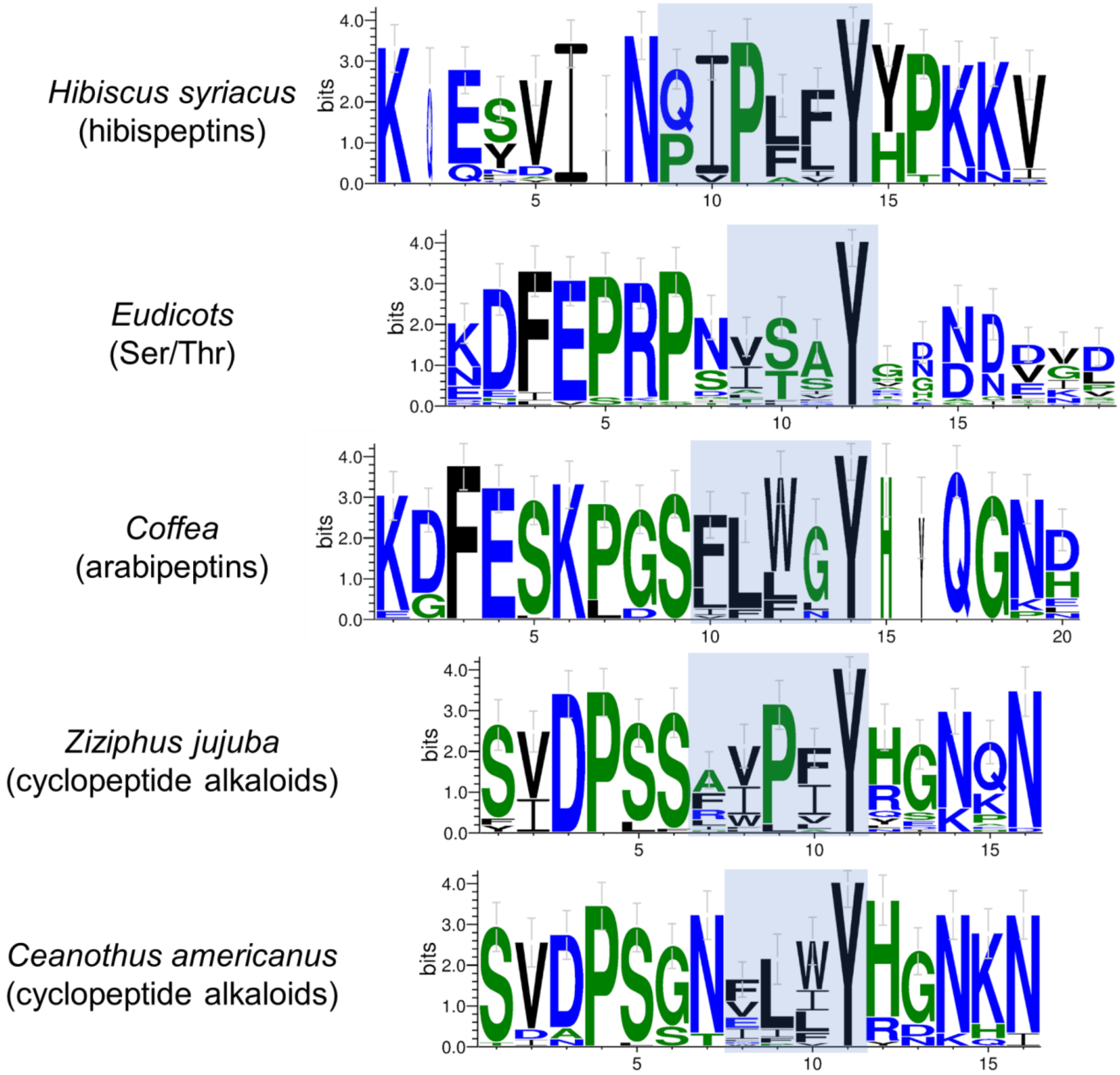


**Figure S11.** Weblogos generated with 30 aligned cores from the indicated SSN clusters. The highlighted region is the predicted core motif. The weblogos are aligned to the conserved Tyr.

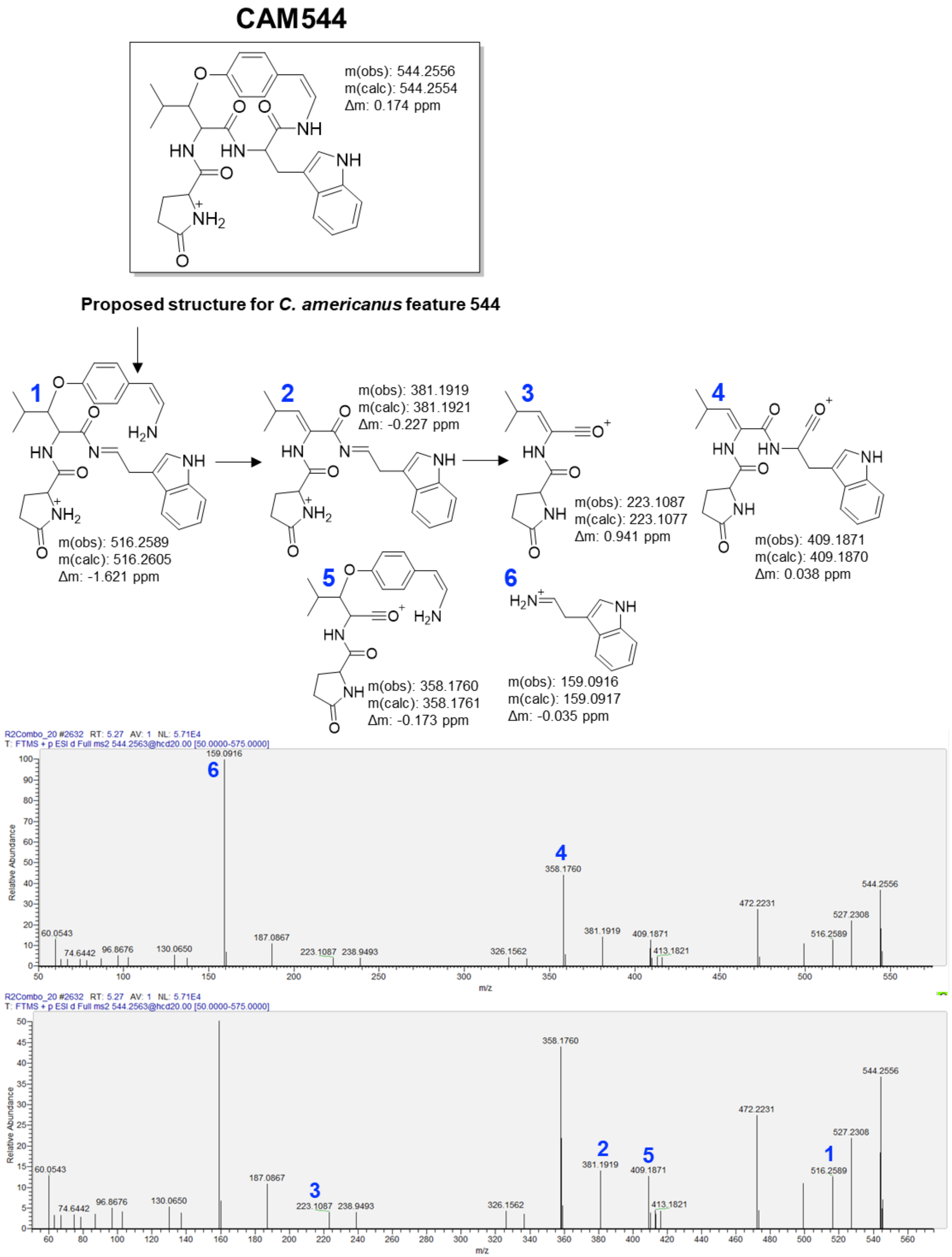


**Figure S12.** MS/MS spectrum of a feature from *C. americanus* root extract consistent with a cyclopeptide alkaloid derived from an E-L-W-Y core.

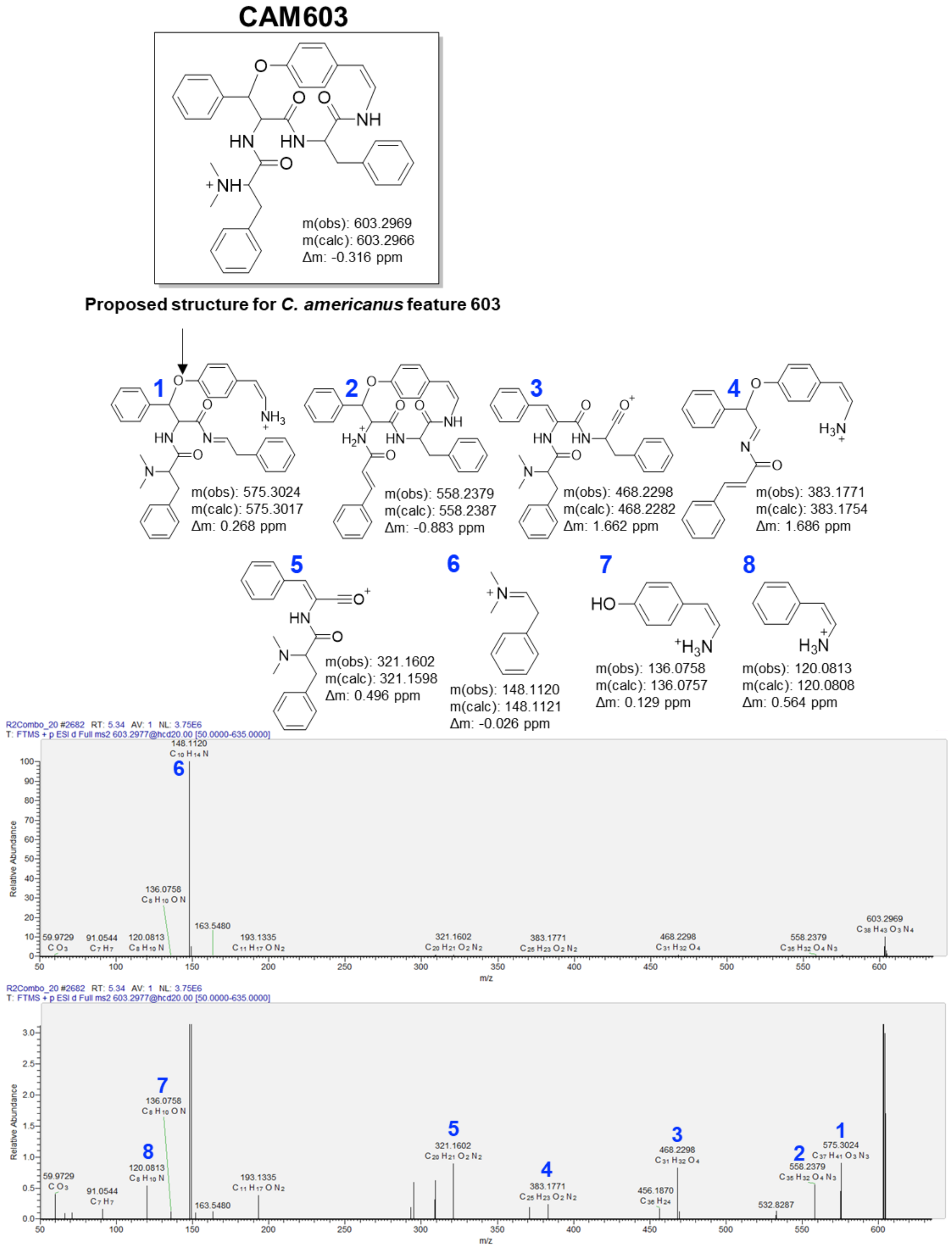


**Figure S13.** MS/MS spectrum of a feature from *C. americanus* root extract consistent with a cyclopeptide alkaloid derived from a F-F-F-Y core.

>KAH7517850.1 [Ziziphus jujuba var. spinosa]
MNLLDIILEHKGVGEKLARSCSGVVLVPPFALSSTIIARKEPVEY
VKTVVEQVAKDLSVDPSFAVPFYRSNKNGQSVDPSSAVPFHHGNK
NGLSIDPSSAIPVYRGNQNGQSIDPSAAIPIYHGNQNGQSIDPSS
ALLIYRINQNGLSIDPSSAVPFYHGNKNGFSIDPSSAVPFYHGNK
NGLSVDPSSAIPVYQGNQNGQSIDPSS**AVPVY**RSNQNGQSIDPSS
AVPIYHGNKNGLSVDPSSAIFVYRGNQNGQSIDPSSAIPVYRGNQ
NGQSIDPSSAIPVYRGNQNGQSIDPSSALLIYRINQNGLSIDPSS
AVPFYHGKKNGFFVDPSSAVPFYHGNKNGLSVDPSSAIPVYQGNQ
NGQSIDPSS**AVPVY**RSNQNGQSIDPSSAVPIYHGNKNGLSVDPSS
AIFVYRGNQNGQSIDPSSAIPIYHGNQNGQSIDPSS**AVPVY**RGNQ
NGQSIDPSSALLIYRINQNGLSIDPSSAVPFYHGNQNGLSMDLSS
AVPFYRGIKNGLSVDPSSAVPFYYENKNGLSVDPSSAVPFYRGNQ
NGEQNSKANEKGATKNLSTDQ

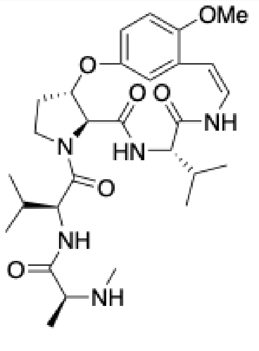

jubanine F I AVPVY

>KAH7517850.1 [Ziziphus jujuba var. spinosa]
MNLLDIILEHKGVGEKLARSCSGVVLVPPFALSSTIIARKEPVEY
VKTVVEQVAKDLSVDPSFAVPFYRSNKNGQSVDPSSAVPFHHGNK
NGLSIDPSS**AIPVY**RGNQNGQSIDPSAAIPIYHGNQNGQSIDPSS
ALLIYRINQNGLSIDPSSAVPFYHGNKNGFSIDPSSAVPFYHGNK
NGLSVDPSS**AIPVY**QGNQNGQSIDPSSAVPVYRSNQNGQSIDPSS
AVPIYHGNKNGLSVDPSSAIFVYRGNQNGQSIDPSS**AIPVY**RGNQ
NGQSIDPSS**AIPVY**RGNQNGQSIDPSSALLIYRINQNGLSIDPSS
AVPFYHGKKNGFFVDPSSAVPFYHGNKNGLSVDPSS**AIPVY**QGNQ
NGQSIDPSSAVPVYRSNQNGQSIDPSSAVPIYHGNKNGLSVDPSS
AIFVYRGNQNGQSIDPSSAIPIYHGNQNGQSIDPSSAVPVYRGNQ
NGQSIDPSSALLIYRINQNGLSIDPSSAVPFYHGNQNGLSMDLSS
AVPFYRGIKNGLSVDPSSAVPFYYENKNGLSVDPSSAVPFYRGNQ
NGEQNSKANEKGATKNLSTDQ

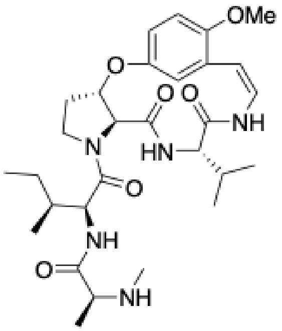

jubanine G I AIPVY

>KAH7517850.1 [Ziziphus jujuba var. spinosa]
MNLLDIILEHKGVGEKLARSCSGVVLVPPFALSSTIIARKEPVEYV
KTVVEQVAKDLSVDPSFAVPFYRSNKNGQSVDPSSAVPFHHGNKNG
LSIDPSSAIPVYRGNQNGQSIDPSA**AIPIY**HGNQNGQSIDPSSALL
IYRINQNGLSIDPSSAVPFYHGNKNGFSIDPSSAVPFYHGNKNGLS
VDPSSAIPVYQGNQNGQSIDPSSAVPVYRSNQNGQSIDPSSAVPIY
HGNKNGLSVDPSSAIFVYRGNQNGQSIDPSSAIPVYRGNQNGQSID
PSSAIPVYRGNQNGQSIDPSSALLIYRINQNGLSIDPSSAVPFYHG
KKNGFFVDPSSAVPFYHGNKNGLSVDPSSAIPVYQGNQNGQSIDPS
SAVPVYRSNQNGQSIDPSSAVPIYHGNKNGLSVDPSSAIFVYRGNQ
NGQSIDPSS**AIPIY**HGNQNGQSIDPSSAVPVYRGNQNGQSIDPSSA
LLIYRINQNGLSIDPSSAVPFYHGNQNGLSMDLSSAVPFYRGIKNG
LSVDPSSAVPFYYENKNGLSVDPSSAVPFYRGNQNGEQNSKANEKG
ATKNLSTDQ

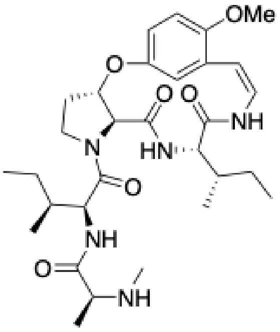

jubanine H I AIPIY

>KAH7517850.1 [Ziziphus jujuba var. spinosa]
MNLLDIILEHKGVGEKLARSCSGVVLVPPFALSSTIIARKEPVEY
VKTVVEQVAKDLSVDPSFAVPFYRSNKNGQSVDPSSAVPFHHGNK
NGLSIDPSSAIPVYRGNQNGQSIDPSAAIPIYHGNQNGQSIDPSS
ALLIYRINQNGLSIDPSSAVPFYHGNKNGFSIDPSSAVPFYHGNK
NGLSVDPSSAIPVYQGNQNGQSIDPSSAVPVYRSNQNGQSIDPSS
**AVPIY**HGNKNGLSVDPSSAIFVYRGNQNGQSIDPSSAIPVYRGNQ
NGQSIDPSSAIPVYRGNQNGQSIDPSSALLIYRINQNGLSIDPSS
AVPFYHGKKNGFFVDPSSAVPFYHGNKNGLSVDPSSAIPVYQGNQ
NGQSIDPSSAVPVYRSNQNGQSIDPSS**AVPIY**HGNKNGLSVDPSS
AIFVYRGNQNGQSIDPSSAIPIYHGNQNGQSIDPSSAVPVYRGNQ
NGQSIDPSSALLIYRINQNGLSIDPSSAVPFYHGNQNGLSMDLSS
AVPFYRGIKNGLSVDPSSAVPFYYENKNGLSVDPSSAVPFYRGNQ
NGEQNSKANEKGATKNLSTDQ

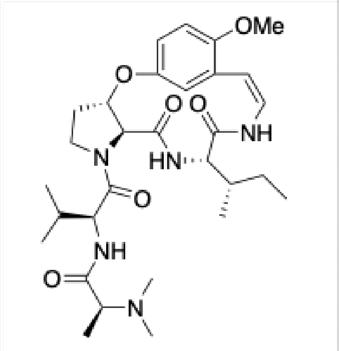


jubanine I I AVPIY

>KAH7517850.1 [Ziziphus jujuba var. spinosa]
MNLLDIILEHKGVGEKLARSCSGVVLVPPFALSSTIIARKEPVEY
VKTVVEQVAKDLSVDPSFAVPFYRSNKNGQSVDPSSAVPFHHGNK
NGLSIDPSSAIPVYRGNQNGQSIDPSA**AIPIY**HGNQNGQSIDPSS
ALLIYRINQNGLSIDPSSAVPFYHGNKNGFSIDPSSAVPFYHGNK
NGLSVDPSSAIPVYQGNQNGQSIDPSSAVPVYRSNQNGQSIDPSS
AVPIYHGNKNGLSVDPSSAIFVYRGNQNGQSIDPSSAIPVYRGNQ
NGQSIDPSSAIPVYRGNQNGQSIDPSSALLIYRINQNGLSIDPSS
AVPFYHGKKNGFFVDPSSAVPFYHGNKNGLSVDPSSAIPVYQGNQ
NGQSIDPSSAVPVYRSNQNGQSIDPSSAVPIYHGNKNGLSVDPSS
AIFVYRGNQNGQSIDPSS**AIPIY**HGNQNGQSIDPSSAVPVYRGNQ
NGQSIDPSSALLIYRINQNGLSIDPSSAVPFYHGNQNGLSMDLSS
AVPFYRGIKNGLSVDPSSAVPFYYENKNGLSVDPSSAVPFYRGNQ
NGEQNSKANEKGATKNLSTDQ

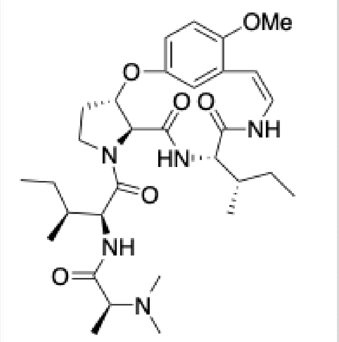


jubanine J I AIPIY

>KAH7517853.1 [Ziziphus jujuba var. spinosa]
MSINTESYTQPLHHSATTHLRTLPKKERIIASHFELFQIQLVIKMK
SFFALLAFSSLLLLSSTITARKEPAEYVKTVVEQVVKDLSVHPSS**A**
**VPFY**RSNKNGQSVDPSS**AVPFY**HENKNDLSVDPSS**AVPFY**RGNQND
QSVDPSS**AVPFY**HGNKNGLSVDPSSNKNGLSVDPSSALPFYRGNQN
GQSVDPSS**AVPFY**HGNKNGLSVDPSSVLPFYRGNQNGQSVDPSS**AV**
**PFY**HGNKNGLSVDPSSVLPFYRGNQNGQSVDPSS**AVPFY**HGNKNGL
SVDPSSVLPFYRGNQNGQSVDPSS**AVPFY**HGNKNGLSVDPSS**AVPF**
**Y**RGNQNGQSVDPSS**AVPFY**HGNKNGLSVDPSS**AVPFY**RGNQNGQSV
DPSS**AVPFY**HGNKNGFFVDPSS**AVPFY**HGNQNGQSVDPSS**AVPFY**H
GNKNGFFVDPSS**AVPFY**HGNQNGQSVDPSS**AVPFY**HGNKNGFFVDP
SSAIPFYYGNKNGLYIDPSS**AVPFY**HSNQNNQSVDPSS**AVPFY**RGN
QNGDQNSKANEEGATKNLPLTNEM**AVPFY**RGNQNGQFVDPSS**AVPF**
**Y**HGNKNSLSVDPSS**AVPFY**RSNQNGQSIDPSS**AVPFY**HR

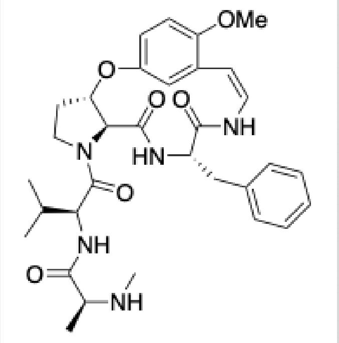


nummularine B I AVPFY

\>KAH7517853.1 [Ziziphus jujuba var. spinosa]
MSINTESYTQPLHHSATTHLRTLPKKERIIASHFELFQIQLVIKM
KSFFALLAFSSLLLLSSTITARKEPAEYVKTVVEQVVKDLSVHPS
S**AVPFY**RSNKNGQSVDPSS**AVPFY**HENKNDLSVDPSS**AVPFY**RGN
QNDQSVDPSS**AVPFY**HGNKNGLSVDPSSNKNGLSVDPSSALPFYR
GNQNGQSVDPSS**AVPFY**HGNKNGLSVDPSSVLPFYRGNQNGQSVD
PSS**AVPFY**HGNKNGLSVDPSSVLPFYRGNQNGQSVDPSS**AVPFY**H
GNKNGLSVDPSSVLPFYRGNQNGQSVDPSS**AVPFY**HGNKNGLSVD
PSS**AVPFY**RGNQNGQSVDPSS**AVPFY**HGNKNGLSVDPSS**AVPFY**R
GNQNGQSVDPSS**AVPFY**HGNKNGFFVDPSS**AVPFY**HGNQNGQSVD
PSS**AVPFY**HGNKNGFFVDPSS**AVPFY**HGNQNGQSVDPSS**AVPFY**H
GNKNGFFVDPSSAIPFYYGNKNGLYIDPSS**AVPFY**HSNQNNQSVD
PSS**AVPFY**RGNQNGDQNSKANEEGATKNLPLTNEM**AVPFY**RGNQN
GQFVDPSS**AVPFY**HGNKNSLSVDPSS**AVPFY**RSNQNGQSIDPSS**A**
**VPFY**HR

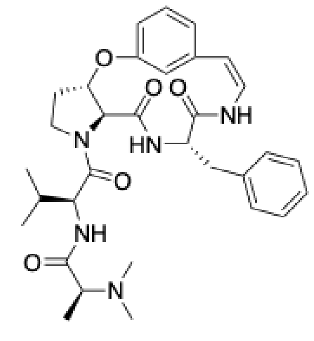

mauritine A I AVPFY

\>KAH7517853.1 [Ziziphus jujuba var. spinosa]
MSINTESYTQPLHHSATTHLRTLPKKERIIASHFELFQIQLVIKM
KSFFALLAFSSLLLLSSTITARKEPAEYVKTVVEQVVKDLSVHPS
S**AVPFY**RSNKNGQSVDPSS**AVPFY**HENKNDLSVDPSS**AVPFY**RGN
QNDQSVDPSS**AVPFY**HGNKNGLSVDPSSNKNGLSVDPSSALPFYR
GNQNGQSVDPSS**AVPFY**HGNKNGLSVDPSSVLPFYRGNQNGQSVD
PSS**AVPFY**HGNKNGLSVDPSSVLPFYRGNQNGQSVDPSS**AVPFY**H
GNKNGLSVDPSSVLPFYRGNQNGQSVDPSS**AVPFY**HGNKNGLSVD
PSS**AVPFY**RGNQNGQSVDPSS**AVPFY**HGNKNGLSVDPSS**AVPFY**R
GNQNGQSVDPSS**AVPFY**HGNKNGFFVDPSS**AVPFY**HGNQNGQSVD
PSS**AVPFY**HGNKNGFFVDPSS**AVPFY**HGNQNGQSVDPSS**AVPFY**H
GNKNGFFVDPSSAIPFYYGNKNGLYIDPSS**AVPFY**HSNQNNQSVD
PSS**AVPFY**RGNQNGDQNSKANEEGATKNLPLTNEM**AVPFY**RGNQN
GQFVDPSS**AVPFY**HGNKNSLSVDPSS**AVPFY**RSNQNGQSIDPSS**A**
**VPFY**HR

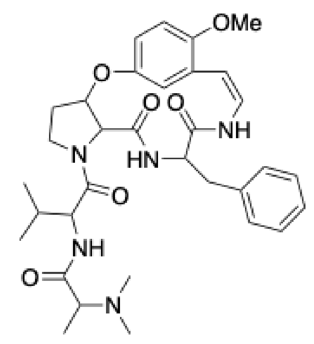

amphibine H I AVPFY

\>KAH7517848.1 [Ziziphus jujuba var. spinosa]
MKSFFALLAFSSLFLFSSTITARKEPVEYVKTVVDQVAKDLYVDPS
SHILLYHGKQNGKDAAKDQSVDPSS**IIPIY**HGNKNGLSIDPSS**IIP**
**IY**HENRNGQSVDPLSFIPIYLGKKNGLSIDASS**IIPIY**HGQKNGFS
VDPSS**IIPIY**GNHNAKDHSVDPSSLIPIYHGNQNGQSVDPSSLIPI
YRDNQNGQSVDPLSLVLLYRGNQQNGQSIDPSSLIPFYRGNQNGQS
VDPLSLIPFYRGNQNGQSVDPSSLIPFYRGNQNGQSVDPSSLIPFY
RGKQQNGQFVDPSSLIPFYRGNQNGQSVDPSSLIPIYRGNQNGDQN
FKANEEGVAKSVSTDQ

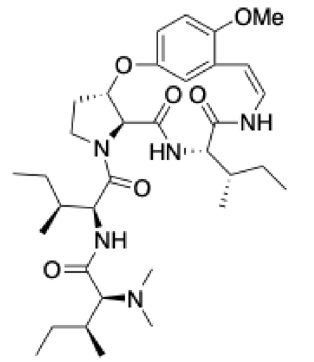

daechuine S3 I IIPIY

>XP_024932849.1 [Ziziphus jujuba]
MKSFFALLAFSSLLLLSSIVIARKEPVEYVKTVVEQVAKDLSIDP
SSR**WPIY**HGNQNGQSVDPLSR**WPIY**HGNQDQQFVDPLSR**WPIY**NG
NQNGQSVDPLSR**WPIY**HGNQNGQSVDPSSR**WPIY**HGNQNGQSVDP
SSR**WPIY**HGNQNGQSVDPSSR**WPIY**HGNRNGKSVDPSSW**WPIY**QN
GQSVDPSSW**WPIY**QNGQFVDPSSR**WPIY**HGNQNRQSVDPSSR**WPI**
**D**HGNQNGQSVDPSSR**WPIY**HGNQNGQSVDPSSK**WPIY**HGNQNGDQ
NSKANEEGAAKSASTDQ

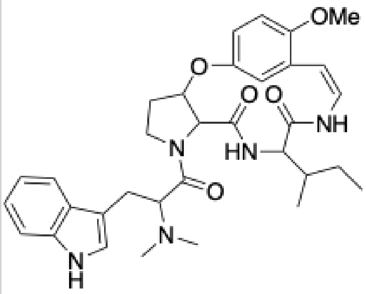

daechuine S10 | WPIY

>XP_024933341.1 [Ziziphus jujuba]
MKSFFALLAFSSLLLLSSIVIARKEPVEYVKTVVEQVAKDLSIDP
SSR**FPIY**HDNQNGQSVDTSSR**FPIY**HDNQNGQSIDPSSRFLIYHG
IDPFSR**FPIY**HGNQNGQSIDPSSR**FPIY**HGNQNGQSIDPSSQFLI
YRNGKSIDPSSR**FPIY**HGNQNGQSVDPSSR**FPIY**HDNQNGQSIDP
SSR**FPIY**HDNQNRQSIDPSSR**FPIY**HGNQNGQSIDPSSRFLIYHG
IDPSSR**FPIY**HGNQNGQSIDPSSR**FPIY**HGNQNGDQNSKANEEGA
AKSASTDQ

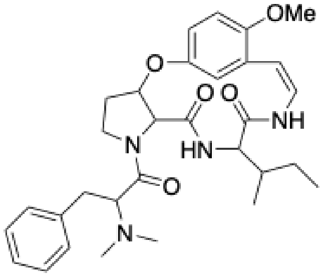

daechuine S6 | FPIY

>XP_024933341.1 [Ziziphus jujuba]
MKSFFALLAFSSLLLLSSIVIARKEPVEYVKTVVEQVAKDLSIDPS
SR**FPIY**HDNQNGQSVDTSSR**FPIY**HDNQNGQSIDPSSRFLIYHGID
PFSR**FPIY**HGNQNGQSIDPSSR**FPIY**HGNQNGQSIDPSSQFLIYRN
GKSIDPSSR**FPIY**HGNQNGQSVDPSSR**FPIY**HDNQNGQSIDPSSR**F**
**PIY**HDNQNRQSIDPSSR**FPIY**HGNQNGQSIDPSSRFLIYHGIDPSS
R**FPIY**HGNQNGQSIDPSSR**FPIY**HGNQNGDQNSKANEEGAAKSAST
DQ

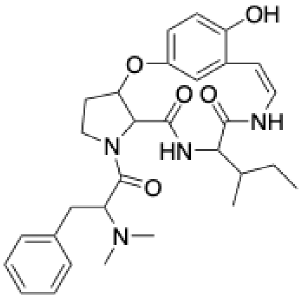

daechuine S26 | FPIY

>KAH7517857.1 [Ziziphus jujuba var. spinosa]
MKSFFALLAFSSLLLLSSTITARKEPGEYVKTVVEQVAEDLFVDP
SSIIPFYHKNKNGQSVDPLS**FLPIY**HGNQNGQSIDPSSLLLIYHG
NQIGQSVDPSS**FLPIY**HSNQNGQSVDPSS**FLPIY**HGNRNGQSVDP
SS**FLPIY**HGNRNGQSVDPSS**FLPIY**HGNRNGQSVDPSS**FLPIY**HG
NRNGQSVDPSS**FLPIY**HGNQNGQSVDPSS**FLPIY**QGNRNGQSVDP
SS**FLPIY**QGNRNGQSVDPSS**FLPIY**HGNLNRQSVDPSS**FLPIY**HG
NLNGQSVDPSS**FLPIY**HGNQNGQSVDPSSFHLLYHGNRNGQSVDP
SS**FLPIY**HGNLNGQSIDPSSRNGQSVDPSSLLLIYRDNQNGEQNP
KANEEGVAKSISTDQ

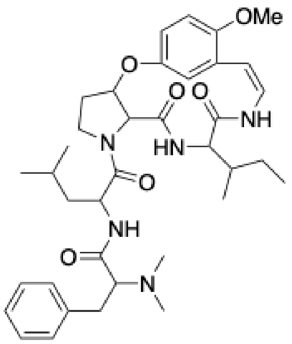


mucronine D I FLPIY

>KAH7517857.1 [Ziziphus jujuba var. spinosa]
MKSFFALLAFSSLLLLSSTITARKEPGEYVKTVVEQVAEDLFVDP
SSIIPFYHKNKNGQSVDPLS**FLPIY**HGNQNGQSIDPSSLLLIYHG
NQIGQSVDPSS**FLPIY**HSNQNGQSVDPSS**FLPIY**HGNRNGQSVDP
SS**FLPIY**HGNRNGQSVDPSS**FLPIY**HGNRNGQSVDPSS**FLPIY**HG
NRNGQSVDPSS**FLPIY**HGNQNGQSVDPSS**FLPIY**QGNRNGQSVDP
SS**FLPIY**QGNRNGQSVDPSS**FLPIY**HGNLNRQSVDPSS**FLPIY**HG
NLNGQSVDPSS**FLPIY**HGNQNGQSVDPSSFHLLYHGNRNGQSVDP
SS**FLPIY**HGNLNGQSIDPSSRNGQSVDPSSLLLIYRDNQNGEQNP
KANEEGVAKSISTDQ

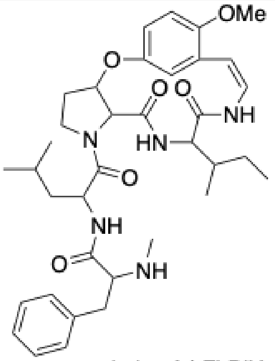


nummularine A I FLPIY

>KAH7517848.1 [Ziziphus jujuba var. spinosa]
MKSFFALLAFSSLFLFSSTITARKEPVEYVKTVVDQVAKDLYVDPS
SHILLYHGKQNGKDAAKDQSVDPSSIIPIYHGNKNGLSIDPSSIIP
IYHENRNGQSVDPLSFIPIYLGKKNGLSIDASSIIPIYHGQKNGFS
VDPSSIIPIYGNHNAKDHSVDPSS**LIPIY**HGNQNGQSVDPSS**LIPI**
**Y**RDNQNGQSVDPLSLVLLYRGNQQNGQSIDPSSLIPFYRGNQNGQS
VDPLSLIPFYRGNQNGQSVDPSSLIPFYRGNQNGQSVDPSSLIPFY
RGKQQNGQFVDPSSLIPFYRGNQNGQSVDPSS**LIPIY**RGNQNGDQN
FKANEEGVAKSVSTDQ

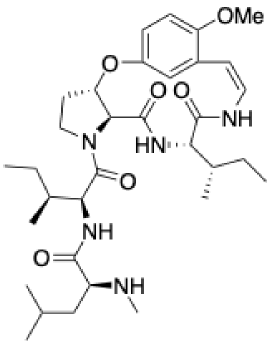


paliurine H I LIPIY

**Figure S14.** Putative core sequences from the precursor peptides map to 15 known cyclopeptide alkaloids from *Ziziphus jujuba*.

```
XP_015887038.1   -----------------------------------------------MKTFFLLMIIFTF
XP_015887040.1   -----------------------------------------------MKTFFLLMIIFTF
KAH7517853.1     M---SINTESYTQPLHHSATTHLRTLPKKERIIASHFELFQIQLVIKMKSFF---ALLAF
KAH7517857.1     -----------------------------------------------MKSFF---ALLAF
XP_024932826.1   -----------------------------------------------MKSFF---ALLAF
XP_015892608.1   -----------------------------------------------MKSFF---ALLAF
XP_024933342.1   -----------------------------------------------MKSFF---ALLAF
XP_024933341.1   -----------------------------------------------MKSFF---ALLAF
XP_024932849.1   -----------------------------------------------MKSFF---ALLAF
XP_015880721.1   M----------REPLHHSS-------------------FSKFQFVIKMKSFF---AVLAF
KAH7517848.1     -----------------------------------------------MKSFF---ALLAF
XP_015892671.1   -----------------------------------------------MKSFF---ALLAF
KAH7517850.1     --------------------------------------MNLLDIILEHKGVG---EKLAR
KAH7533561.1     MYGVSINTESYERA-------------------SSSFELLLIPICHQDEKFLC-ASRLLF
KAH7533587.1     MYGVSINTESYERA-------------------SSSFELLQIPICHQDEKFLC-GSRLLF
                                                                 : .      :

XP_015887038.1   SSL-------LLFSNTITARKDPGEYWTVIMENQPIPKAIQGLLGNIANSRNGGQFAKEF
XP_015887040.1   SSL-------LLVPNTITARKDPGEYWAVIMENQPIPKAIQGLLGNIANSRNGGKFAKDF
KAH7517853.1     SSL-------LLLSSTITARKEPAEYVKTVVE----------------------QVVKDL
KAH7517857.1     SSL-------LLLSSTITARKEPGEYVKTVVE----------------------QVAEDL
XP_024932826.1   SSL-------LLLSSTITARKEPGEYVKTVVE----------------------QVAEDL
XP_015892608.1   SSL-------LLLSSIVIARKEPVEYVKTVVE----------------------QVAKDL
XP_024933342.1   SSL-------LLLSSIVIARKEPVEYVKTVVE----------------------QVAKDL
XP_024933341.1   SSL-------LLLSSIVIARKEPVEYVKTVVE----------------------QVAKDL
XP_024932849.1   SSL-------LLLSSIVIARKEPVEYVKTVVE----------------------QVAKDL
XP_015880721.1   SSL-------LLLASTITARKEPAEYVKTALK----------------------QIAKDL
KAH7517848.1     SSL-------FLFSSTITARKEPVEYVKTVVD----------------------QVAKDL
XP_015892671.1   SSL-------FLFSSTITARKEPVEYVKTGVD----------------------QVAKDL
KAH7517850.1     SCSGVVLVPPFALSSTIIARKEPVEYVKTVVE----------------------QVAKDL
KAH7533561.1     TSSG----KDQKLASTITARKEPAEYVKTTLE----------------------QVAKDL
KAH7533587.1     TSSG----KDQKLASTITARKEPAEYVKTALK----------------------QIAKDL
                 :.          ... : ***:* **  . :.                      :..:::

XP_015887038.1   SAETSSNLWTY
XP_015887040.1   SAETSSNFWTY
KAH7517853.1     SVHPSSAVPFY
KAH7517857.1     FVDPSSIIPFY
XP_024932826.1   FVDPSSIIPFY
XP_015892608.1   SIDPSSRFPIY
XP_024933342.1   SIDPSSRFPIY
XP_024933341.1   SIDPSSRFPIY
XP_024932849.1   SIDPSSRWPIY
XP_015880721.1   SIDPSSFVPLY
KAH7517848.1     YVDPSSHILLY
XP_015892671.1   YVDPSSHILLY
KAH7517850.1     SVDPSFAVPFY
KAH7533561.1     SIDPSSFVPFY
KAH7533587.1     SIDPSSFVPLY
                     ..*      *
```

**Figure S15.** Putative precursor peptides from *Ziziphus* have an N-terminal signal peptide (yellow) as predicted by SignalP 6.0.[15] A core-less leader peptide-like region (un-highlighted) is found immediately after the signal peptide. Core sequences are highlighted in red.

```
KAH7517851.1    MQ------------LYSIG-----------------------------------------
QXY82431.1      MAQSLILLFLIVMGCYDVGAIERVHSAKEGVSEAMNDQGHPTIANKAVPASVEDPSQADA
                *              *.:*
KAH7517851.1    ------------------------------------------------------------
QXY82431.1      DNILLYPSYAAKKAVPEDPSQADADNVLFYRSYAAKKAVPASVEDPSHDADNVLFYPSYV

KAH7517851.1    ------------------------------------------------------------
QXY82431.1      AKKAVPASVEDPSQADADNFLLYPYAAKKAVPASVEDPSHDADNVLFYPSYVAKKAVPAS

KAH7517851.1    ------------------------------------------------------------
QXY82431.1      VEDPSQADADNFLFYPYAAKKAVPASVEDPSHDADNVLFYPSYVAKKAVPASVEDPSHDA

KAH7517851.1    ---------------------------------------------------NGYSQHHNH
QXY82431.1      DNVLFYPSYVAKKAVPASVEDPSQADADNVLFYPSYAARPKAPSTMDHDATHNAAQNMHH
                                                                   :. :*: :*
KAH7517851.1    QQDDGDDPLDGARKVGFFTVDDLYVGKVVGIQFYTRDPSSLPPFLSKEVANHFLFSIK—
QXY82431.1      QKG----------ALLFFRMNTLKVGGEVVI----------PSLASHALGGRKLMTPRLE
                *:.           : ** :: * **  * *          *.: *: :..: *:: :
KAH7517851.1    ------ELLLILQLFGMALGSLEAQLAERTLRFCAEEPTKGEQKTCATSIESFIEFASSM
QXY82431.1      AMLGSMELPRLLSVLKIPLDSTLARKSKTNLGNCRSPPLQGEKKACVSSIRSMTNFARSV
                      **  :*.:: :.*.*  *: :: .*  * . * :**:*:*.:**.*: :** *:
KAH7517851.1    LGGGQYGVDFRSIKTTHLGKPVSIYQNYTFLDIKEVHSPIVVAFHIMDYPYIVYMCHSQT
QXY82431.1      LE------DKKPLEHLNPSRPVSSPEHFKIMDVKVVTENSVVTCHPMVFPYALYMCHFVP
                *       * :.::  : .:***  :::.::*:* * .  **: * * :** :****  .
KAH7517851.1    SRVYQIKIVGQNVGDALNAF-VICHIDTSQWAPDHISFKLLSVKPGTILICHFFGPHNPV
QXY82431.1      KSV-PIKVTLQDDHDKLVVVPVMCHMDTSEFDPSHLSFKILNTKPGEAEMCHWM-PNSHI
                . *  **:. *:  * * .. *:**:***:: *.*:***:*..***   :**:: *:. :
KAH7517851.1    -------------
QXY82431.1      MWYTSDGKTRDVL
```

**Figure S16.** Amino acid sequence alignment of a split BURP-domain protein from *Ziziphus jujuba* (KAH7517851.1) and the fused BURP-domain responsible for the biosynthesis of selanine A and B from *Selaginella kraussiana* (QXY82431.1).

```
>> XP_003588556.2  BURP domain protein RD22 [Medicago truncatula]
   #    score  bias  c-Evalue  i-Evalue hmmfrom  hmm to    alifrom  ali to    envfrom  env to     acc
 ---   ------ ----- --------- --------- ------- -------    ------- -------    ------- -------    ----
   1 !   29.9   0.0   5.1e-08   0.00036       8     105 ..      21     127 ..      15     297 .. 0.80

  Alignments for each domain:
  == domain 1  score: 29.9 bits;  conditional E-value: 5.1e-08
    cpaAlignment   8 kllkkkMkslfaflvlfslllfantieARkdpgeYWksvmkdqpmpeaIkgllhqdsssskekadchtss.........ekkeksfvkdfeprpn 93
                     +l   +M+ +++ ++ f++l ++ t  A   p+ YW+s+  + pmp+a  +llh     skeka    +           +   ++ +d ++ p  
  XP_003588556.2  21 QLHLSNMEFHLFHIITFLMLVLVGTNAAMLPPQLYWQSMLPNSPMPKAFTNLLHPAGYWSKEKARDASNGgldvgvrkgYEGGGTYLNDEKIIPL 115
                     556689**999999999999999*******************************99775666666333334777776664456777777777777 PP

    cpaAlignment  94 vsiYhndvkkke 105
                     + +Y+  + ++e
  XP_003588556.2 116 IYFYPIPIPLNE 127
                     777766655555 PP
```

**Figure S17**. Our custom HMM identified fused BURP-domains due to conservation of both the signal peptide and N-terminal sequence. An example from XP_003588556.2 is shown.

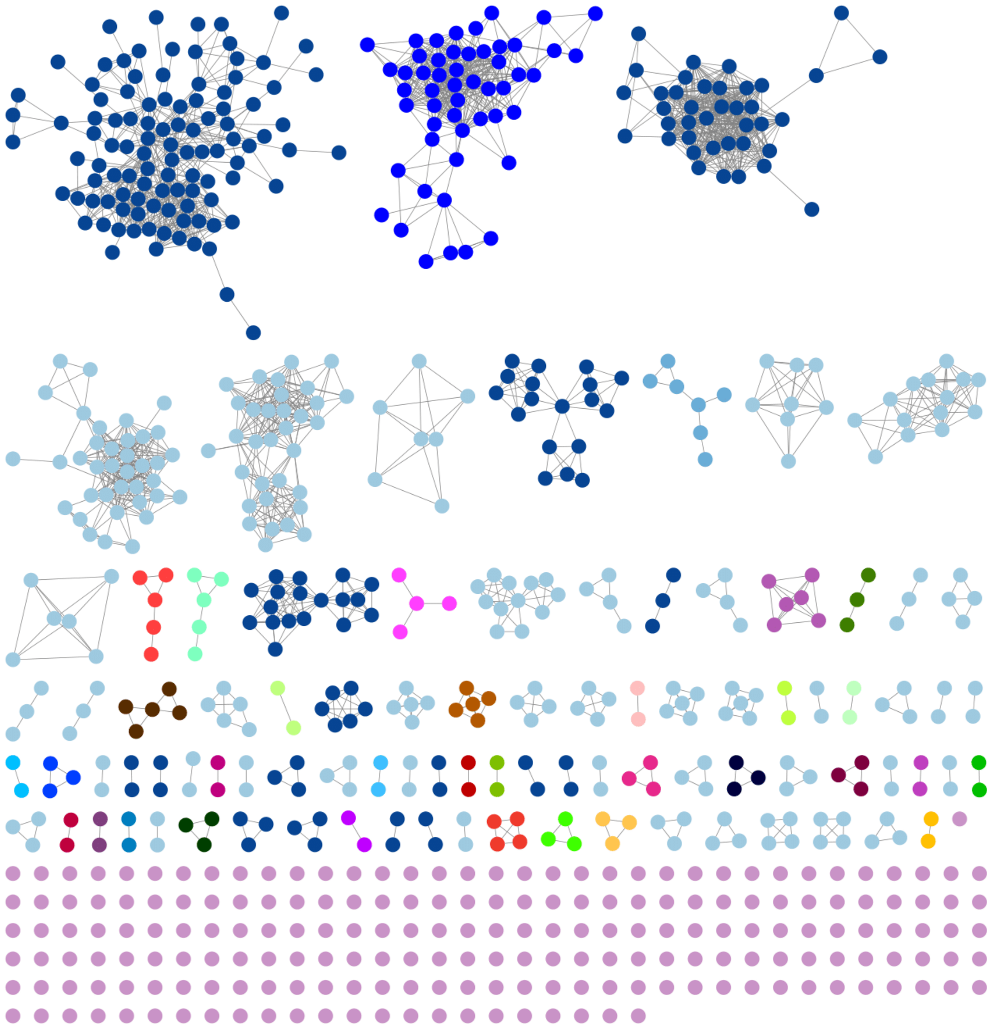

**Figure S18**. A second SSN (SSN2) was generated using the singlets from SSN1. An alignment score of 20 was used. Navy blue clusters contain serine or threonine at the second position of the core. Light blue clusters contain isoleucine at the second and third positions. A cluster of *C. americanus* sequences are seen as a pink triplet.

>XP_039019285.1 [Hibiscus syriacus]
MKSTLVFFAFLCILLFTKTIAARKAPGEECNDETKEKFQTETTKEAKQQY
VIN**QIPLFY**YPKNVDGQEMDNAGKELAINQIPLLYYPKKDDGQEMDKARK
ESVINQIPLLYYPKKVDGQEMDNAGKESDIN**QIPLFY**YPKKVDGQEMDNA
GRESVIN**QIPLFY**YPKKVDGQEMDNAGKEFVIN**QIPLFY**YPKKVDGQKMD
KAGKESVINQIPLLYYPKKVDGQEMDNARKESVINQIPLLYYPKKVDGQK
MDNAGKESDIN**QIPLFY**YPKKVDGQEMDNAGKESDIN**QIPLFY**YPKKVDG
QQMDNAIKESVINQVPLVYYPKKVGSA

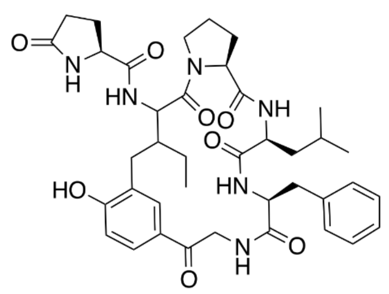

hibispeptin A | QIPLFY

>XP_039019285.1 [Hibiscus syriacus]
MKSTLVFFAFLCILLFTKTIAARKAPGEECNDETKEKFQTETTKEAKQQY
VINQIPLFYYPKNVDGQEMDNAGKELAIN**QIPLLY**YPKKDDGQEMDKARK
ESVIN**QIPLLY**YPKKVDGQEMDNAGKESDINQIPLFYYPKKVDGQEMDNA
GRESVINQIPLFYYPKKVDGQEMDNAGKEFVINQIPLFYYPKKVDGQKMD
KAGKESVIN**QIPLLY**YPKKVDGQEMDNARKESVIN**QIPLLY**YPKKVDGQK
MDNAGKESDINQIPLFYYPKKVDGQEMDNAGKESDINQIPLFYYPKKVDG
QQMDNAIKESVINQVPLVYYPKKVGSA

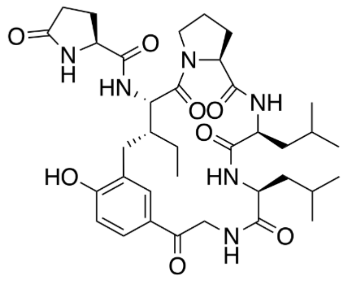

hibispeptin B | QIPLLY

>XP_039019285.1 [Hibiscus syriacus]
MKSTLVFFAFLCILLFTKTIAARKAPGEECNDETKEKFQTETTKEAKQQY
VINQIPLFYYPKNVDGQEMDNAGKELAINQIPLLYYPKKDDGQEMDKARK
ESVINQIPLLYYPKKVDGQEMDNAGKESDINQIPLFYYPKKVDGQEMDNA
GRESVINQIPLFYYPKKVDGQEMDNAGKEFVINQIPLFYYPKKVDGQKMD
KAGKESVINQIPLLYYPKKVDGQEMDNARKESVINQIPLLYYPKKVDGQK
MDNAGKESDINQIPLFYYPKKVDGQEMDNAGKESDINQIPLFYYPKKVDG
QQMDNAIKESVIN**QVPLVY**YPKKVGSA

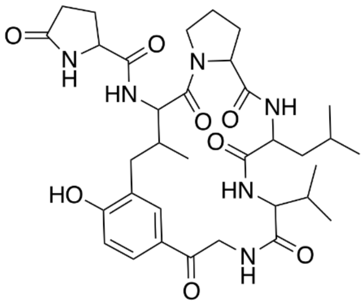

HIS669 | QVPLVY

**Figure S19**. Putative core sequences from the precursor peptides map to hibispeptin A and B cyclopeptides from *H. syriacus*. Core sequences corresponding to Q-V-P-L-V-Y (HIS669) are also shown.

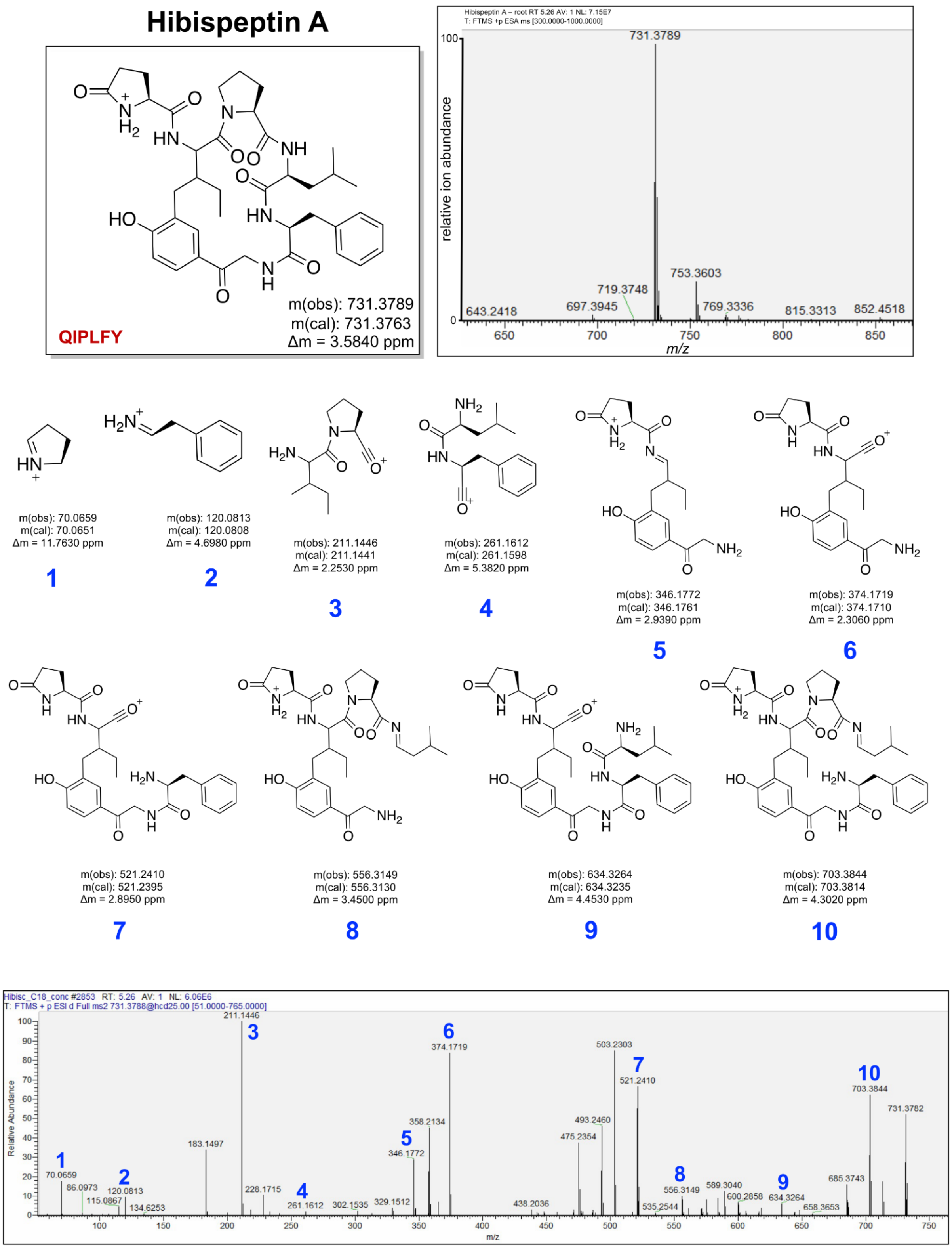


**Figure S20.** MS/MS spectrum of hibispeptin A derived from methanol extraction of dry *H. syriacus* roots.

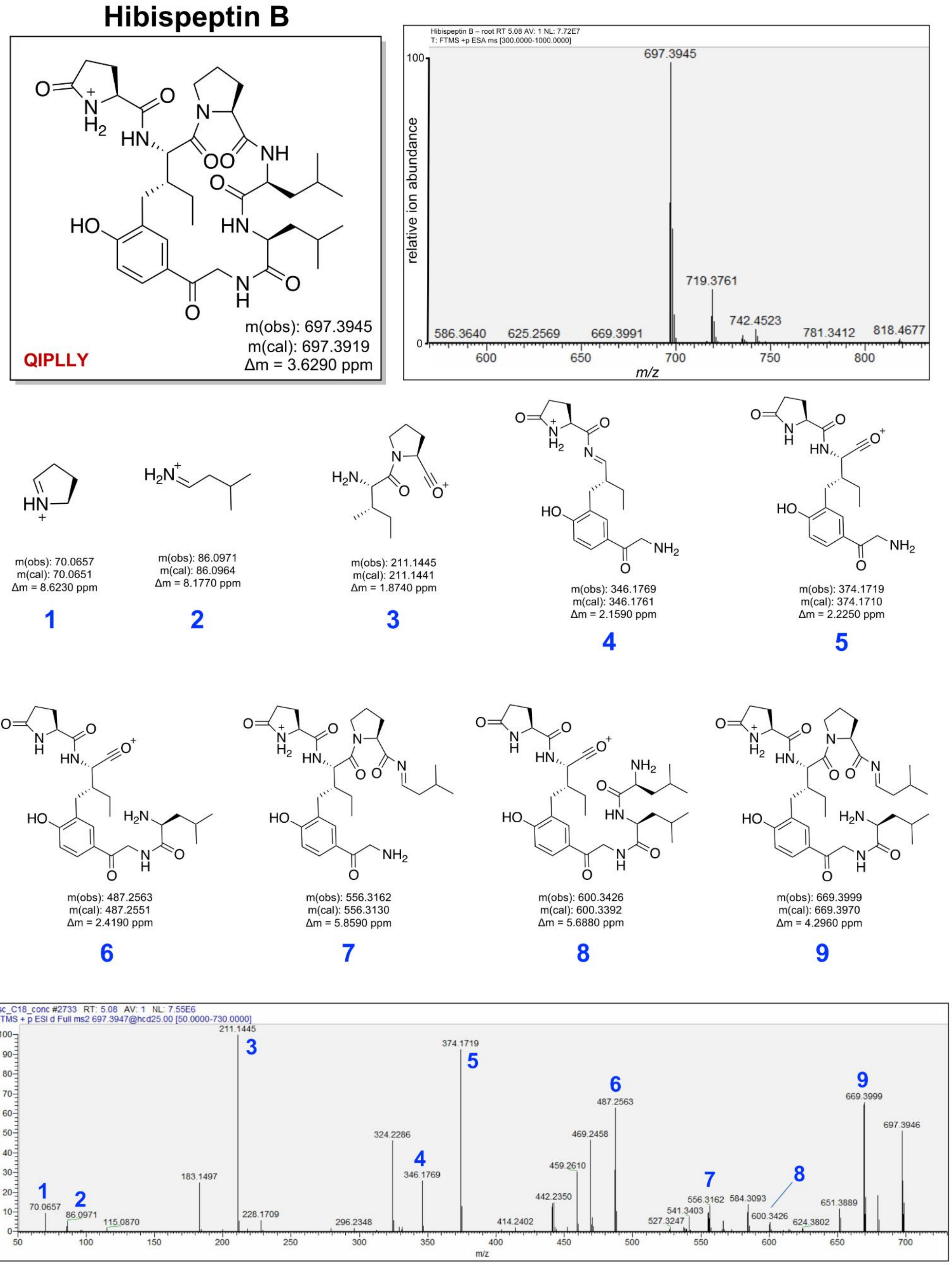


**Figure S21.** MS/MS spectrum of hibispeptin B derived from methanol extraction of dry *H. syriacus* roots.

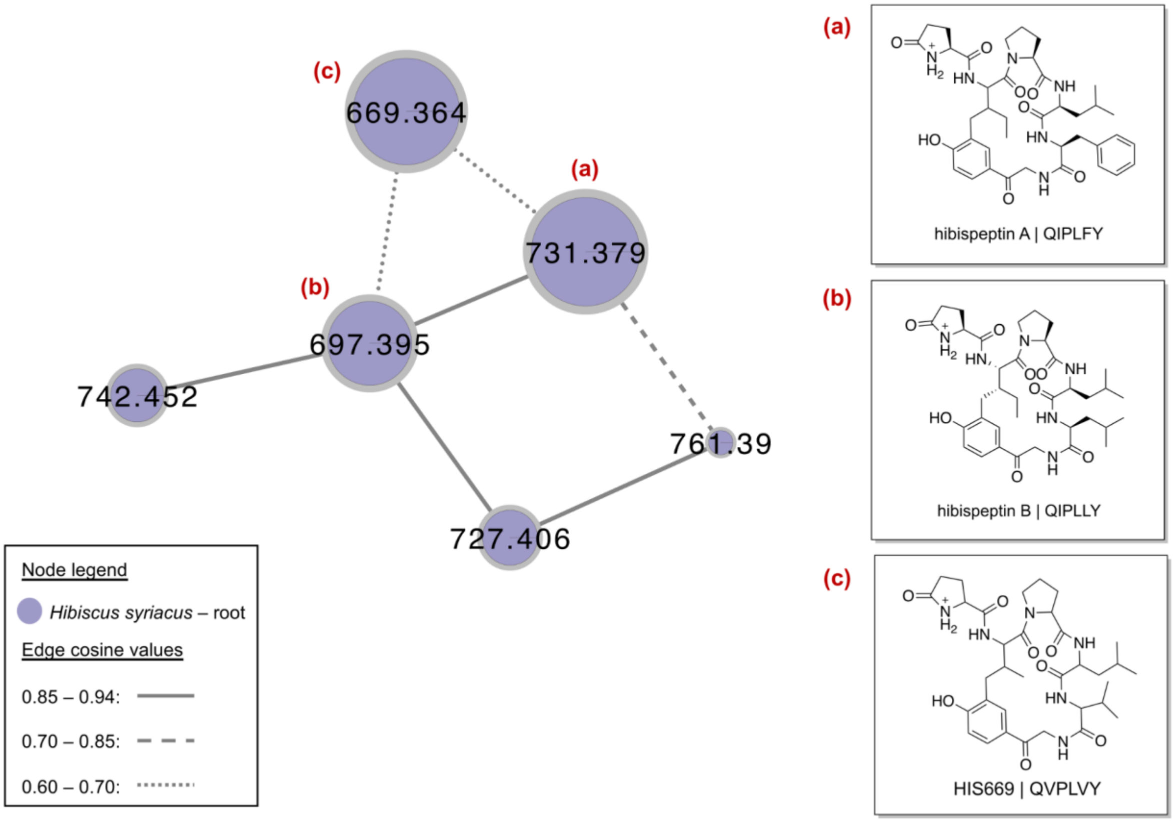


**Figure S22.** *Hibiscus syriacus* cyclopeptide-containing spectral cluster of hibispeptin A, B, and HIS669 LC-MS/MS datasets of ground plant roots. Node sizes represent numbers of spectra. Node values are precursor ion mass values.

### HIS669

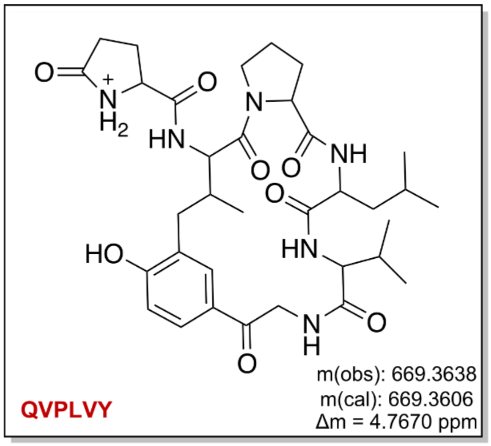


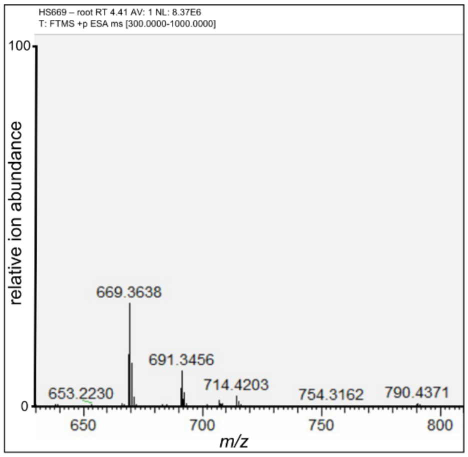


**Proposed structure for *H. syriacus* feature 669**

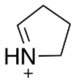

m(obs): 70.0658
m(cal): 70.0651
Δm = 10.1930 ppm

**1**

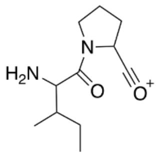

m(obs): 211.1446
m(cal): 211.1441
Δm = 2.3950 ppm

**2**

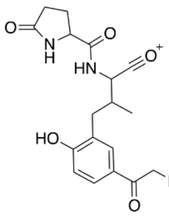

m(obs): 360.1562
m(cal): 360.1554
Δm = 2.2840 ppm

**3**

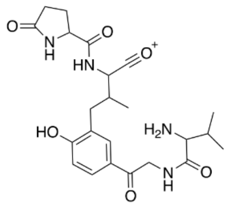

m(obs): 459.2250
m(cal): 459.2238
Δm = 2.6110 ppm

**4**

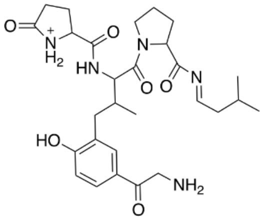

m(obs): 542.2973
m(cal): 542.2973
Δm = -0.0190 ppm

**5**

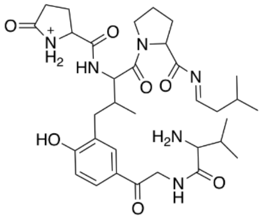

m(obs): 641.3686
m(cal): 641.3657
Δm = 4.4680 ppm

**6**

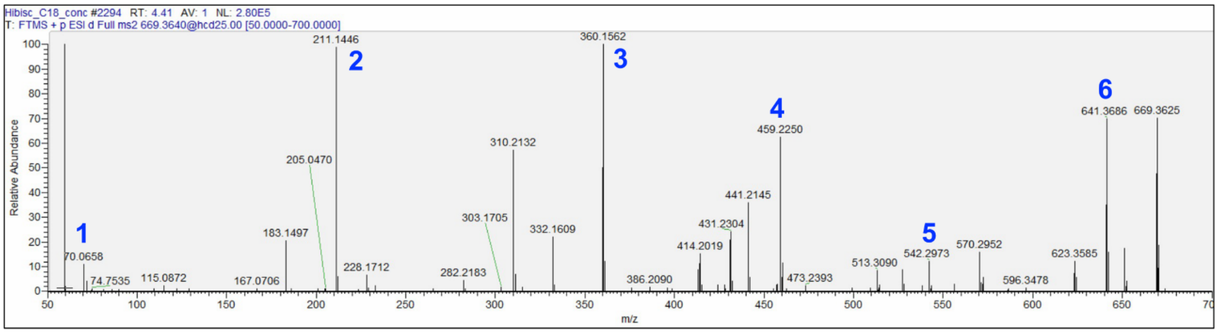


**Figure S23.** MS/MS spectrum of HIS669 derived from methanol extraction of dry *H. syriacus* dry roots.

>XP_027065604.1 [Coffea arabica]
MASSITLIAVFSIALFACITEARKNPTDFLQSAVINEHTEDNHHA
ESSLSNQKKTSNGNTLKDFESKPGS**FLWGY**QGNDAESKSKEEKPL
MKGFESKPGS**FLWGY**QGNDAESKSKEEKPLMKGFESKPGS**FLWGY**
QGNDVESKSKEEKPLTKDFESKPGS**FLWGY**QGNHAESKSKKEKPL
MKDFESKPGS**FLWGY**QGNHAEYKEKKPLVKDN

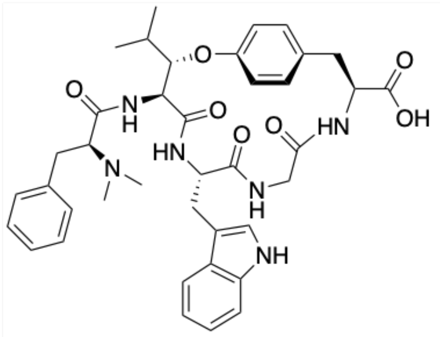

arabipeptin A | FLWGY

>XP_027086167.1 [Coffea arabica]
MRSSVALVAFFSIALLACFTEARKDPRGILRPAASPGAFTEQNEH
LGSNTLNEFESKPGSILHADEPRSILPYHGRDANSKEEKPQMKDF
ESKSESVLLFYGGDKANLQEAKPYMKDFESKPESVLILYRGDKAN
LQEAKPHMKDFESKPESVLLFYGGDKANLQEAKPYMKDFESKPES
**FLILY**GGDKANLQEAKPHMKDFESKPESVLPFYGGDKANLQEAKP
YMKDFESKPES**FLILY**GGDKANLQEGKLHI

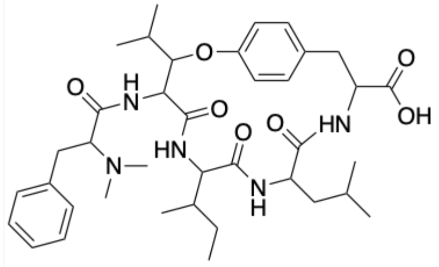

CAR694 | FLILY

>XP_027086167.1 [Coffea arabica]
MRSSVALVAFFSIALLACFTEARKDPRGILRPAASPGAFTEQNEH
LGSNTLNEFESKPGSILHADEPRSILPYHGRDANSKEEKPQMKDF
ESKSES**VLLFY**GGDKANLQEAKPYMKDFESKPESVLILYRGDKAN
LQEAKPHMKDFESKPES**VLLFY**GGDKANLQEAKPYMKDFESKPES
FLILYGGDKANLQEAKPHMKDFESKPESVLPFYGGDKANLQEAKP
YMKDFESKPESFLILYGGDKANLQEGKLHI

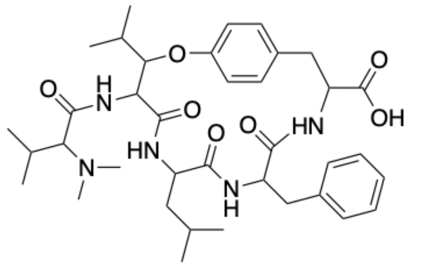

CAR680 | VLLFY

>XP_027086167.1 [Coffea arabica]
MRSSVALVAFFSIALLACFTEARKDPRGILRPAASPGAFTEQNEH
LGSNTLNEFESKPGSILHADEPRSILPYHGRDANSKEEKPQMKDF
ESKSESVLLFYGGDKANLQEAKPYMKDFESKPES**VLILY**RGDKAN
LQEAKPHMKDFESKPESVLLFYGGDKANLQEAKPYMKDFESKPES
FLILYGGDKANLQEAKPHMKDFESKPESVLPFYGGDKANLQEAKP
YMKDFESKPESFLILYGGDKANLQEGKLHI

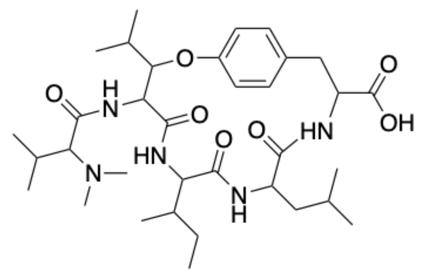

CAR646 | VLILY

>XP_027072356.1 [Coffea arabica]
MASSITLIAVFSIALFACITEARKNPTDSLQSAVINEHTEDNHHA
ELSLSNQKKTSDGNTLKDFESKPGSLLWNYQGNHAESKSKEEKPL
MKDFESKLGS**LLLY**HYQGNHAESKSKEEKPLMKDFESKLG

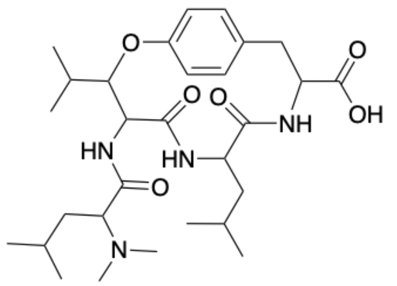

CAR547 | LLLY

**Figure S24.** Putative core sequences from the *C. arabica* precursor peptides map to arabipeptin A and other mass spectrometric features observed in the GNPS network.

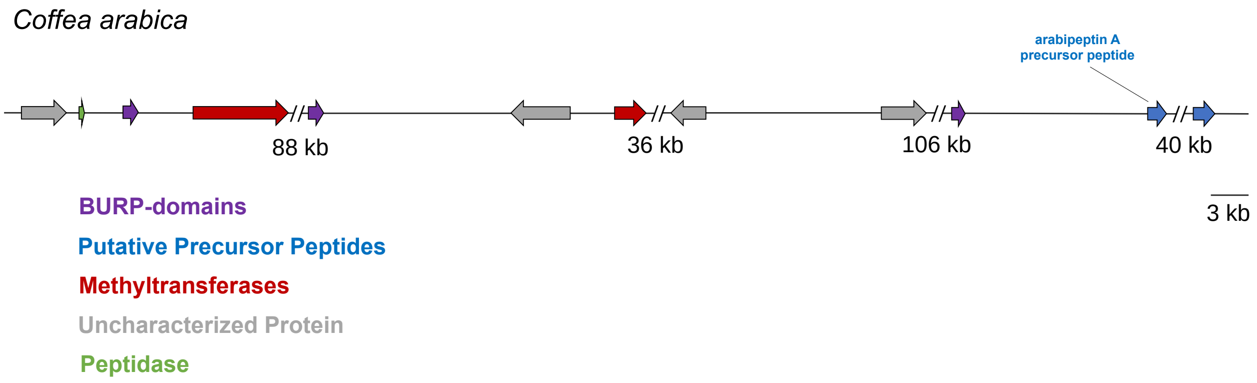


**Figure S25.** Genomic organization of the arabipeptin A precursor peptides from *C. arabica* (NCBI accession: XP_027066141.1).

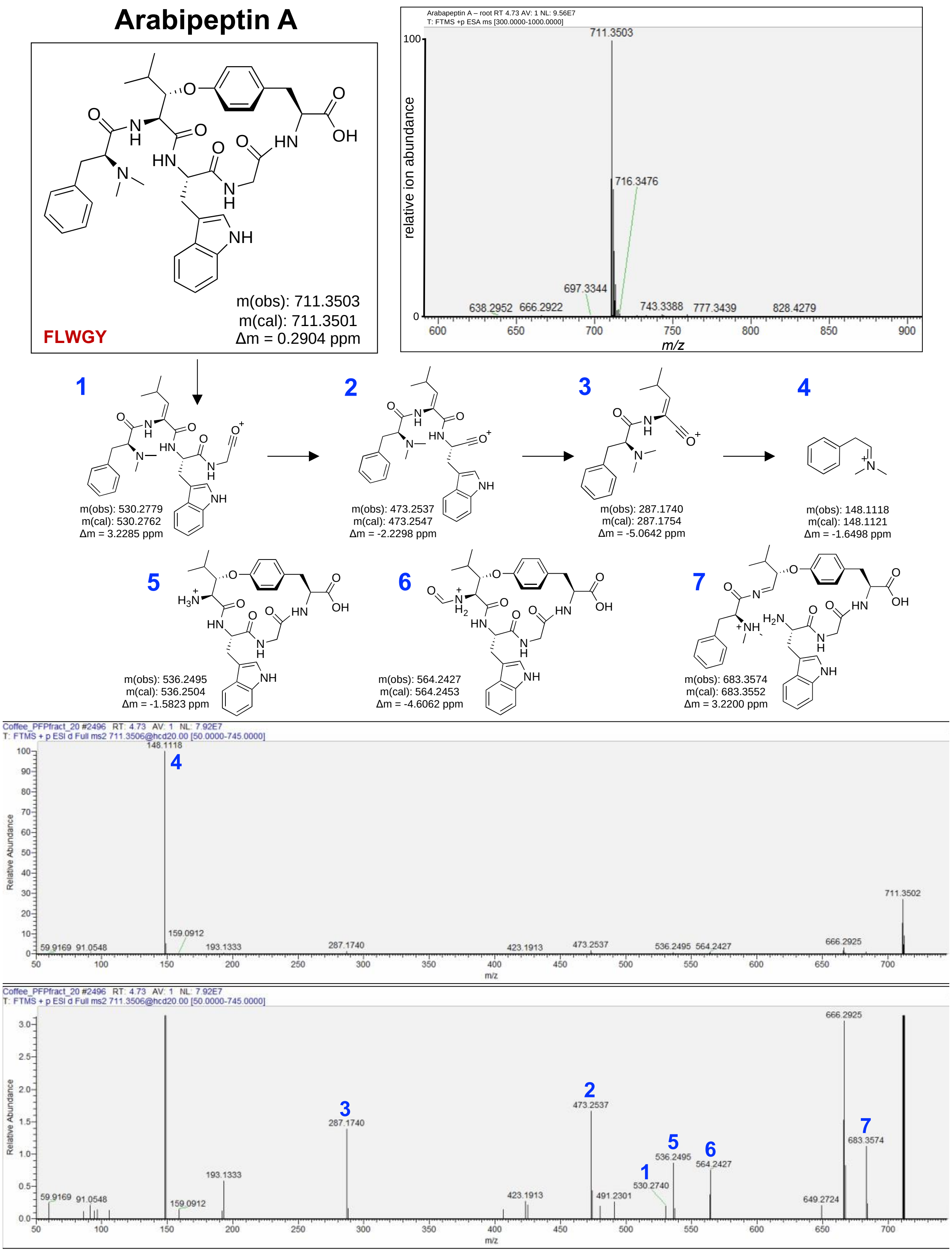


**Figure S26.** MS/MS spectrum of arabipeptin A derived from 1-butanol extraction of *C. arabica* dry roots.

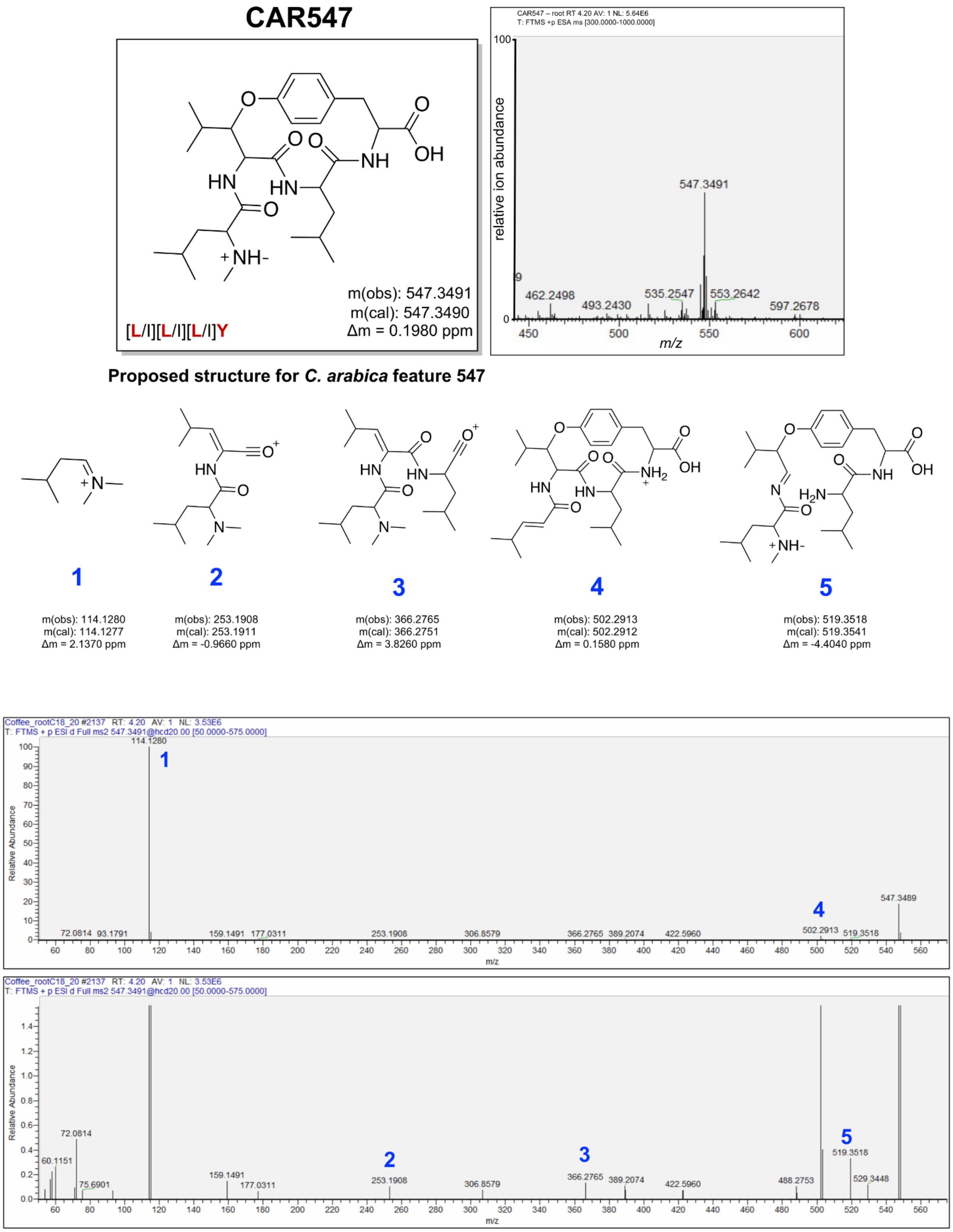


**Figure S27.** MS/MS spectrum of CAR547 derived from methanol extraction of dry *C. arabica* roots. Amino acids indicated in the brackets show the alternative structure by replacing L with I, or I with L. The red amino acid is observed in the precursor peptide.

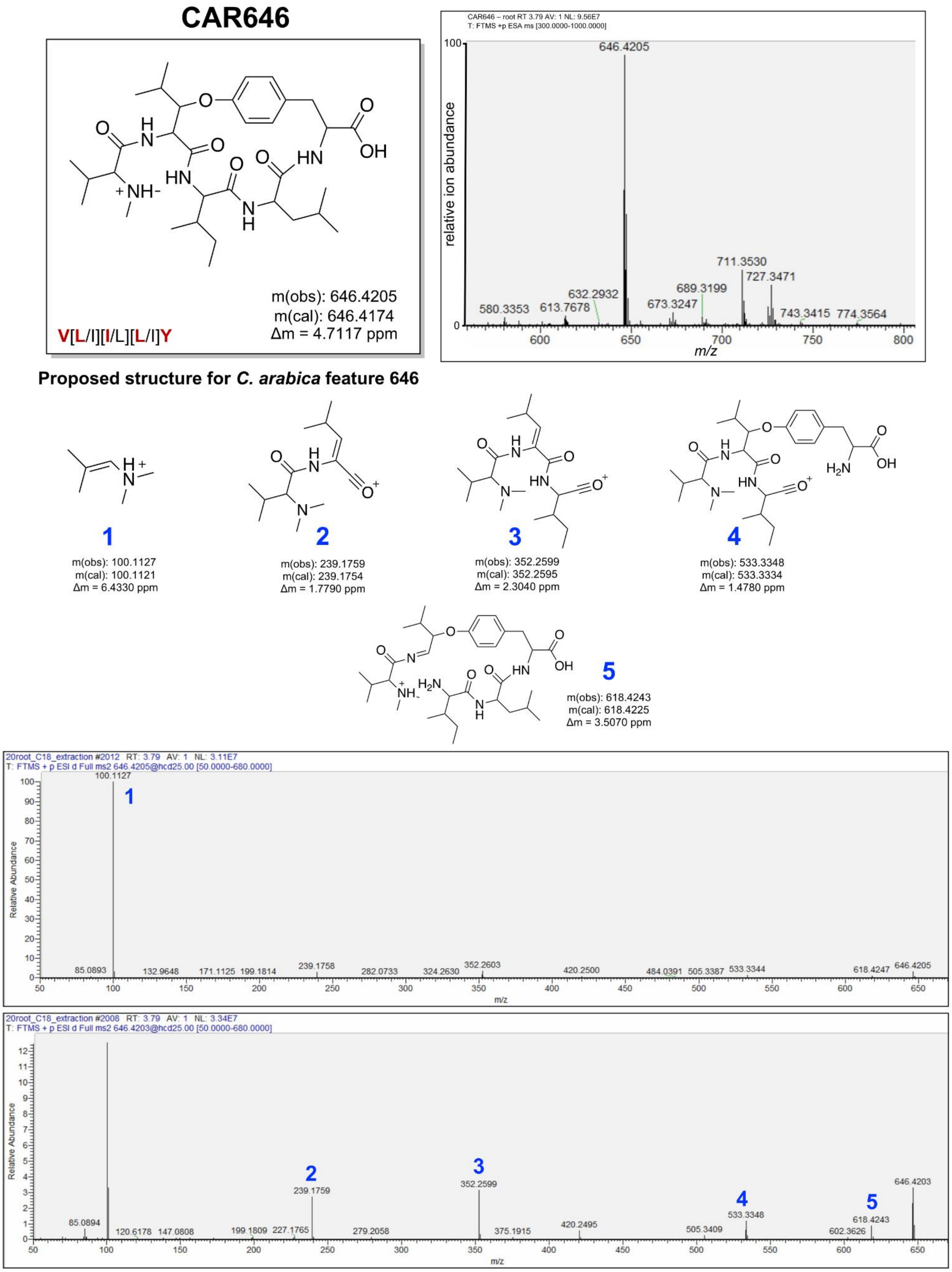


**Figure S28.** MS/MS spectrum of CAR646 derived from methanol extraction of dry *C. arabica* roots. Amino acids indicated in the brackets show the alternative structure by replacing L with I, or I with L. The red amino acid is observed in the precursor peptide.

### CAR680

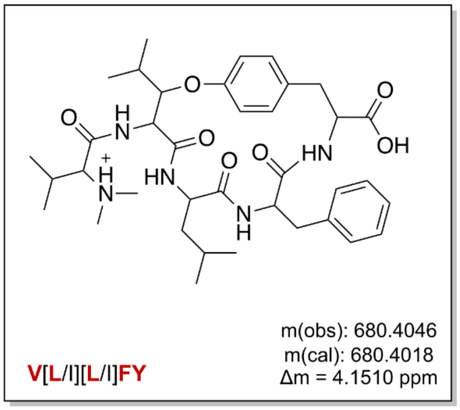


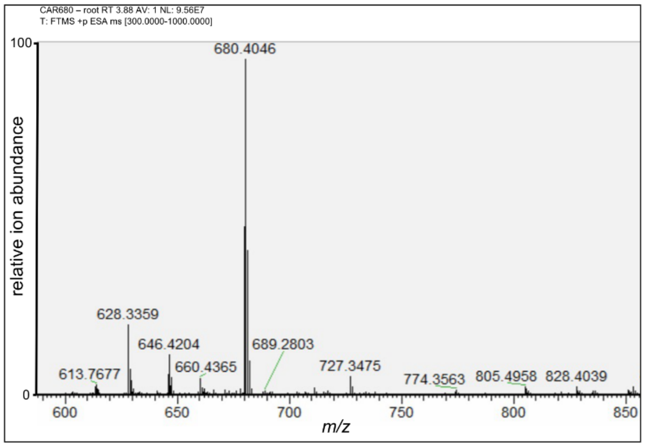


**Proposed structure for *C. arabica* feature 680**

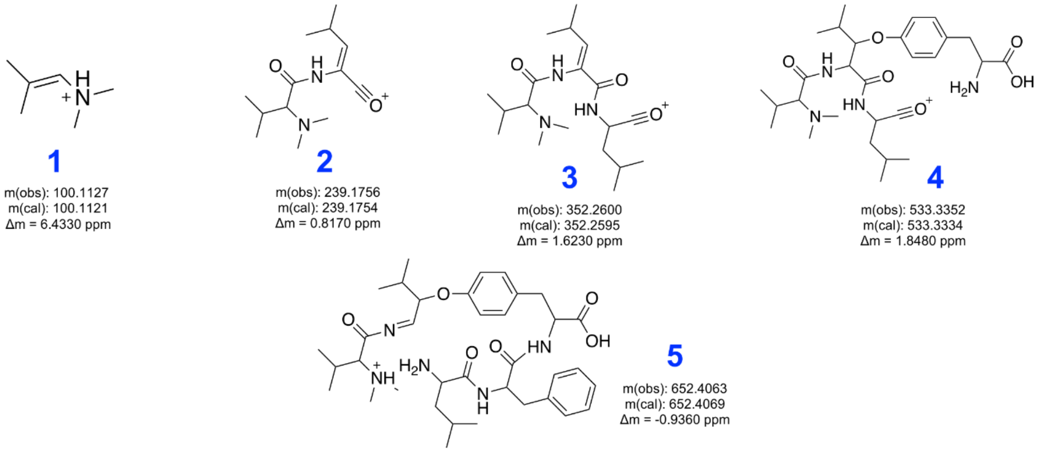


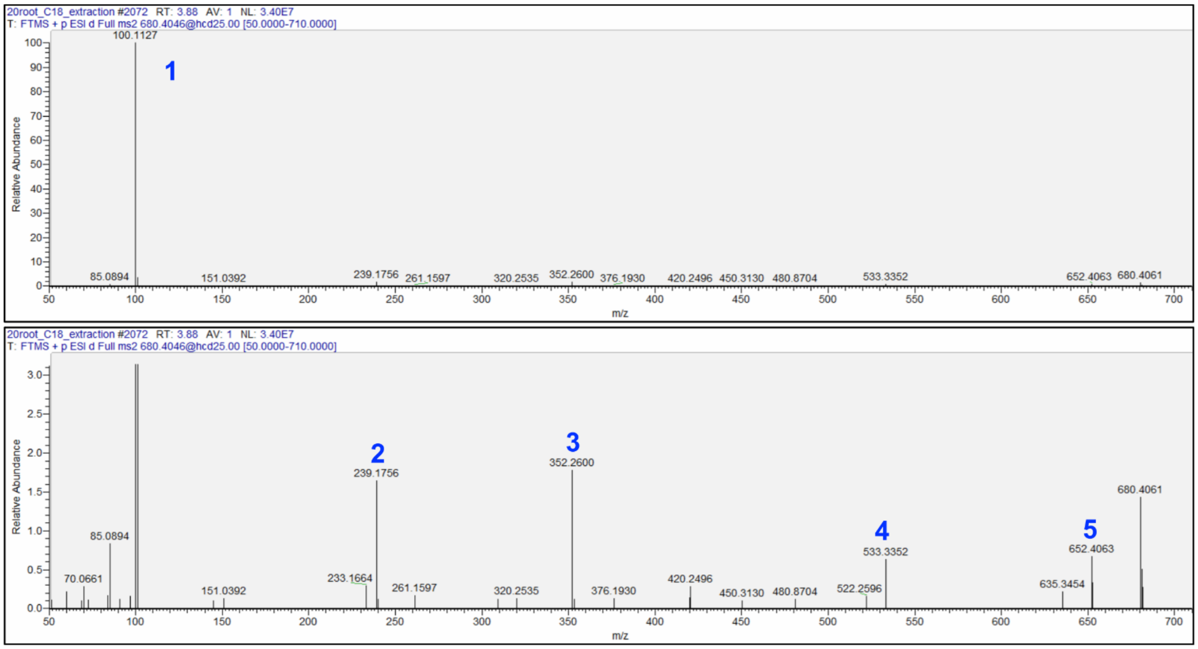


**Figure S29.** MS/MS spectrum of CAR680 derived from methanol extraction of dry *C. arabica* roots. Amino acids indicated in the brackets show the alternative structure by replacing L with I, or I with L. The red amino acid is observed in the precursor peptide.

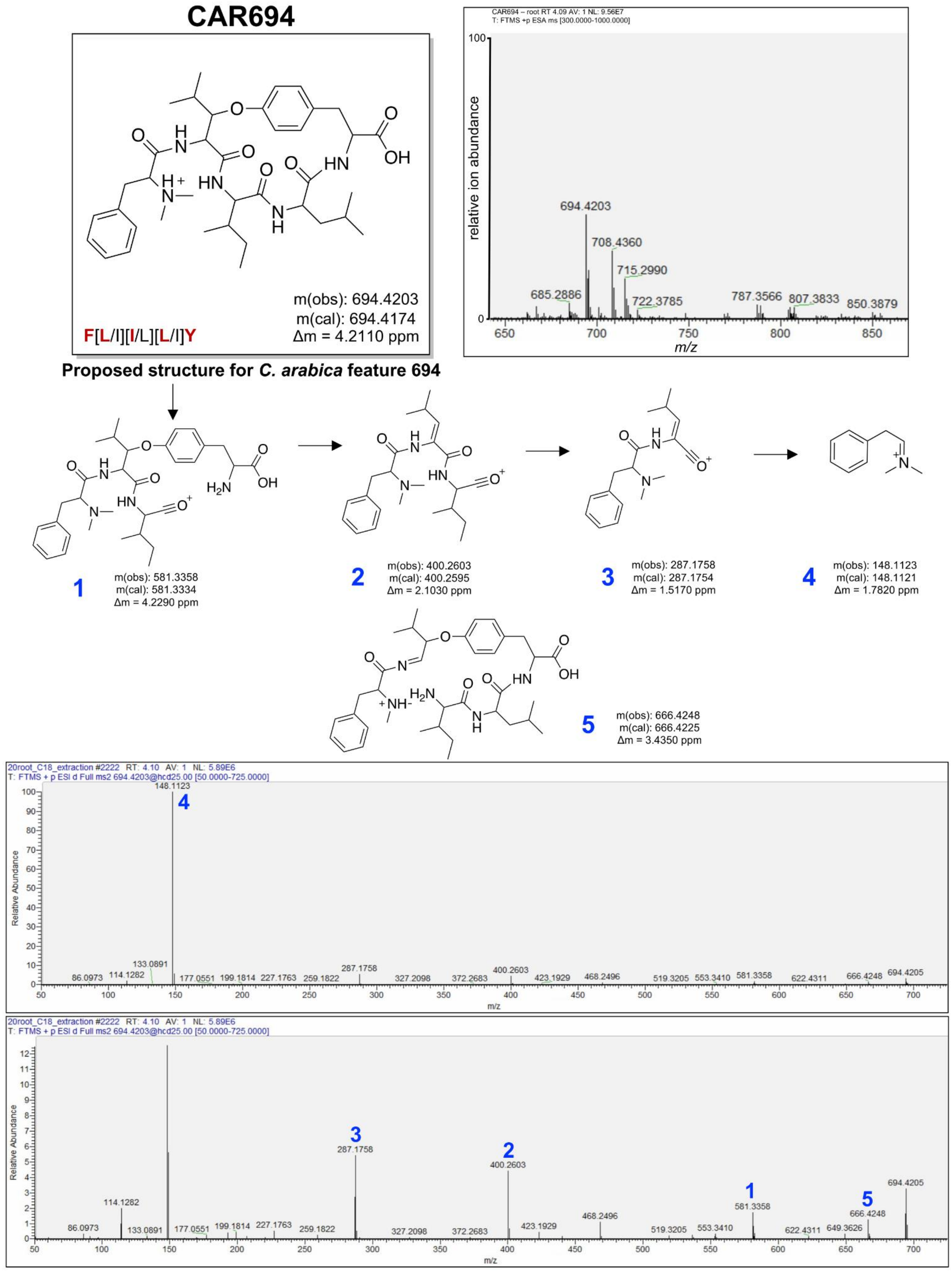


**Figure S30.** MS/MS spectrum of CAR694 derived from methanol extraction of dry *C. arabica* roots. Amino acids indicated in the brackets show the alternative structure by replacing L with I, or I with L. The red amino acid is observed in the precursor peptide.

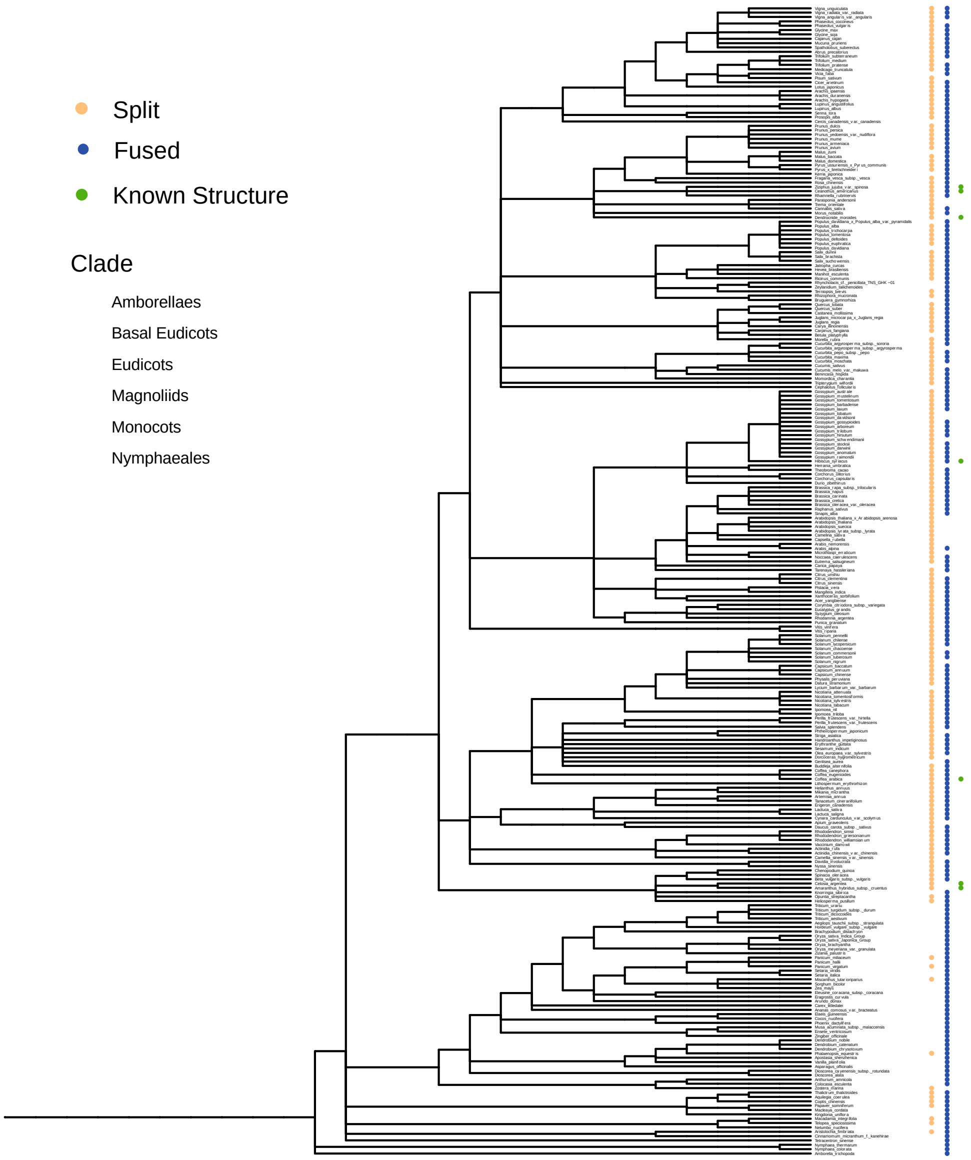


**Figure S31.** Cladogram of Viridiplantae species. Blue and gold dots correspond to the presence of fused BURP-domains and stand alone precursor peptides, respectively. As every genome is not present or fully annotated, the lack of a dot does not suggest an absence in the organism. Green dots indicate the presence of known structures derived from a stand alone precursor peptide. Clades are highlighted: amborellales (orange), basal eudicots (purple), eudicots (blue), magnoliids (pink), monocots (yellow), nymphaeales (gray).

### NMR Analysis

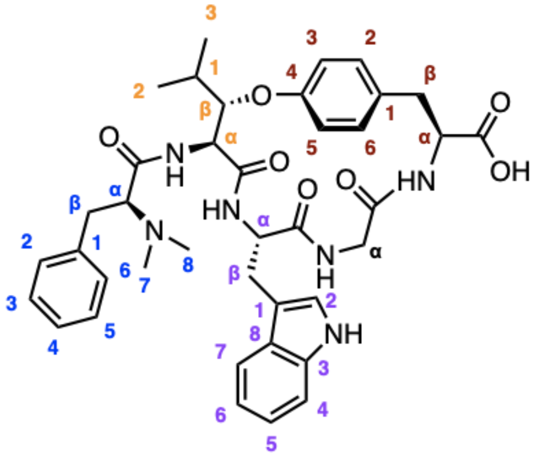

**arabipeptin A | FLWGY**

Numbering scheme for isolated arabipeptin A.

NMR correlations summary

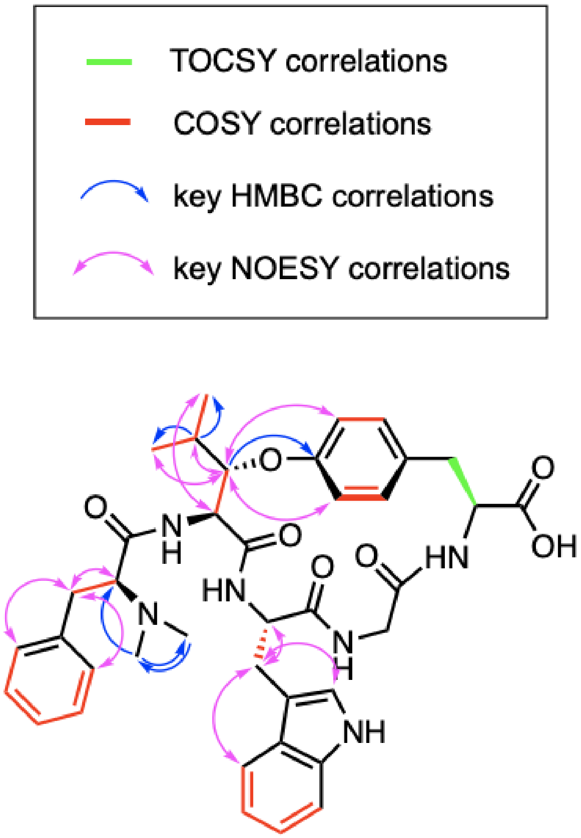

### $^1$H NMR table for arabipeptin A

| Residue | C | δ($^{13}$C) [ppm] | H | δ($^1$H) (*J*, d) [ppm] | HMBC |
|---|---|---|---|---|---|
| Phe[1] | Cα | 70.2 | Hα | 3.43 (5.2, 9.1, dd) | $C7_{Phe}$, $C8_{Phe}$ |
| | Cβ | 35.2 | Hα, Hβ | 3.07 (9.2, 13.7, dd); 2.98 (m) | $C\alpha_{Phe}$, $C2_{Phe}$, $C6_{Phe}$ |
| | C1 | 137.8 | - | - | |
| | C2 | 128.4 | H2 | 7.20 (4.4, d) overlap | $C\beta_{Phe}$ |
| | C3 | 129.0 | H3 | 7.20 (4.4, d) overlap | |
| | C4 | 127.6 | H4 | 7.11 (m) | |
| | C5 | 129.0 | H5 | 7.20 (4.4, d) overlap | |
| | C6 | 128.4 | H6 | 7.20 (4.4, d) overlap | $C\beta_{Phe}$ |
| | C7 (N-$CH_3$) | 41.5 | Hα, Hβ, Hγ | 2.43 (s) | $C8_{Phe}$, $C\alpha_{Phe}$ |
| | C8 (N-$CH_3$) | 41.5 | Hα, Hβ, Hγ | 2.43 (s) | $C7_{Phe}$, $C\alpha_{Phe}$ |
| Leu[2] | Cα | 54.5 | Hα | 4.62 (10.1, d) | $C\beta_{Leu}$ |
| | Cβ | 80.4 | Hβ | 4.42 (2.0, 10.1, dd) | $C4_{Tyr}$, $C2_{Leu}$ |
| | C1 | 29.5 | H1 | 1.97 (6.3, p) | |
| | C2 | 13.4 | Hα, Hβ, Hγ | 0.98 (6.8, d) | $C1_{Leu}$, $C3_{Leu}$, $C\beta_{Leu}$ |
| | C3 | 19.6 | Hα, Hβ, Hγ | 0.92 (6.8, d) | $C1_{Leu}$, $C2_{Leu}$, $C\beta_{Leu}$ |
| Trp[3] | Cα | 54.5 | Hα | 4.28 (6.8, t) | |
| | Cβ | 30.3 | Hα, Hβ | 2.98 (m); 2.98 (m) | $C\alpha_{Trp}$, $C\beta_{Trp}$, $C2_{Trp}$ |
| | C1 | 109.5 | - | - | |
| | C2 | 123.9 | H2 | 7.01 (s) | $C8_{Trp}$ |
| | C3 | 127.8 | - | - | |
| | C4 | 111.1 | H4 | 7.26 (8.0, d) | |
| | C5 | 121.5 | H5 | 7.03 (7.5, t) | $C7_{Trp}$ |
| | C6 | 118.6 | H6 | 6.97 (8.0, 7.0, dd) | |
| | C7 | 118.4 | H7 | 7.47 (7.9, d) | $C8_{Trp}$ |
| | C8 | 137.9 | - | - | |
| Gly[4] | Cα | 42.4 | Hα, Hβ | 3.77 (17.0, d); 3.00 (12.6, d) | |
| Tyr[5] | Cα | 54.8 | Hα | 4.33 (m) | |
| | Cβ | 36.1 | Hα, Hβ | 2.83 (7.1, dd); 3.15 (m) | |
| | C1 | 138.1 | - | - | |
| | C2 | 129.3 | H2 | 6.87 (8.5, d)* | |
| | C3 | 111.1 | H3 | 6.61 (8.0, d)* | |
| | C4 | 157.8 | - | - | |
| | C5 | 118.8 | H5 | 6.73 (8.4, d)* | |
| | C6 | 130.4 | H6 | 6.82 (8.5, d)* | $C4_{Tyr}$ |

*Interchangeable hydrogens

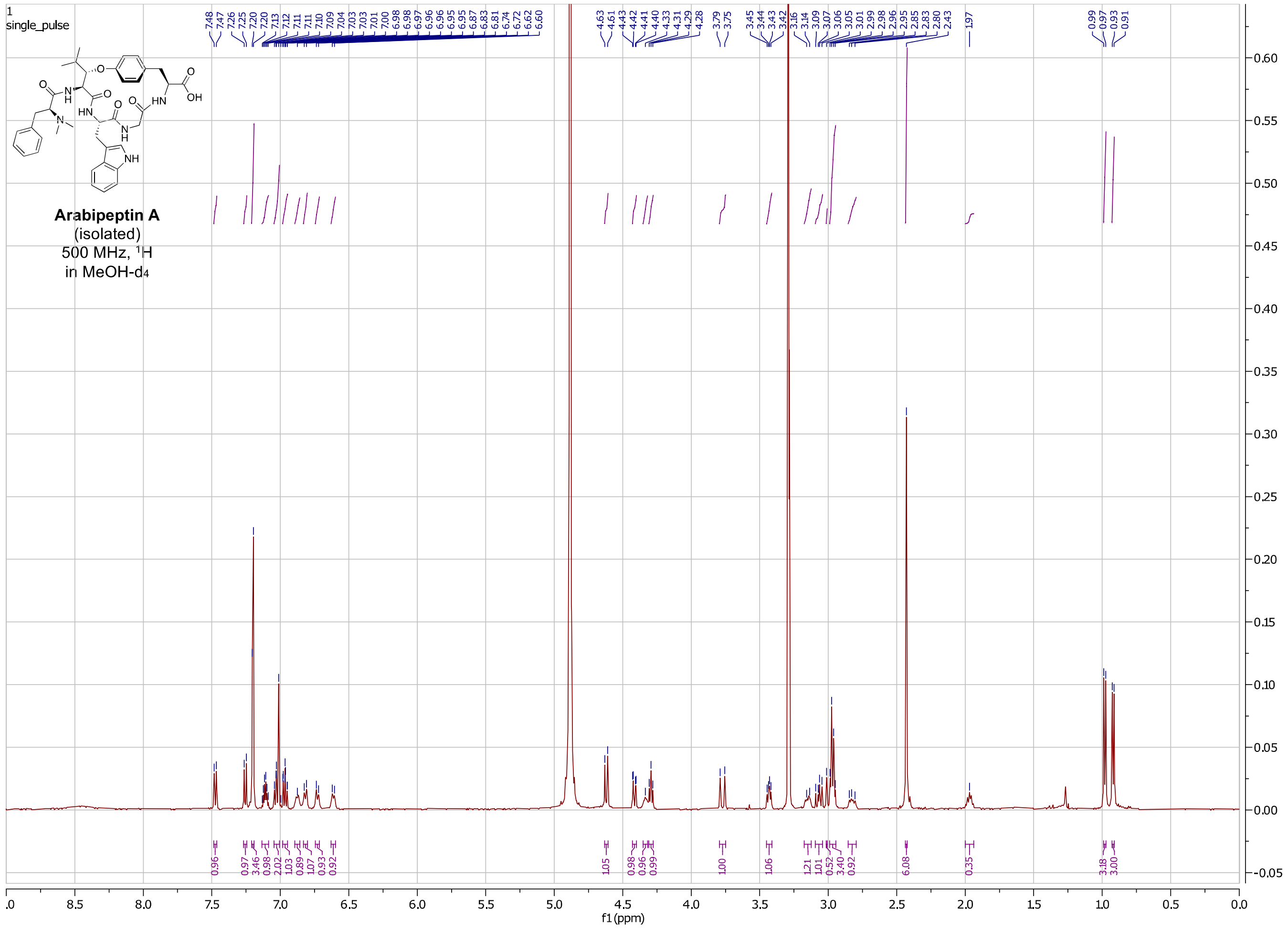


**Figure S32.** $^{1}$H-NMR spectrum of arabipeptin A in MeOH-$d_4$ (500 MHz).

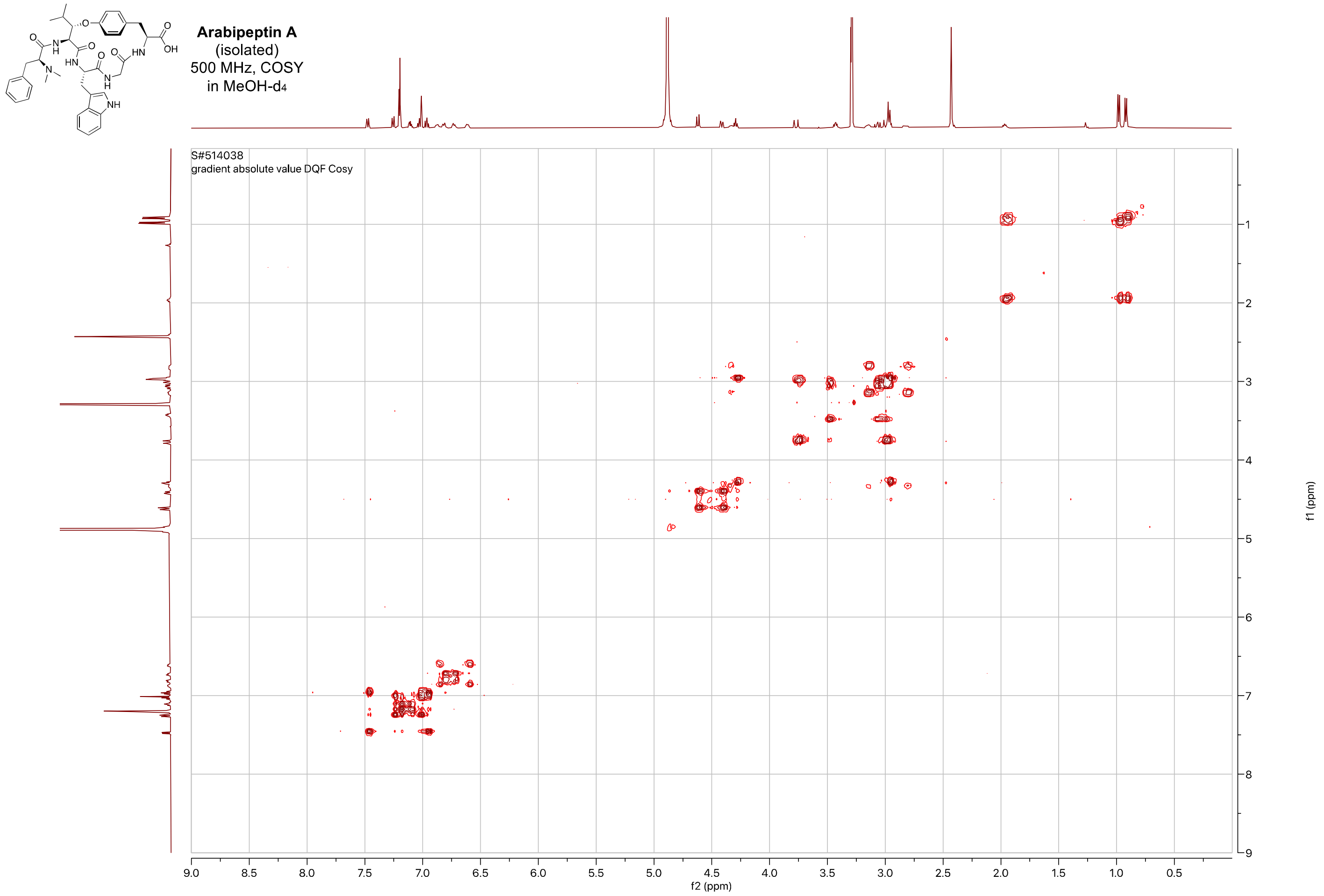


**Figure S33.** [1]H-[1]H-COSY NMR spectrum of arabipeptin A in MeOH-$d_4$ (500 MHz).

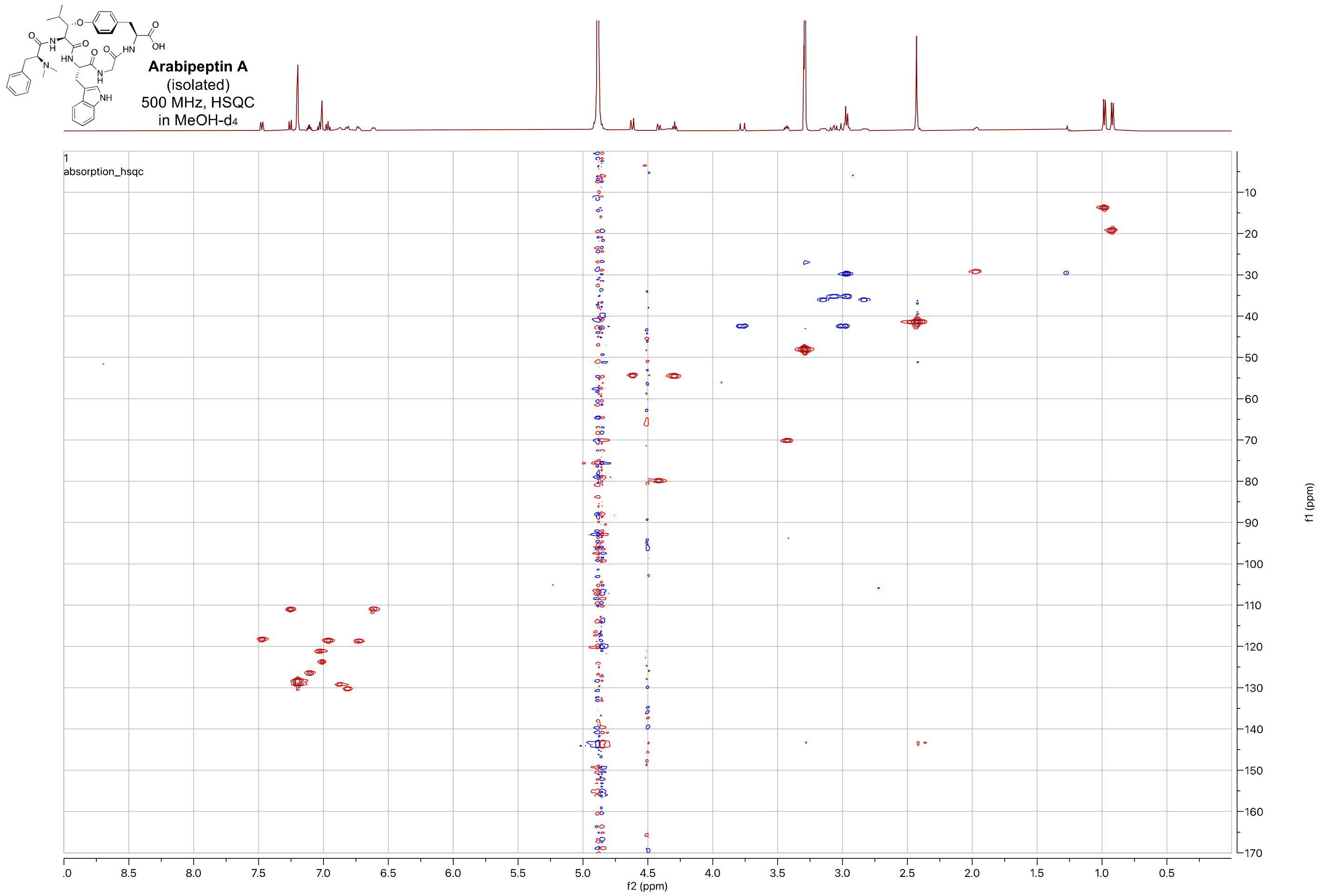


**Figure S34.** $^{1}H$-$^{13}C$-HSQC NMR spectrum of arabipeptin A in MeOH-$d_4$ (500 MHz).

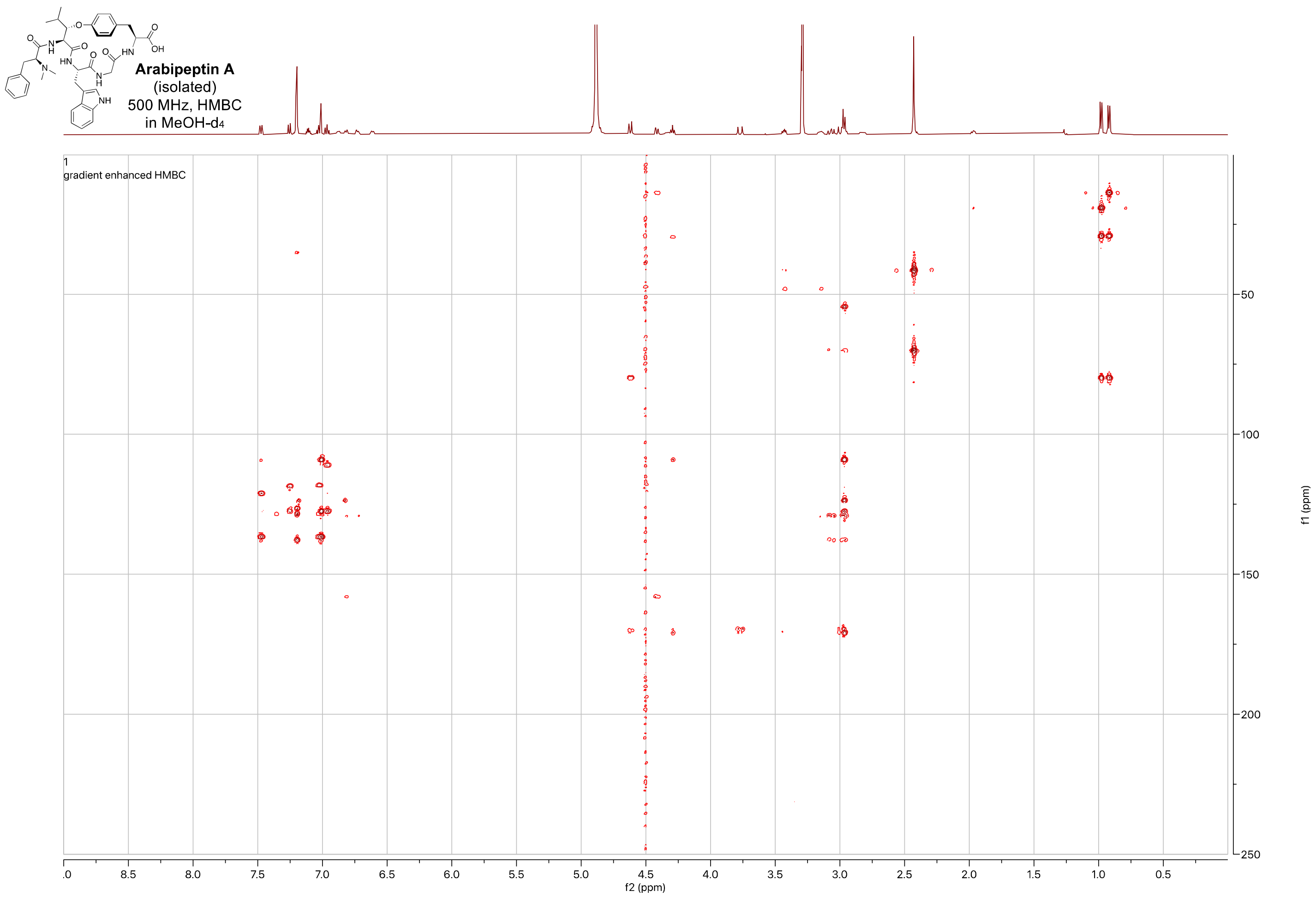


**Figure S35.** $^{1}$H-$^{13}$C-HMBC NMR spectrum of arabipeptin A in MeOH-$d_4$ (500 MHz).

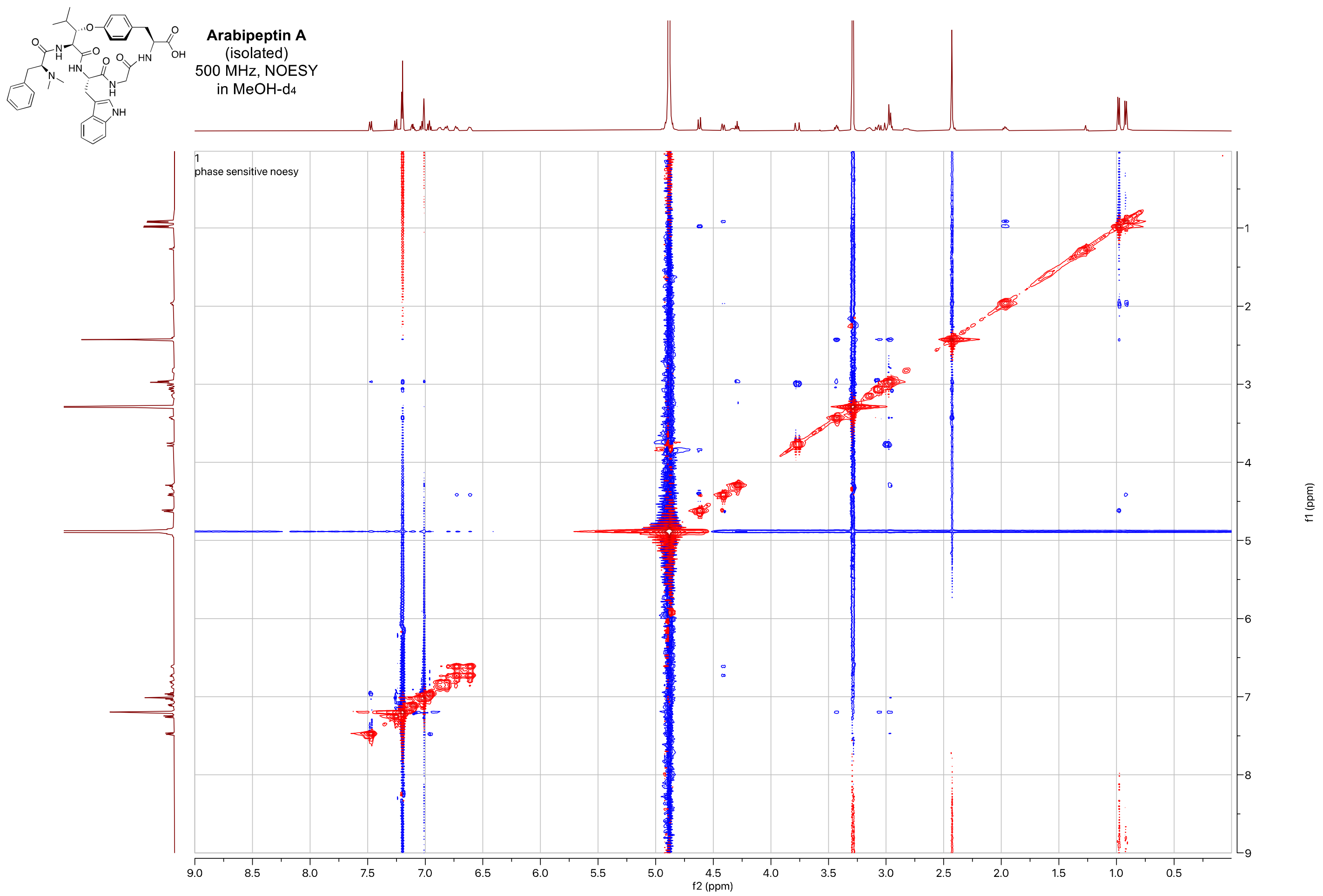


**Figure S36.** $^{1}H$-$^{1}H$-NOESY NMR spectrum of arabipeptin A in MeOH-$d_4$ (500 MHz).

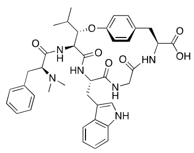

**Arabipeptin A**
(isolated)
500 MHz, TOCSY
in MeOH-d4

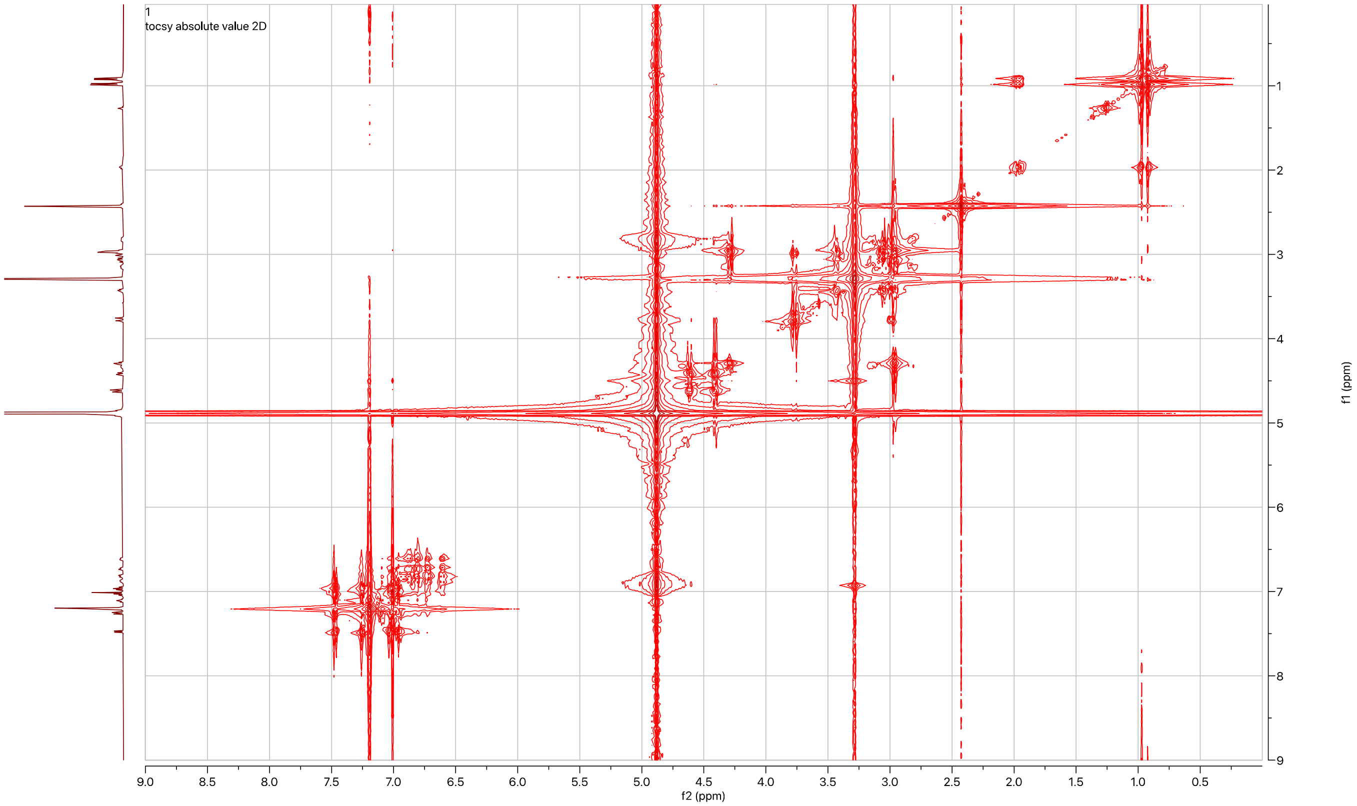


**Figure S37.** ¹H-¹H-TOCSY NMR spectrum of arabipeptin A in MeOH-$d_4$ (500 MHz).

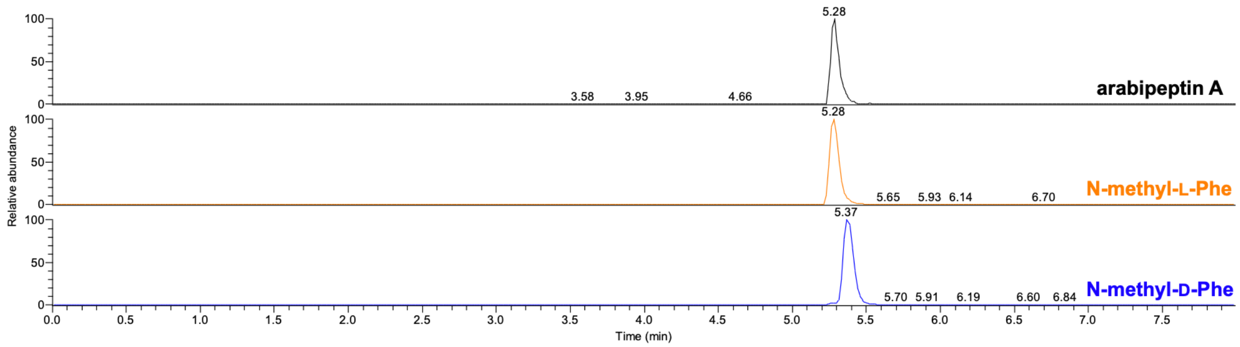
5.28
3.58 3.95 4.66
arabipeptin A
5.28
5.65 5.93 6.14 6.70
N-methyl-L-Phe
5.37
5.70 5.91 6.19 6.60 6.84
N-methyl-D-Phe
Relative abundance
Time (min)

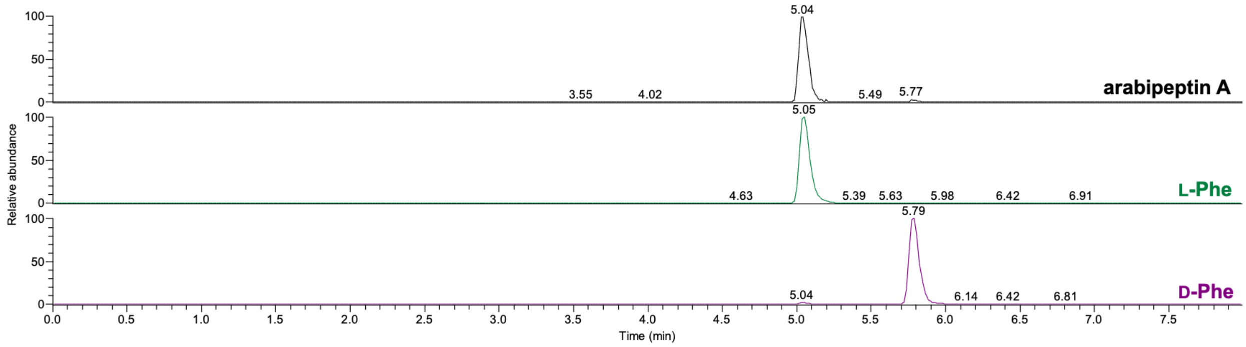
5.04
3.55 4.02 5.49 5.77
arabipeptin A
5.05
4.63 5.39 5.63 5.98 6.42 6.91
L-Phe
5.79
5.04 6.14 6.42 6.81
D-Phe
Relative abundance
Time (min)

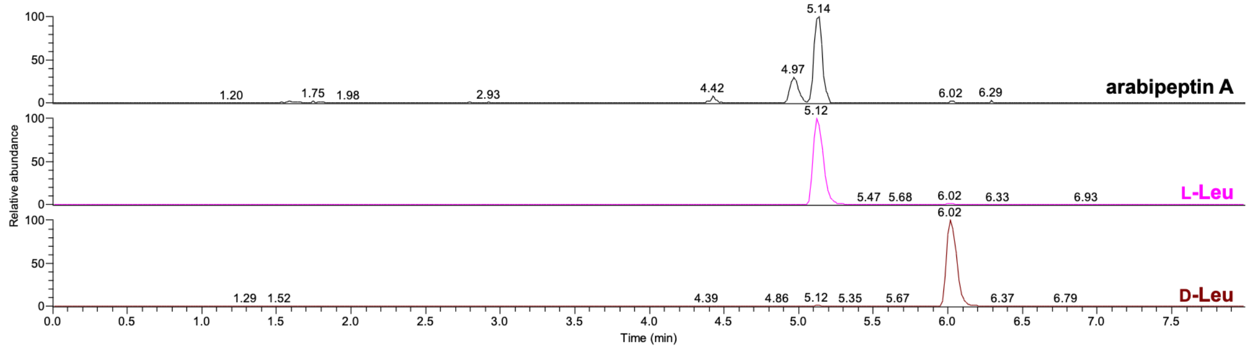
5.14
4.97
1.20 1.75 1.98 2.93 4.42 6.02 6.29
arabipeptin A
5.12
5.47 5.68 6.02 6.33 6.93
L-Leu
6.02
1.29 1.52 4.39 4.86 5.12 5.35 5.67 6.37 6.79
D-Leu
Relative abundance
Time (min)

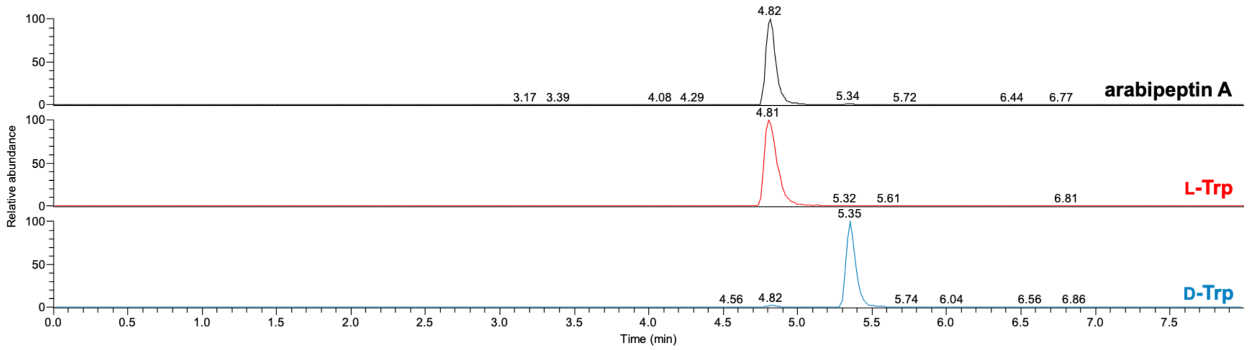
4.82
3.17 3.39 4.08 4.29 5.34 5.72 6.44 6.77
arabipeptin A
4.81
5.32 5.61 6.81
L-Trp
5.35
4.56 4.82 5.74 6.04 6.56 6.86
D-Trp
Relative abundance
Time (min)

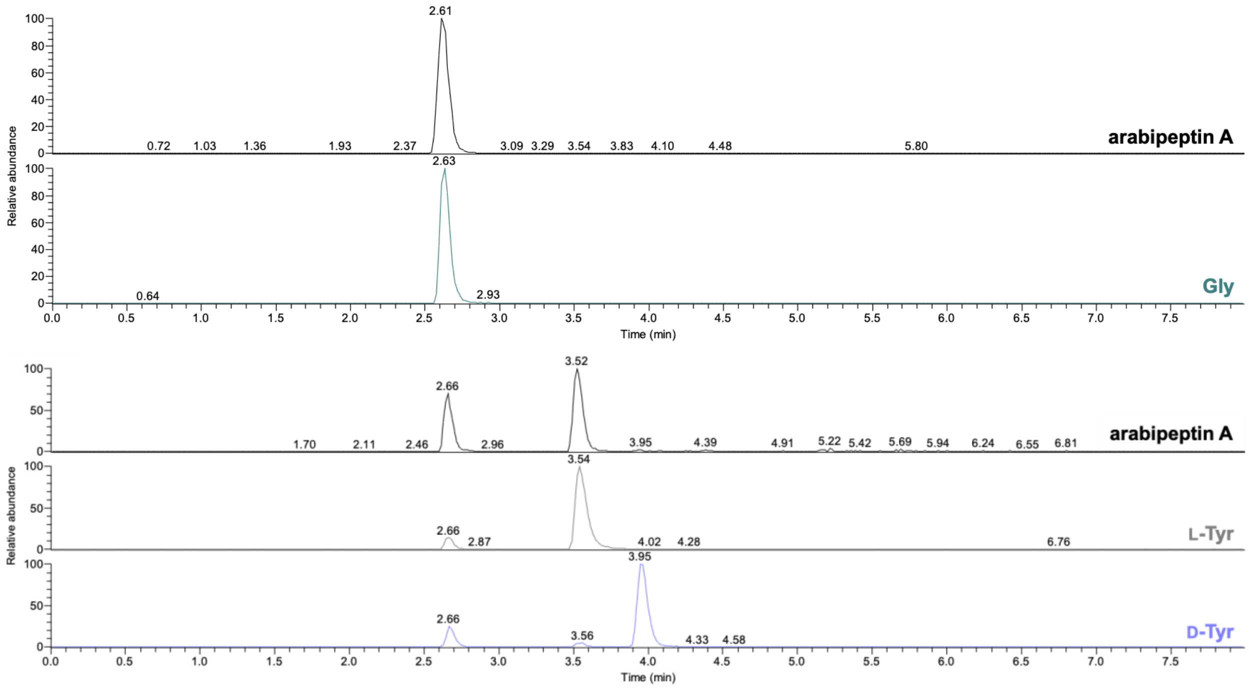


**Figure S38.** Marfey's analysis for arabipeptin A. Positive mode extracted ion chromatogram for hydrolyzed and derivatized arabipeptin A sample. Standard amino acids: *N*-methylphenylalanine, phenylalanine, leucine, tryptophan, glycine, and tyrosine ($[M+H]^+$ 432.1514, 418.1357, 384.1514, 457.1466, 328.0888, 434.1306 *m/z* respectively).

### Supplementary References


[1] M. Wang, J. J. Carver, V. V. Phelan, L. M. Sanchez, N. Garg, Y. Peng, D. D. Nguyen, J. Watrous, C. A. Kapono, T. Luzzatto-Knaan, C. Porto, A. Bouslimani, A. V. Melnik, M. J. Meehan, W.-T. Liu, M. Crüsemann, P. D. Boudreau, E. Esquenazi, M. Sandoval-Calderón, R. D. Kersten, L. A. Pace, R. A. Quinn, K. R. Duncan, C.-C. Hsu, D. J. Floros, R. G. Gavilan, K. Kleigrewe, T. Northen, R. J. Dutton, D. Parrot, E. E. Carlson, B. Aigle, C. F. Michelsen, L. Jelsbak, C. Sohlenkamp, P. Pevzner, A. Edlund, J. McLean, J. Piel, B. T. Murphy, L. Gerwick, C.-C. Liaw, Y.-L. Yang, H.-U. Humpf, M. Maansson, R. A. Keyzers, A. C. Sims, A. R. Johnson, A. M. Sidebottom, B. E. Sedio, A. Klitgaard, C. B. Larson, C. A. B. P, D. Torres-Mendoza, D. J. Gonzalez, D. B. Silva, L. M. Marques, D. P. Demarque, E. Pociute, E. C. O'Neill, E. Briand, E. J. N. Helfrich, E. A. Granatosky, E. Glukhov, F. Ryffel, H. Houson, H. Mohimani, J. J. Kharbush, Y. Zeng, J. A. Vorholt, K. L. Kurita, P. Charusanti, K. L. McPhail, K. F. Nielsen, L. Vuong, M. Elfeki, M. F. Traxler, N. Engene, N. Koyama, O. B. Vining, R. Baric, R. R. Silva, S. J. Mascuch, S. Tomasi, S. Jenkins, V. Macherla, T. Hoffman, V. Agarwal, P. G. Williams, J. Dai, R. Neupane, J. Gurr, A. M. C. Rodríguez, A. Lamsa, C. Zhang, K. Dorrestein, B. M. Duggan, J. Almaliti, P.-M. Allard, P. Phapale, L.-F. Nothias, T. Alexandrov, M. Litaudon, J.-L. Wolfender, J. E. Kyle, T. O. Metz, T. Peryea, D.-T. Nguyen, D. VanLeer, P. Shinn, A. Jadhav, R. Müller, K. M. Waters, W. Shi, X. Liu, L. Zhang, R. Knight, P. R. Jensen, B. O. Palsson, K. Pogliano, R. G. Linington, M. Gutiérrez, N. P. Lopes, W. H. Gerwick, B. S. Moore, P. C. Dorrestein, N. Bandeira, *Nat. Biotechnol.* **2016**, *34*, 828–837.

[8] "Pfam: Family: BURP (PF03181)," can be found under http://pfam.xfam.org/family/PF03181, **n.d.**

[9] I. Letunic, "phyloT : a phylogenetic tree generator," can be found under https://phylot.biobyte.de/, **n.d.**

[15] F. Teufel, J. J. Almagro Armenteros, A. R. Johansen, M. H. Gíslason, S. I. Pihl, K. D.

Tsirigos, O. Winther, S. Brunak, G. von Heijne, H. Nielsen, *Nat. Biotechnol.* **2022**, DOI 10.1038/s41587-021-01156-3.
